# Evolution of Origin Sequence and Recognition for Licensing of Eukaryotic DNA Replication

**DOI:** 10.64898/2026.03.10.710760

**Authors:** Jack Bauer, Narges Zali, Om Prakash Chouhan, Osama El Demerdash, Kaiser Loell, Matthew Tuttle, Justin B. Kinney, Leemor Joshua-Tor, Bruce Stillman

## Abstract

The large size of eukaryotic chromosomes necessitates that the initiation of DNA replication occurs at numerous origins of DNA replication. In *S. cerevisiae*, origins are highly DNA sequence-specific and are recognized by the Origin Recognition Complex (ORC). In contrast, most eukaryotes lack features in ORC subunits that contribute to DNA sequence-specific recognition, raising the question of how origins are identified. An analysis of origins in the genome of the distantly related budding yeast *Yarrowia lipolytica* identified considerable variability in origin sequence and structure. High-resolution structures demonstrate that *Y. lipolytica* origins are recognized by a combination of ORC and Cdc6 in a manner different from *S. cerevisiae*. The structure of *Yarrowia* ORC-Cdc6 bound to different origins shows considerable plasticity in protein-DNA interactions. We compare these yeast structures to the structure of human ORC-CDC6 bound to DNA. These studies reveal information on the evolution of origins and origin recognition.

*Nomenclature note:* There is a different nomenclature for proteins in yeast and human cells. For example, Cdc6 in yeasts is CDC6 in human cells.

## Introduction

The genome in eukaryotic cells is distributed over multiple large chromosomes that each contain numerous origins of DNA replication to ensure that all of the DNA is duplicated precisely once per cell division cycle ^1–4^. The location of origins in the genome is marked by the assembly of pre-Replicative Complexes (pre-RCs) prior to the initiation of actual DNA synthesis from each origin. Pre-RCs are assembled on all potential origins, usually following exit from the previous mitosis or during G1-phase ^5–11^. The best characterized system for understanding the biochemistry of complete DNA replication, including pre-RC assembly, derives from studies of the budding yeast *S. cerevisiae* ^5,7,12–17^.

In *S. cerevisiae*, pre-RCs are assembled by the binding of the Origin Recognition Complex (ORC) to specific DNA sequences called Autonomously Replicating Sequences (ARSs) that determine the location of ∼500 origins in the 12.2 Mb genome ^7,18,19^. ORC, a six-subunit ATPase, binds to and bends the origin DNA and then recruits the Cdc6 ATPase. Together, these proteins load two copies of the Mcm2-7 hexamer that are chaperoned by the Cdt1 protein to form the MCM double hexamer (DH) ^5,6,16,17,20^. The MCM DH is destined to separate into two divergent replicative helicases called the CMG (Cdc45-Mcm2-7-GINS), which encompasses Cdc45, the Mcm-2-7 hexamer, and the four-subunit GINS complex ^21^. The assembly of the helicase and subsequent replication of DNA occurs following activation of the pre-RC by the S-phase Cyclin-Dependent Kinases (CDKs) and the Cdc7-Dbf4 kinase (DDK) ^16^.

The origins of DNA replication in *S. cerevisiae* consist of multiple essential or important DNA elements. The A and B1 DNA elements are recognized by ORC, whereas the B2 element is a weak ORC binding site that is in an inverted orientation and of variable distance from the A and B1 elements ^12,22–26^. Depending on this distance, two modes of assembly of pre-RCs can occur, one requiring only a single ORC and the other involving two separate ORCs ^12,17,26,27^. Since the genome of *S. cerevisiae* is relatively compact with little repeat sequences and has very short intergenic DNA regions, origins of DNA replication have most likely evolved to be highly DNA sequence-specific and located in non-transcribed regions of the genome so that the initiation of DNA replication does not conflict with gene transcription ^28^. As a consequence, most *S. cerevisiae* origins are located within the short intergenic DNA sequences. ^18,29^ Origin specificity in *S. cerevisiae* occurs in part by the interaction of an α-helix in the Orc4 subunit that inserts into a major groove in the origin DNA, a loop in the Orc2 subunit that inserts into a minor groove in the origin DNA, and a lysine-rich region in the intrinsically disordered domain of Orc1 that also binds a DNA minor groove ^28,30–32^.

A small clade of budding yeasts that are evolutionarily related to *S. cerevisiae*, including *Kluyveromyces lactis* and *Lachancea kluyveri* have ARSs and origins that are related in sequence to the *S. cerevisiae* origins ^33,34^. The Orc2 loop and Orc4 α-helix in these species are conserved ^35^. In contrast, all other eukaryotes, including other budding yeasts and fungi, and all animals and plants have either lost completely or truncated these origins recognition elements ^1,35^. In some budding yeasts such as *Candida albicans* and *Pichia pastoris (now called Komagataella phaffii)*, ARS sequences have been characterized and are very different from the *S. cerevisiae* clade of ARS sequences, ^36,37^ but the manner in which the proteins interact with them has not been addressed. Other yeasts, such as the fission yeast *S. pombe*, have gained an unusual A/T-rich hook domain in Orc4 that binds to the A.T-rich origins of DNA replication, but this mode of origin recognition is not common. Similar to pre-RC assembly using purified *S. cerevisiae* proteins, ^38,39^ pre-RC assembly has been reconstituted with purified human proteins, demonstrating both a one-ORC and a two-ORC mechanism of MCM DH loading onto non-specific DNA ^17,40–42^. While this may suggest that ORC can determine the location of origins of DNA replication in human cells, a meta-analysis of multiple studies that mapped ORC and MCM binding sites in the human genome showed a very poor correlation with the location of origins of DNA replication ^43^. This may be due to technical reasons, but there remains the matter of how origin recognition, and hence the specification of origin location in most eukaryotes occurs.

In animal cells, such as *C. elegans*, *Drosophila* and mammalian cells, including human cells, origins of DNA replication have been mapped and they correlate with genomic features such as histone modifications, higher-order chromosome structure, and in many cases transcription start sites ^1–3,44,45^. For example, in *C. elegans*, the efficiency of origins of DNA replication is associated with histone H3-lysine-4-dimetylation (H3K4me2) and histone H3-lysine27-acetylation (H3K27Ac) ^46^. In *Drosophila* and human cells, DNA topology and certain chromatin features mark replication origins, and they are commonly associated with regions that contain nearby predicted G4-quartet DNA structures ^47–51^. In human cells, origins of DNA replication are located both at specific loci such as open chromatin regions, but initiation of DNA replication can also occur in a distributed fashion, where stochastic origin firing takes place in chromosome replication initiation domains ^44,45,52–57^. How these specific and distributed origins are specified is not known, but speculation about epigenetic marking of the initiation of DNA replication is common ^4,44,58,59^.

In a study of the mechanism of origin specificity in *S. cerevisiae*, we noticed that the Orc4 α-helix and Orc2 loop that provided DNA sequence-specific interactions with origins were only conserved in the small clade of *S. cerevisiae*-related budding yeasts, whereas many other budding yeasts and all other eukaryotes, including plants and animals, including human ORC, lacked these conserved features ^28^. In this report, we first determined the structure of human ORC-CDC6 bound to a G/C rich DNA. Though we observed DNA bending seen in all ORC-DNA-CDC6 (ODC) complexes to date as well as a surprising minor groove contact by HsORC2, we reasoned that a stronger evolutionary perspective would aid in understanding origin specification. We therefore began studies on *Yarrowia lipolytica,* which lacked the origin-recognition features seen in *S. cerevisiae*. *Y. lipolytica* is a non-conventional, oleaginous yeast that is widely used in biotechnology whose last common ancestor with *S. cerevisiae* existed ∼300 million years ago ^60^. Unlike *S. cerevisiae*, *Yarrowia* is heterothallic, having two separate mating types, MatA and MatB. Previous studies identified a few origins that are located near centromeres, probably because, unlike *S. cerevisiae* ARSs, propagation of extra-chromosomal plasmids in *Yarrowia* requires both a centromere and an origin sequence on the plasmid ^61–64^. To study DNA replication in *Yarrowia* more thoroughly, we mapped the location of origins in all six chromosomes, demonstrating a genome organization of replication timing domains reminiscent of those in the genomes of animal cells, including human cells. Genetic analysis of two of these origins, one a centromere-associated origin and the other an origin on a chromosome arm, uncovered a short ∼30 bp essential region, and massive parallel mutational analysis revealed that *Y. lipolytica* origins of DNA replication are heterogeneous. Structural studies of ORC and Cdc6 bound to the two different origin DNA sequences demonstrated that, unlike *S. cerevisiae*, *Y. lipolytica* origin recognition required both ORC and Cdc6 for base-specific interactions and hence origin recognition, with some protein-DNA interactions varied between the two origins. The results show a surprising plasticity in origin sequences, structure, and recognition in different eukaryotes. We discuss the evolution of origin recognition and specificity.

## Results

### Cryo-EM structure of the human ORC–DNA–CDC6 complex

The human ORC and CDC6 bound to DNA (HsODC) was reconstituted by combining HsORC1– 5, HsCDC6, and DNA *in vitro*. The sequence specificity of HsORC and origins of DNA replication in human cells remains unknown, but origins are known to be G.C-rich ^65^. Therefore, we selected a defined DNA 60 base-pair fragment with 70% G/C content for complex assembly after confirming ORC binding through biochemical assays. The 2.6 Å-resolution cryoEM map (**Figures 1A, S1A and S1B)** enabled the unambiguous placement of all five ORC subunits, HsCDC6, and the bound DNA. The N-terminal regions of HsORC1 (amino acids 1-465), HsORC2 (aa 1-165), and HsCDC6 (aa 1-151) proteins, which consist of intrinsically disordered regions (IDRs) are disordered and therefore not visible. Of the 60-bp DNA used, 29-bp were built with confidence, and density corresponding to four ATP analogs was clearly resolved at the conserved nucleotide-binding sites within ORC. Local resolution analysis demonstrated that the AAA+ (ATPases Associated with diverse Activities) core of the complex, comprising RecA-like domains from each ORC subunit and HsCDC6, was better resolved than the central DNA, which exhibited flexibility and correspondingly lower resolution.

**Figure 1.**
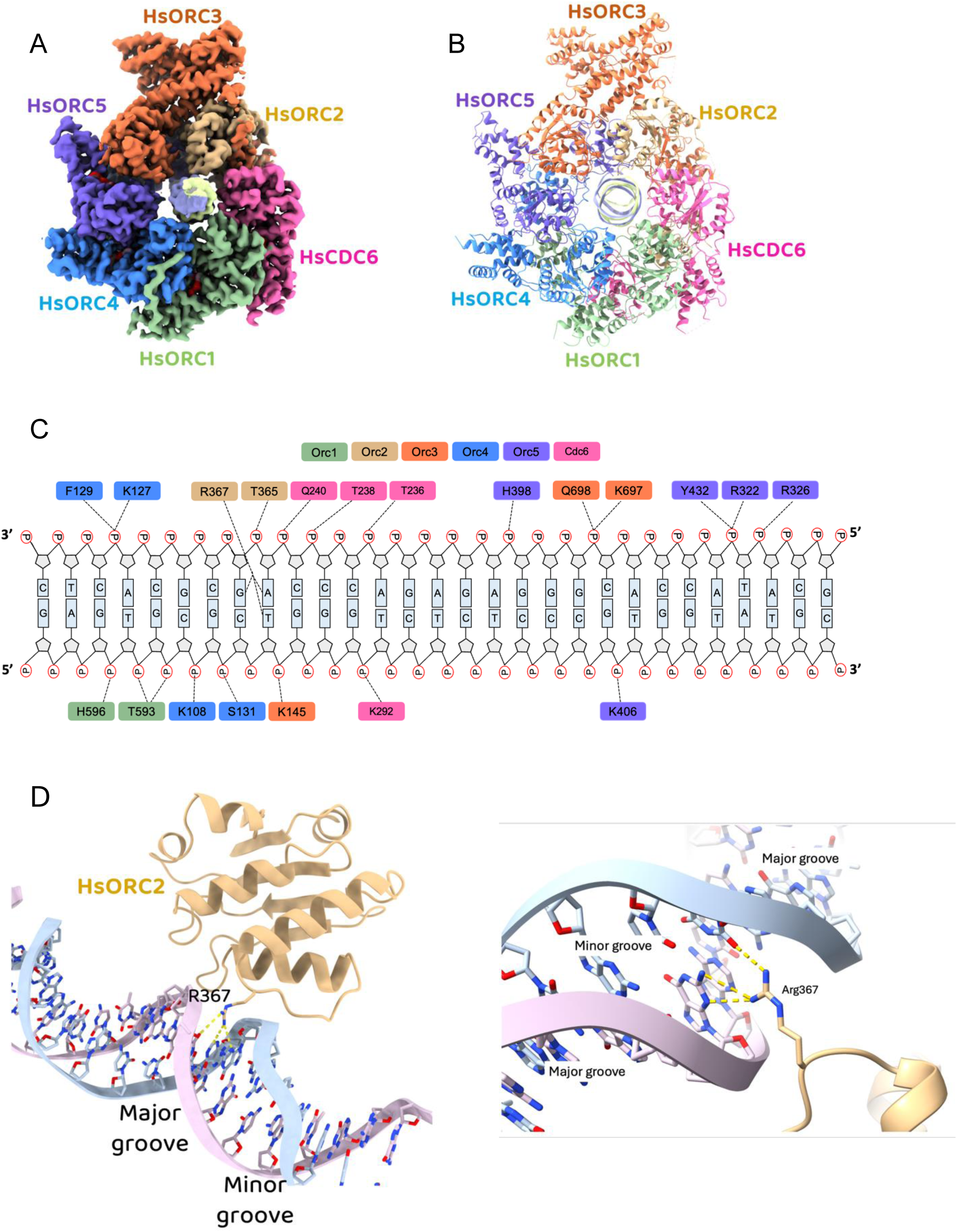
Structure of the HsORC–Cdc6–DNA complex. **(A)** Cryo-EM 3D map of the HsODC complex bound to a 60-bp DNA duplex, with each protein subunit shown in a distinct color. **(B)** Cartoon representation of the atomic model fitted into the cryo-EM map, highlighting all protein subunits of HsORC, Cdc6, and the DNA molecule. This view emphasizes the arrangement of the proteins around the central DNA-binding channel. **(C)** Interactions between HsORC subunits and DNA, shown at the amino acid level. Interacting residues are colored as above, providing a clear visualization of how each subunit contributes to DNA engagement. **(D)** Close-up view of residue R367 from the HsORC2 subunit, which establishes distinct contacts with DNA nucleotides, specifically thymine from chain H and guanine from chain I.

#### Overall architecture

HsODC adopts a closed-ring conformation, in which all six protein components encircle a centrally located DNA duplex. The complex has a two-tiered layered appearance, with the AAA+ domains forming one layer and the winged-helix domains (WHD) forming a second layer. Each WHD from one subunit sits atop the AAA+ domain of a neighboring subunit (**Figure 1B, Suppl. Video 1**). The HsODC structure is very similar to HsORC1-5 bound to DNA that copurified from the expression host cells (PDB ID: 7JPS) ^66^, with HsCDC6 closing the ring around the DNA. The overall RMSD between the two structures is 4.0 Å over backbone atoms, without the DNA and HsCDC6, though the RMSD between individual subunits is considerably lower (between 0.6 and 1.5 Å). The HsCDC6 AAA+ domain is nestled between the AAA+ domains of HsORC1 and HsORC2, and its WHD sits on top of the AAA+ domain of HsORC1. The higher resolution of this structure, compared to ORC-DNA alone^66,67^, brings additional features into view. The HsORC2 WHD was not visible in the HsORC structure but there is clear density for this domain in the HsODC structure, which is situated above the RecA domain of HsCDC6. There is a slight widening of ∼4.2Å at the interface between HsORC1 and HsORC2, creating sufficient space to accommodate HsCDC6. This local broadening allows the RecA domain of HsCDC6 to insert into the gap, where it establishes contacts with both HsORC1 and HsORC2. An additional ATP-binding site is formed between HsCDC6 and HsORC1, as is the case for ScODC and DmODC ^68–70^.

#### DNA bending

Previous studies in *S. cerevisiae* have shown that DNA bending mediated by ORC is important for origin licensing and subsequent MCM loading ^17,25,31,71^. In *S. cerevisiae*, replication origins contain an A/T-rich ARS consensus sequence (ACS) and ORC bends DNA downstream of the ACS site by 40–55° relative to the axis of ORC’s central DNA-binding channel. This bending is primarily driven by a basic amino acid patch within the Orc5 subunit (ORC5-BP), which contains a long loop (amino acids 350 - 370) enriched with glycine and alanine residues, which confer flexibility, along with basic amino acids that extend into the DNA minor groove. In our HsODC structure, we observe clear density for only two HsORC5 arginine residues that make limited contacts with the DNA backbone and are at the very beginning of this highly flexible loop (**Figure 1C, Figure S1C**). The rest of the loop is disordered but it contains additional basic resides (**Figure S1C**). These interactions might be insufficient to achieve the degree of bending seen in *S. cerevisiae* ODC, resulting in a more modest bend of ∼20° relative to the axis of the central DNA-binding channel. This difference suggests that the mechanism of DNA engagement and bending by human ODC may either be inherently less pronounced than in yeast, potentially reflecting species-specific adaptations in origin recognition, or that reduced bending may reflect the fact that the DNA used is not a *bona fide* origin.

#### ORC-CDC6 engages DNA through backbone interactions with all protein subunits

Detailed examination of the cryo-EM structure revealed that the five subunits of human ORC (HsORC1– 5) and HsCDC6 engage directly with the DNA duplex (**Figure 1C and S1D**). The majority of these interactions are mediated through the RecA-like domains of the ORC subunits. Notably, only the WHDs of HsORC3 and HsORC5 make direct contacts with the DNA, rather than all of them in the case of ScODC, suggesting a specialized role for these WHDs in stabilizing DNA binding within the closed-ring architecture during the assembly of the pre-Replicative Complex. The high-resolution of the cryo-EM map enabled detailed mapping of the protein–DNA interface within the ODC. HsORC1 engages the DNA through residues T593 and H596, positioned to interact with the phosphate backbone near the DNA entry point of the complex. HsORC2 contacts DNA via T365 and R367. HsORC3 contributes a cluster of residues—R641, K697, and Q698 that interact with the DNA backbone. Like all other ORC subunits, HsORC4 also exhibits a DNA-binding interface, with residues K127, F129, S131, and T391 forming a broad contact surface that likely plays a role in anchoring the DNA and stabilizing the complex (**Figure 1C**).

A particularly notable feature is the DNA-binding mode of HsORC5, which utilizes its WHD to engage the DNA through a cluster of basic residues—R322, R326, and Y432. These residues form a distinct basic patch that establishes strong electrostatic interactions with the phosphate backbone of the DNA. This interaction likely contributes to the bending of the DNA toward the protein surface. Such localized bending may play a critical role in the structural remodeling of the origin DNA, enabling the recruitment and loading of downstream replication factors, such as the MCM2–7 helicase, as shown for ScODC ^71^. HsCDC6 also participates in DNA binding, contributing contacts via residues T236, T238, and Q240 to the phosphate backbone of the DNA. These protein-DNA interactions across all six subunits establish a high-affinity DNA-binding surface that facilitates origin recognition and pre-Replicative Complex assembly.

#### A direct interaction with a DNA base

Since sequence-specific origins have yet to be identified in metazoans, we initially expected to observe only non-specific interactions with the DNA backbone. However, residue R367 from HsORC2 extends into the minor groove of the DNA and interacts with a nitrogen base edge (**Figure 1D and S1E**). Although the cryo-EM density for the full guanidinium group of the arginine side chain is incomplete, the visible portion is sufficient to model its orientation and infer that it is positioned to form hydrogen bonds with a guanine and a thymine on the adjacent base pair (**Figure S1E**). This type of contact is noteworthy, as minor groove base interactions can contribute to sequence-preferential recognition, even in proteins or DNA that are not strictly sequence-specific. The neighboring residue, T365, binds the DNA phosphate backbone and likely stabilizes the orientation of R367, effectively “locking” it into position. Intriguingly, R367 is on a loop that occupies the same position that the Orc2 DNA-binding loop does in *S. cerevisiae* and R367 itself is in a similar position to the *S. cerevisiae* Orc2 DNA-binding residue, W396 **(Figure S1E-G)**. The R367 residue is conserved in most vertebrates (**Figure S1G**). Although the DNA used in this study is not derived from a *bona fide* origin sequence, the structural arrangement of T365 and R367 suggests a potential mechanism by which HsORC2 could engage in limited DNA sequence-dependent recognition. Comparison with available human ORC structures, in the OCCM (PDBIDs 8S0E, 8RWV) and the ORC1-5-DNA complex (PDBID 7JPS), shows that the ORC2 WHD fold is conserved in all these structures. The side chain of R367 in the OCCM structures is modeled away from the DNA and does not contact it, but the density for this region is either completely absent or of low resolution. Notably, the GC content in these structures is lower (55 and 29%) than in the structure presented here. However, R367 in the ORC1-5-DNA structure, the DNA conformation is different than in the OCCM, and the side chain is modeled in proximity to the DNA. The DNA in that case was copurified with the protein and is a mixture of sequences.

#### Nucleotide binding sites

The HsODC complex was assembled in the presence of the non-hydrolyzable ATP analog AMPPNP. As in most AAA+ ATPases, including ORC, nucleotide binding occurs at the interface between adjacent subunits, where conserved motifs from neighboring proteins contribute to the formation of the nucleotide-binding pocket ^72^. Upon addition of CDC6 to the human ORC complex, we observed additional density corresponding to a nucleotide at the interface between CDC6 and ORC1, with HsORC1 R670 serving as the arginine finger, and HsCDC6 residues R388 and K208 coordinate the β- and γ-phosphates of ATP. The cryo-EM densities for all four-nucleotide binding regions were sufficiently well-resolved to allow modeling of a magnesium ion coordinated near the ATP analog (**Figure S1H**).

### Genome-wide identification of origins of DNA replication in *Yarrowia lipolytica*

To investigate the evolution of origin recognition, we identified origins of DNA replication in *Yarrowia lipolytica* (**Figure 2A**). A strain of *Y. lipolytica* was constructed that expressed the Herpes Simplex Virus thymidine kinase (HSVTK) and the human Equilibrative Nucleoside Transporter 1 (ENT1) proteins to enable incorporation of the thymidine analog 5-ethynyl-2′-deoxyuridine (EdU). The temporal dynamics of DNA replication in *Y. lipolytica* was investigated by EdU-labeling following synchronization of cells by nutrient starvation and release into the cell division cycle by re-feeding. EdU-positive cells were visualized using fluorescence microscopy and quantified alongside budding index as an indicator of S-phase entry (**Figure S2A and S2B**). The peak of S phase under these conditions was 60-75 minutes post release, however, when the DNA synthesis inhibitor hydroxyurea (HU) was added, replication progression was slower, but more synchronous (**Figures 2A and S2C**).

**Figure 2:**
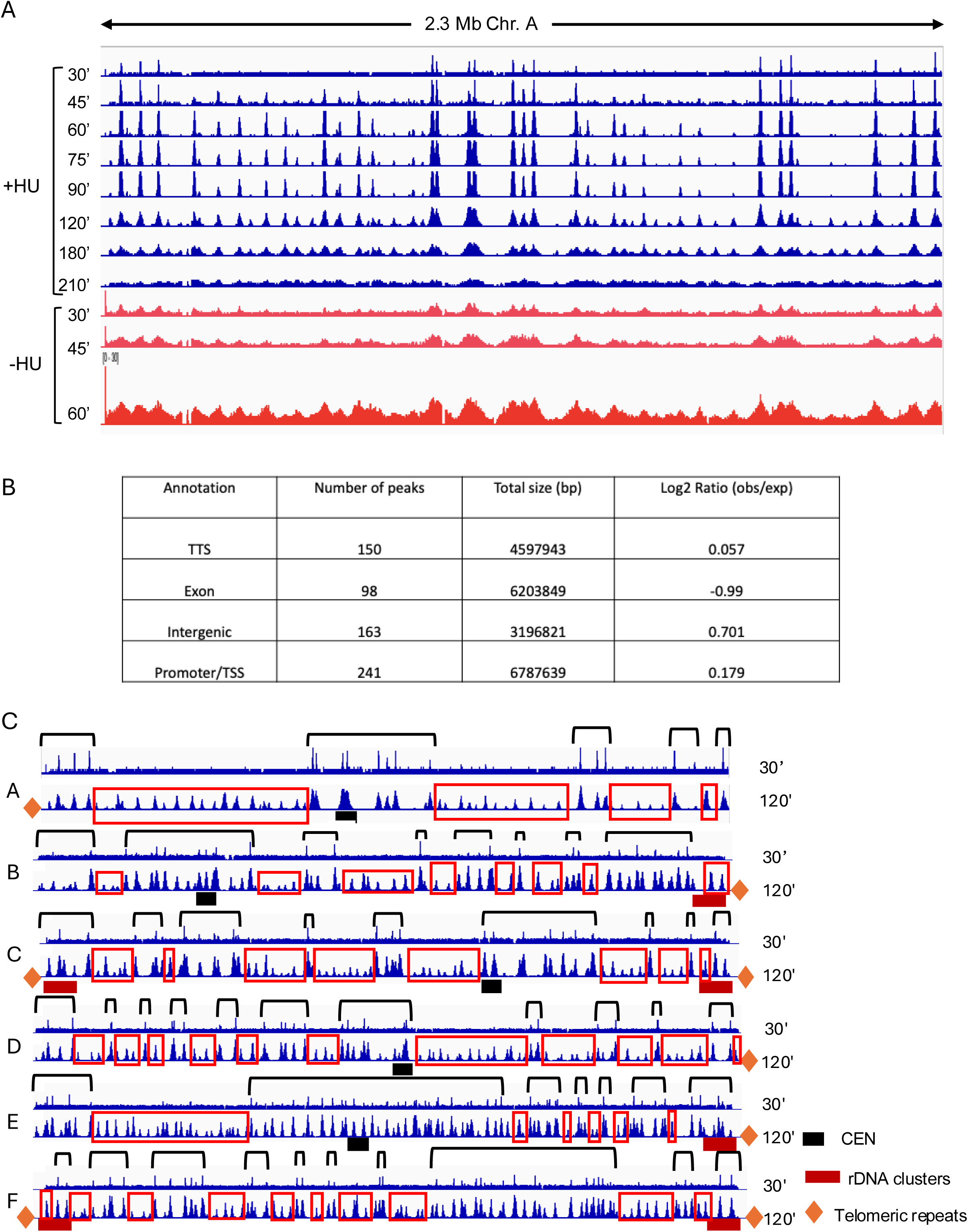
Genome-wide profiling and genomic context of replication origins in *Yarrowia lipolytica*. **(A)** Temporal mapping of replication origin activity throughout *Yarrowia lipolytica* genome using EdU-seq. EdU-seq signal tracks across a 2.3 Mb region of Chromosome A following release from starvation into S phase in the presence (blue) or absence (red) of 5 mM hydroxyurea (HU). Samples were collected at the indicated time points. HU-treated cells show temporally resolved activation of replication origins, while untreated cells exhibit more extensive EdU incorporation. **(B)** Enrichment analysis of EdU-seq peaks relative to genomic annotations. Intergenic and promoter/TSS regions are enriched for replication origins, while exonic regions are depleted. Log2(observed/expected) values indicate the degree of enrichment or depletion across different genomic features. **(C)** Genome-wide origin firing maps across all six *Y. lipolytica* chromosomes at 30 and 120 minutes post-release in the presence of HU. Each track shows EdU-seq signal along individual chromosomes (A–F), with peaks corresponding to active replication origins. Regions highlighted in black boxes represent early-firing origins (30’), while red boxes highlight later-firing origins (120’). Centromeres (black boxes), rDNA clusters (red boxes), and telomeric regions (orange diamonds) are annotated for reference.

The whole genome landscape of DNA replication in *Y. lipolytica* was determined by continuous labeling of DNA replication with EdU in the presence and absence of HU and harvesting the cells at different times post release. Compared to *S. cerevisiae*, where 200mM HU inhibits DNA replication and cell viability, *Y. lipolytica* is very sensitive to HU. While in the presence of 5mM HU, cells can still progress through the cell cycle, higher levels of HU inhibit cell proliferation and cell viability. Labeled DNA was detected by sequencing and mapped to a high-quality genome assembly ^73^. A total of 556 replication origin peaks were identified with much sharper peaks when HU was added due to checkpoint inhibition of replication fork progression, allowing better definition of the temporal activation of origins during S phase (**Figure 2A**). Importantly, HU did not alter origin location. Early replicating origins were first detected at 30 minutes post release and the latest active origins appeared at 120 minutes. Of the 556 replication origins identified genome-wide, 346 were classified as early-firing under HU treatment. Most origins were in intergeneic regions of the genome, with some at transcription start sites (**Figure 2B).**

Importantly, the spatial distribution of timing of activation of replication origins across the chromosomes was not distributed uniformly, but instead formed discrete 150–300 kb clusters of origins that were activated at the same time (**Figure 2C**), closely resembling the replication timing domains described in the chromosomes of higher eukaryotes ^74,75^.

### Genetic analysis of two *Yarrowia* origins of DNA replication

The 556 origins were annotated by referring to the chromosome (A through F) and counting each origin from the left telomere to the right telomere. For example, *OriA-006* is a newly discovered origin that is toward the left telomere on chromosome A. To genetically characterize the DNA sequences under the EdU-seq peaks, we tested two origins using ARS assays: the previously characterized, centromere-associated *OriC-061* (previously called *ARS18* or *Ori3018*) ^61,62^ and *OriA-006*. Unlike ARSs in *S. cerevisiae* in which an origin can support high frequency transformation (HFT) and plasmid stability, *Y. lipolytica* requires both a centromere and an origin of DNA replication be present on the mini-chromosome (**Figure S3A)** ^61^. Both *OriA-006* and *OriC-061* supported robust plasmid replication when cloned into an *Ori*⁻/CEN⁺ plasmid backbone, including equivalent ARS activity (both HFT and plasmid stability) when placed in either orientation or at variable distances from the centromere (**Figure S3B**). In contrast, two 600 bp fragments not associated with EdU-seq peaks, one derived from a coding region on Chromosome D and the other from a non-coding intergenic region on Chromosome E, could not support ARS activity (**Figure S3C**). These findings demonstrate that not all genomic sequences can function as replication origins, highlighting the requirement for specific DNA elements. Furthermore, they reinforce the reliability of EdU-seq in identifying biologically active replication origins and provide a foundation for dissecting the sequence and structural features critical for origin function in *Y. lipolytica*.

Linker scan mutagenesis ^76^ overcomes concerns that deletion mutations can alter the spacing of essential DNA sequences and was previously employed to dissect the structure of *S. cerevisiae* origins ^22–24^. We therefore used linker scanning to analyze the sequence requirements of *Y. lipolytica* origins within the 600 bp *OriC-061* and *OriA-006* fragments (**Figure 3A and 3B**). Each mutant was screened for high-frequency transformation (HFT) and plasmid stability. In *OriC-061*, 45 linker mutants were tested and substitutions at positions 2–6 severely reduced or abolished HFT and plasmid stability, identifying this region as critical for replication initiation (**Figure 3A**). Mutations outside this core retained wild-type–like stability (∼41%), suggesting they are dispensable for origin function. For *OriA-006*, linker insertions between positions 25–29 reduced or eliminated both transformation efficiency and plasmid maintenance (**Figure 3B**). Together, these results demonstrated that replication origin activity in *Y. lipolytica* is dependent on a short ∼30 bp sequence of DNA. These sequences are sufficient for origin activity since for each origin, a 50 bp fragment supported ARS activity in the presence of a CEN (**Figure S3D**).

**Figure 3.**
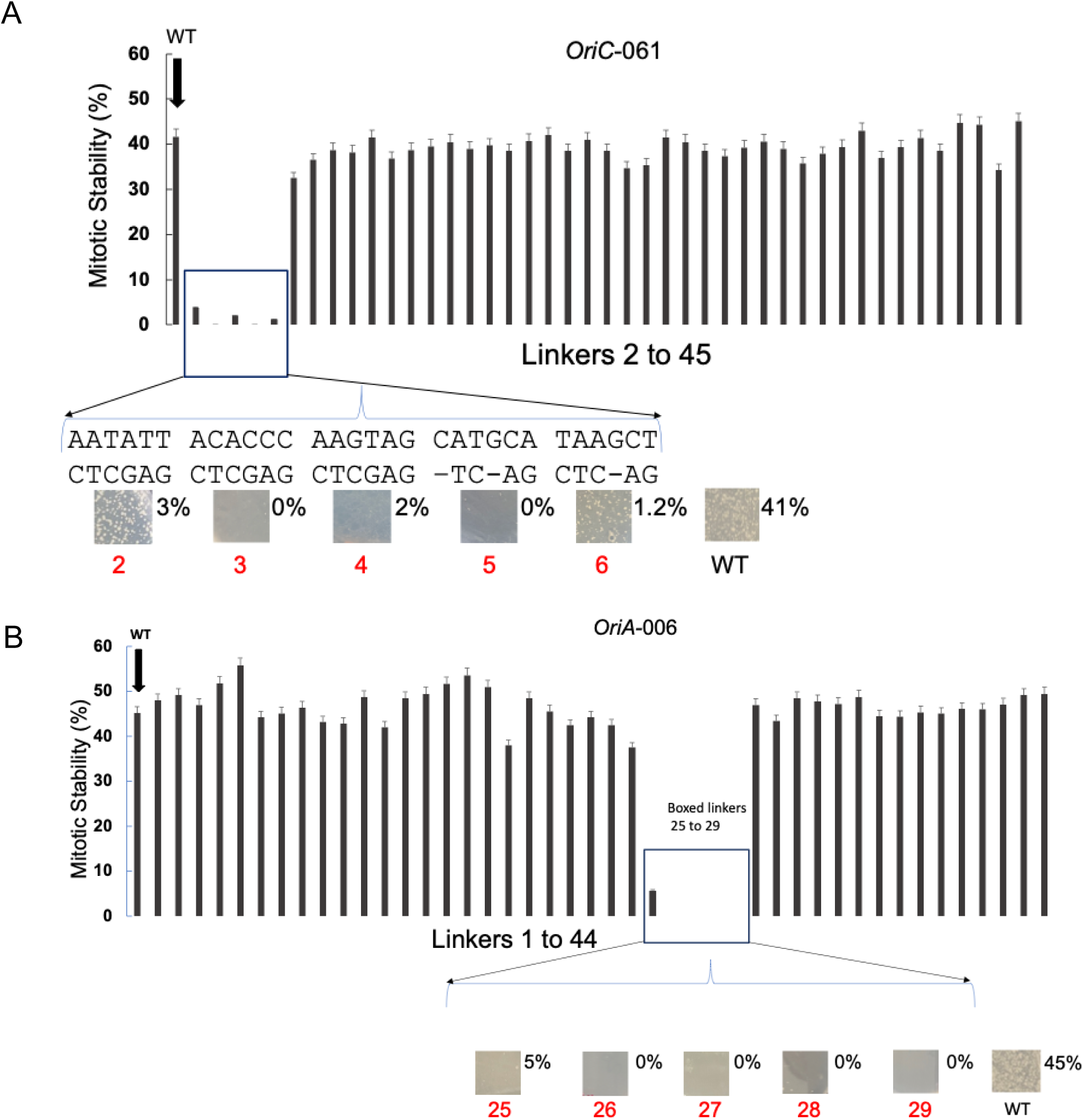
Linker-scanning mutagenesis reveals essential regions for origin activity in *Y. lipolytica OriC-061* and *OriA-006*. **(A)** Mitotic stability assay of *OriC-061* mutants carrying XhoI linker substitutions (CTCGAG) at positions 2–45. Images show colony formation following plasmid transformation and selection and percentage (% URA+ retention) for different linker mutations. Wild-type (WT) *OriC-061* shows ∼41% mitotic stability, while substitutions in linkers 2 to 6 drastically reduce origin activity to ≤3%. A zoomed-in view highlights the six critical linker mutants with corresponding colonies on the plate after the initial. transformation, with the percentage plasmid stability values shown. **(B)** Mitotic stability assay of *OriA-006* mutants with linker substitutions at positions 1–44. WT *OriA-006* exhibits ∼45% mitotic stability. A cluster of mutations (linkers 25–29) results in complete loss of origin activity (0–5% stability), identifying a key functional region. Sequences and representative colony phenotypes for these five mutants are shown below the graph.

### Structures of Yarrowia ORC-Cdc6 Bound to Different Origin DNAs

#### Origin DNA sequences are required for Cdc6 DNA binding

A Cdc6 DNA-binding assay using size exclusion chromatography (SEC) was developed to test whether specific DNA sequences were required for Cdc6 to co-elute with ORC. A 54 bp fragment of *OriC-061* or a scrambled version of the DNA with the same G/C content was used. Wild-type *OriC-061* DNA and some Cdc6 co-eluted with ORC (**Figure 4A top, fractions 3 and 4**), however, when the scrambled DNA was used, ORC binding to DNA was greatly reduced and Cdc6 no longer co-eluted with ORC (**Figure 4A, bottom**). This was validated by mass photometry (**Figure S4A**).

**Figure 4.**
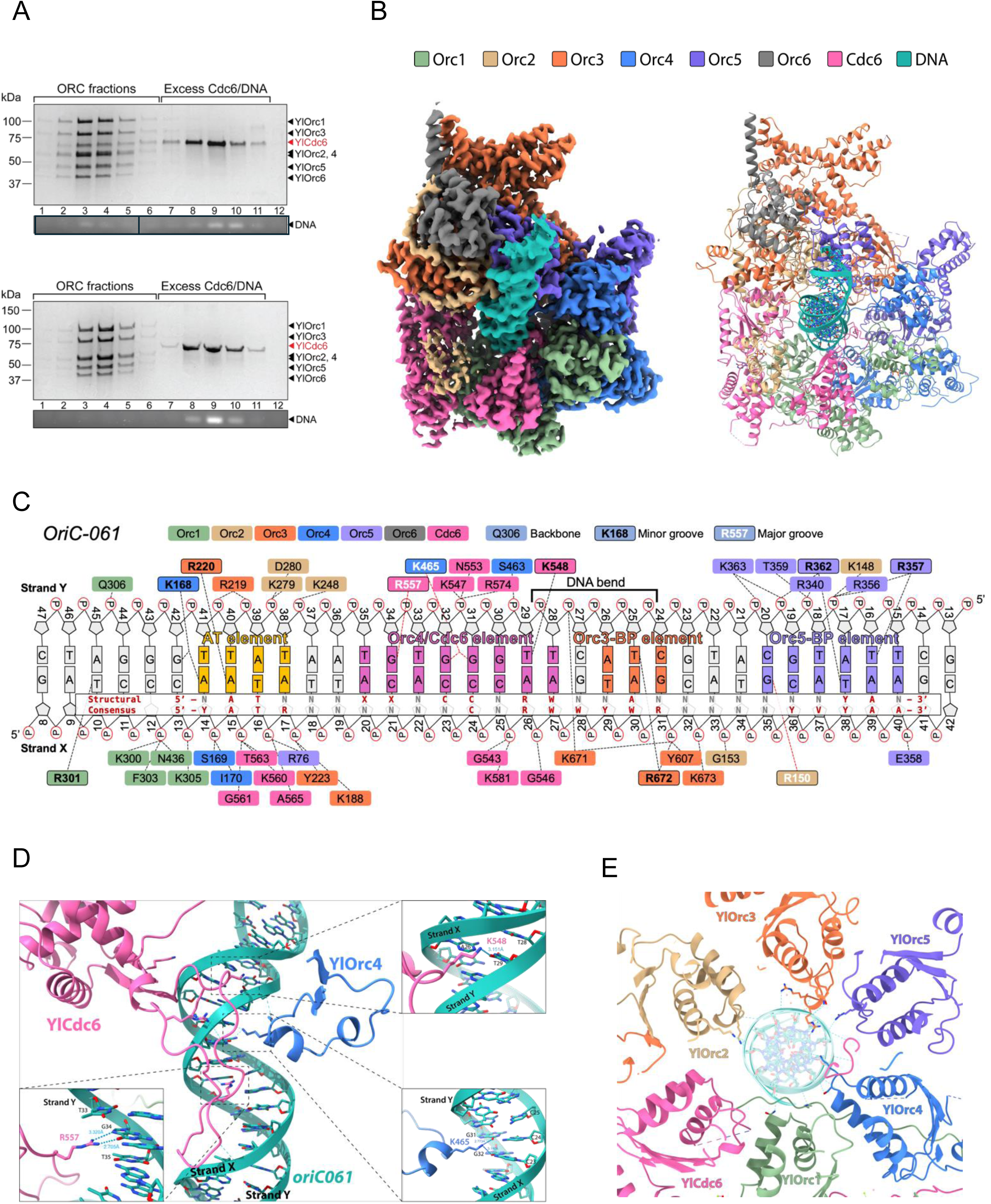
YlORC and YlCdc6 coordinate to bind origins specifically. **(A)** SDS-PAGE and agarose gel electrophoresis of gel filtration fractions from samples containing YlOrc1-6[SST-YlOrc1], YlCdc6, and a 54-bp DNA oligonucleotide derived from *OriC-061* (top panel) and a scrambled sequence thereof (bottom panel). The agarose gel in the top panel was spliced together as shown in the black boxes, but represents DNA from the same experiment. **(B)** (*left*) The 2.7 Å resolution unsharpened map of the YlORC-DNA^54bp*OriC-061*^–YlCdc6 complex and (*right*) the ribbon representation of the derived structure. **(C)** A diagram of the visible protein-DNA interactions seen in the YlORC-DNA^54bp*OriC-061*^–YlCdc6 structure. N indicates any nucleotide; W denotes an A or T; XX denotes a C in either of these positions; R indicates a purine; Y indicates a pyrimidine. **(D)** A ribbon representation of YlOrc4 (*blue*) and YlCdc6 (*pink*) near the Orc4/Cdc6 element of *OriC-061* in the YlORC-DNA^54bp*OriC-061*^–YlCdc6 structure, with insets of the YlOrc4 insertion helix (*bottom right*) and the YlCdc6 DNA binding loop (*top right, bottom left*). **(E)** A top-down representation of the complex near the AT element shows all proteins except YlOrc6 forming electrostatic interactions with *OriC-061* in the region.

#### Structure of YlORC-DNA*^OriC-^*^061^–YlCdc6 suggests sequence-specific binding

Cryo-electron microscopy (cryo-EM) was used to gain insight into the mechanism of YlORC origin binding. Using the 54bp DNA fragment of *OriC-061* in complex with YlORC and YlCdc6, a 2.7 Å resolution structure of the YlORC-DNA^54bp*OriC-061*^–YlCdc6 (ODC) complex was obtained (**Figure 4B, Suppl. Video 2 and Video 3, Figure S4B**). The winged-helix (WHD) and AAA+(-like) domains of all YlOrc1-5 and YlCdc6 proteins were visible in addition to the second TFIIB domain and C-terminal α-helix of YlOrc6 (**Figure S4C**). A somewhat weaker and less defined density for Orc2-WHD indicates flexibility of the domain while bound to DNA and Cdc6. In addition, neither the Orc1 bromo-adjacent homology (BAH) domain nor the N-terminal TFIIB domain of Orc6 were visible.

Akin to many AAA+ protein complexes and all ORC-Cdc6 structures determined to date ^68–70^, Orc1-5 and Cdc6 form a two-tiered hexameric ring with ATP-binding sites between the AAA+ domains of Cdc6/Orc1, Orc1/Orc4, Orc4/Orc5, and Orc3/Orc5, along with the contacts between the AAA+ domains of the other subunits forming the first tier (**Figure S4D**). ATP and Mg^2+^ were observed in the ATPase binding sites between Orc1/Orc4 and Cdc6/Orc1, while the Orc3/Orc5 and Orc4/5 sites contained ADP in the YlORC-DNA^54bp*OriC-061*^–YlCdc6 maps generated from gel filtration-derived samples (**Figure S4D**). The winged-helix domain of each subunit sits atop the adjacent subunit’s AAA+ domain, forming a domain-swapped second tier. Similar to the *S. cerevisiae* ODC (ScODC) structures ^68,70^, the C-terminal TFIIB domain (TFIIB-B) and C-terminal α-helix of Orc6 are visible. Like ScOrc6, YlOrc6 makes limited contacts with DNA and binds to the complex in multiple places: the TFIIB-B domain contacts part of the Orc2 N-terminal coil (residues 106-165), the WH domain of Orc3, and a small portion of the Orc5 basic patch (Orc5-BP, residues 348-364), and to the Orc3 protrusion with Orc6’s C-terminal α-helix. Differing from previous ODC structures, two small segments of the Cdc6 N-terminal IDR are bound to Orc1: Cdc6[1-13] binds to the exterior surface of the complex between Orc1-AAA and Orc4-AAA, while Cdc6[13-20] binds near the interface of Orc1-AAA and Cdc6-WHD (**Figure S4E**). Due to the partial occupancy of both segments within the density map, we suggest that this region can bind to Orc1 in either of the two conformations.

The sharpness of the DNA in the cryo-EM map was immediately evident, with purine and pyrimidine densities at each respective position easily discernible, and base identities apparent at most positions, indicating that the DNA is positioned in a discrete manner relative to the protein, implying that YlORC-Cdc6 binds to *Ori* sequences in a DNA sequence-dependent manner (**Figure S4F**). The pattern of identifiable bases was used to define its register. The DNA is significantly bent, similar to the DNA in ScODC structures ^68,70^, with a 40° bend occurring near the interface between the WHD and AAA+ domains of the complex.

#### YlORC/YlCdc6 binds *Ori* DNA specifically

From the structural analysis, the interactions between ORC/Cdc6 and DNA can be grouped into four sequence elements: A central region with major/minor groove and phosphate backbone contacts comprised predominantly of Orc4 and Cdc6 side chains we coined the Orc4/Cdc6-interacting element; the AT element consisting of minor groove and backbone interactions involving multiple subunits and a water-mediated hydrogen bonding network at one end of the origin; the Orc5 basic patch (Orc5-BP) element on the opposite end of the origin, with minor groove, backbone, and water-mediated hydrogen bonding interactions carried out by Orc2 and Orc5; and the Orc3 basic patch (Orc3-BP) element, which is between the Orc4/Cdc6 and Orc5-BP elements, at the bend in the DNA, and consists of minor groove and backbone interactions (**Figure 4C**).

The Orc4/Cdc6-interacting element is proximal to the large bend in the DNA and a site of significant minor groove compression. K465 of YlOrc4 makes sequence-specific contacts with the carbonyls of G31 and G32 in the major groove of the Y strand DNA (**Figures 4C and 4D, bottom right**). This lysine emanates from an α-helix in the insertion loop, similar in location to the α-helix of *S. cerevisiae* Orc4 (ScOrc4) in the ScODC complex that inserts itself into the major groove of the DNA for sequence-specific binding. The YlOrc4 insertion helix is considerably smaller than that of ScOrc4 (**Figure S4G**). It is angled so that a stretched K465 side chain reaches into the DNA major groove, and salt-bridge interactions between K462, R466, and D474 stabilize the helix. YlOrc4 K465 appears to form the only apparent sequence-specific interaction in the Orc4 insertion helix. Unlike the ScOrc4 insertion helix, which is highly conserved among fungi that are predicted to bind origins in a similar manner to *S. cerevisiae,* this lysine is not conserved (**Figure S4G**).

Near the YlOrc4 insertion helix resides a unique extended loop region of the Cdc6 WHD (**Figure 4D**). Sequence alignments to other eukaryotic Cdc6 proteins show that a large portion of the loop comes from an insertion that is strictly conserved within the family *Dipodascales*, but analysis shows that the existence of a similar feature may be generally conserved within fungi, with the relatives of *S. cerevisiae* being a noteworthy exception (**Figure S4H**). The density of this extended loop region is sharp and shows the loop interacting with the DNA over an entire turn of the double helix, making major/minor groove and backbone contacts throughout. Starting from the N-terminus of the loop, the sidechain of K548 is inserted into the minor groove and hydrogen bonds with the O2 carbonyl of Y-T29. R557 reaches into the major groove of the DNA and interacts with Y-G34 (**Figure 4D**). The DNA-binding loop appears to be held in place by several electrostatic interactions with the DNA extending from the Orc4/Cdc6-interacting element into the AT element. From these interactions, a structure-based preliminary binding motif was constructed centered around the Orc4/Cdc6-interacting element motif, deemed to be 5’-CNCCNRW-3’, where N denotes any nucleotide, R denotes purines, and W is A or T.

Located upstream of the Orc4/Cdc6-binding element, the AT element lacks major groove interactions and is recognized by a network of backbone, minor groove, and water-mediated hydrogen bond interactions involving all ODC subunits except Orc6 (**Figure 4C**). The most prominent sequence-dependent interaction in this element is between the Orc3 R220 sidechain and N3 of A15 via a coordinated water, and is further stabilized by an aspartate (D218) (**Figure S4I**). The positioning of Orc3 R220 likely precludes a C-G base pair at the adjacent position 16 due to potential steric clashes between R220 and the minor groove amine of a guanine. The sidechain of Orc4 K168 enters the minor groove near position 14 and may interact with the N3 nitrogen of A14, but the sidechain density is weak (**Figure S4I**). Adjacent to the AT element, the Orc1[300-305] basic patch is visible, with a mix of backbone and minor groove interactions. In addition, Orc2, Orc5, and Cdc6 all make contact with the phosphate backbone in this region. These interactions extend the binding motif in the 5’ direction of *OriC-061* to 5’-ATNNNNCNCCNRW-3’.

On the opposite side of the Orc4/Cdc6-binding element are the sites of two clusters of protein-DNA interactions. The first cluster encompasses X-strand positions 27 and 30 to 33 mediated predominantly by Orc3, which we named Orc3-BP. In addition to K671/K673 backbone interactions, Orc3 R672 forms a minor groove H-bond with T25 O2 carbonyl on the Y strand. Water molecules near X-A30 and the arginine sidechain expand the base specificity to having an A at that position (**Figure S4J**). The second cluster is mediated by Orc2 and the Orc5 basic patch (Orc5-BP) from positions X-35-41 (**Figure S4K**). Orc5 R357 forms a hydrogen bond with the O2 carbonyl of Y-T16, while R362 forms a hydrogen bond to the carbonyl of X-T38 (**Figure S4J**). The other significant interaction in the Orc5-BP element is an H-bond between Orc2 R150 and X-G35 in the major groove. With the addition of the Orc3-BP and the Orc5-BP elements, the inferred binding motif of YlORC and YlCdc6 to *OriC-061* extends in the 3’ direction to 5’-ATNNNNCNCCNRWNNANNNNGNNYR-3’, where Y denotes a pyrimidine.

#### Comparison of YlODC with two different origin sequences provide a consensus recognition sequence

To determine whether the interactions between YlORC/YlCdc6 and DNA are consistent between different origin sequences, a 2.6 Å cryo-EM structure was determined using a 60-bp fragment from *OriA-006,* the other origin validated by the linker scan assay (**Figures S4L, S4M**). Many of the critical interactions remained the same (**Figure S4N)**, with a Cα RMSD of 0.60 Å between the two structures, however, an excess of ATP was used for this cryo-EM sample and ATP is now visible in each ATP binding site.

In the AT element, the sidechain of Orc4 K168 appears to have stronger density overall compared to YlODC^54bp*OriC-061*^, but still appears to have multiple conformations. Unlike YlODC^54bp*OriC-061*^, the lysine sidechain could hydrogen bond to a water between K168 and the T51 O2 carbonyl on the Y strand, the O2 carbonyl of X-T11, or a water between the lysine amine and the N3 nitrogen of Y-A50. In the Orc3-BP element, Orc3 R672 is again coordinating Y-T34, with a water coordinated the corresponding adenine. The most significant change in the Orc5-BP element is the loss of Orc2 R150 density in the map with the change of G35 to a cytosine, consistent with the loss of this interaction.

The overall conformations of the Orc4 insertion helix and Cdc6 DNA-binding loop are very similar between the two origins, with Orc4 K465 coordinating two guanines. However, the positioning of the Cdc6 R557 sidechain is different: with, the arginine H-bonding with G34 on the Y strand in the major groove in *OriC-061*, while with *OriA-006*, R557 interacts with G44 on the Y strand, the equivalent of one base pair away (**Figure S4O**). This would then alter the Orc4/Cdc6 element motif from CNCCNRW in *OriC-061* to CNNCCNRW in *OriA-006*, pointing towards added flexibility for ORC/Cdc6 binding requirements. With the observed changes at other elements included, we suggest the structure-based YlORC- and YlCdc6-binding sequence 5’-ATNNNXXNCCNRWNNANNNNNNNYA-3’, where at least one X is a cytosine.

### Mutation of origin sequences disrupt ODC assembly

The importance of the sequence-specific interactions was determined by measuring the effects of origin mutations on *in vivo* genetic assays, including colony formation and plasmid stability (**Figure 5A**) and a biochemical assay for DNA/Cdc6 binding to ORC (**Figure 5B**). The first series of mutant *ori* fragments tested were in the Orc4/Cdc6-interacting element of *OriA-006*, as the site involves multiple major groove interactions which could confer base specificity. Changing individual cytosines (C17A or C21A) along with the complementary base in the motif does not affect the phenotype *in vivo*, but mutation of two cytosines [C20A, C21A] leads to smaller colonies, a 50% reduction in plasmid stability, and reduced Cdc6 loading (**Figure 5A, 5B and S5A**). All triple mutants (C◊A, C◊G, or C→T at positions 17, 20, and 21) are inviable *in vivo* (**Figure 5A and S5A**) and mutation of all three cytosines to guanines greatly reduced Cdc6 binding to ORC-DNA (**Figure 5B and S5A**).The lack of sensitivity to the X-strand position 17 mutant alone could be caused by the flexible nature of the Cdc6 R557 sidechain coordinating it, as it can potentially shift to the adjacent position as seen in *OriC-061* while still affecting binding specificity (**Figure S4N**). Changes to the *Ori* sequence at position 23, the site of the Cdc6 K548 interaction, led to a large decrease in the number of colonies, a 90% decrease in plasmid stability once returned to non-selective media (**Figure 5A**), and a sharp reduction of Cdc6 loading *in vitro* (**Figure 5B**), indicating the importance of this minor groove interaction for proper origin licensing. Mutations in other regions of the *Ori* sequence had similar effects. Both A12G and T13C mutations to the AT element of *OriA-006* result in a decrease in *in vitro* Cdc6 loading as do mutations to the Orc3-BP region (**Figure 5B**).

**Figure 5.**
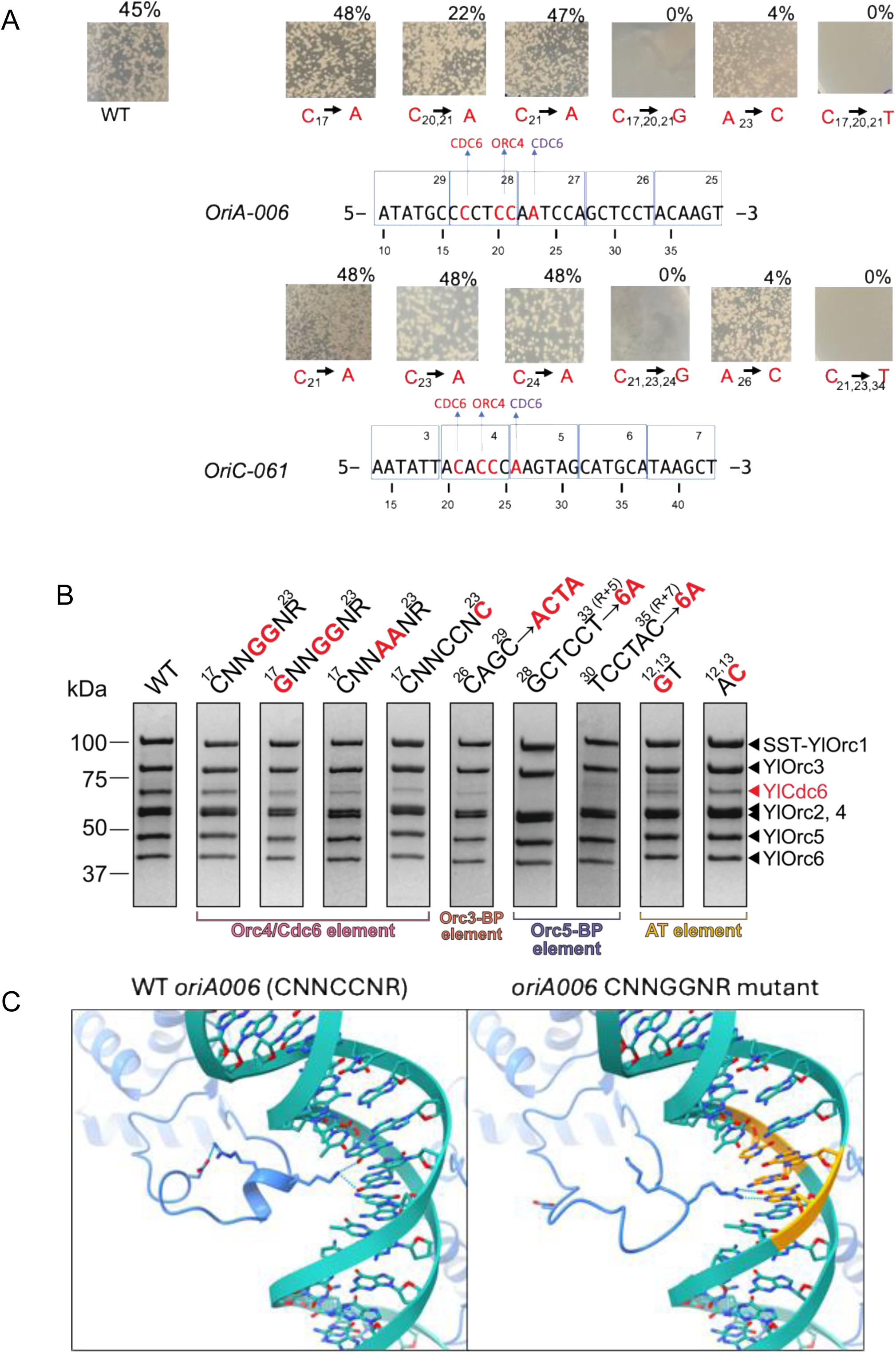
Effects of mutations on origin recognition and function. **(A)** Mutational analysis of the Orc4/Cdc6-interacting element motif in *OriA-006* (top) and *OriC-061* (bottom) and the effect on origin activity. Images show colony formation following plasmid transformation and selection and percentage (% URA+ retention) for different mutations. Base changes are highlighted in red, and their colony formation is shown in the adjacent images. Boxes represent the linker scan mutations (see Figure 3). **(B)** SDS-PAGE results of the peak fractions from the Cdc6 loading assay utilizing mutant *OriA-006* sequences, indicating the differences in Cdc6 co-elution (numbers refer to the X-strand). **(C)** Comparison of the structure of the YlOrc4 insertion loop between YlORC-DNA^60bp*OriA-006*^–YlCdc6 (*left*) and YlORC-DNA^60bp*OriA-CNNGGNR*^-YlCdc6 (*right*) structures. The yellow bases represent the bases that were mutated in *OriA-006*.

To study the structural effects of *Ori* sequence mutations on the binding of the ODC, a 2.6 Å resolution cryo-EM structure was determined for the double cytosine to guanine mutant [C20G, C21G] of *OriA-006* (YlODC^60bp*OriA-006*–CNNGGNR^) (**Figure S5D,E**). The structure is nearly identical to YlODC^60bp*OriA-006*–WT^, with the largest change occurring at the Orc4 α-helix (**Figure 5C**). The secondary structure of this helix unravels and becomes a structured loop. Orc4 K465, which contacts consecutive guanines in the WT *OriA-006*, points away from the DNA, and Orc4 D474 is unable to form a stabilizing salt bridge, as Orc4 R466 now forms multiple H-bonds with the mutant guanine, now located on the opposite strand. The Cdc6 DNA-binding loop appears to have no noticeable changes to its structure, suggesting that Cdc6 loading and specificity might be independent of Orc4 sequence-specific binding. Outside of this region, interactions between ORC/Cdc6 and DNA remain virtually identical.

### DNA deformability and bendability are critical for DNA replication

Compared to the strict sequence requirements for origin licensing in *S. cerevisiae*, *Y. lipolytica* has fewer essential base-specific contacts in the Orc4/Cdc6-binding element to allow for origin licensing. Are there other factors that could play a role in origin licensing specificity? Noticeable in the YlODC structure was a significant compression of the minor groove to allow the DNA to bend (**Figure S5K**). The bend is supported by interactions with the Orc5-BP (**Figure S5K**), as was also shown for *S. cerevisiae*. If the bendability of the *Ori* has a significant role in origin licensing specificity, replacing segments of the bent region of the *Ori* DNA with a rigid dA tract should inhibit origin licensing ^77,78^. The structural properties of DNA have been suggested previously to play a part in *Drosophila* ORC binding preference, and more recent studies have shown that bending of DNA via the Orc5-BP was critical for DNA replication in *S. cerevisiae* ^25,71,79^. To test this, we examined the effect on DNA binding, Cdc6 loading, and the *in vivo* effects of incorporating a 6 nt-long dA tract around the region of *OriA-006* that is bent in the YlODC structure, starting at position 23 (abbreviated as A[23-28]) and moving it downstream 1 bp at a time (**Figure S5B**).

There were discrepancies between the Cdc6 loading assay and the *in vivo* phenotypic effects of the 6A mutants tested (**Figure S5B and S5C).** At A[24-29] a decrease in Cdc6 loading was seen, but no deleterious effects were observed *in vivo*. Mutant A[25-30] had >85% of WT *OriA-006* activity, Cdc6 loading efficiency increased, whereas with A[26-31] and A[27-32] Cdc6 loading assay results had near WT Cdc6 loading efficacy, but either an extreme reduction or a complete lack of growth was observed *in vivo*. Insertions from A[28-33] through A[31-36], showed complete inhibition of Cdc6 loading and no growth *in vivo*, as the 6A tract enters the Orc5-BP element (**Figure 5B**). Decreases in the 260/280 ratio, indicating reduced DNA binding, were seen in the A[28-33] through A[31-36] mutants compared to WT.

### Large-scale mutational analysis of *Yarrowia* Origins of DNA Replication

To identify sequence preferences required for origin function, a Massively Parallel Origin Selection (MPOS) assay was performed using mutagenized libraries (∼15% mutation density ^35^) spanning 90 bp regions of *OriC-061* and *OriA-006*. These libraries were cloned into *Ori*⁻/CEN⁺ plasmids and introduced into *Y. lipolytica* for growth-based competitive selection. Deep sequencing of plasmids from pre-selection and post-selection time points was performed, and a quantitative model was trained using MAVE-NN to predict post-selection enrichment as a function of DNA sequence ^80^.

In *OriA-006*, the most striking changes occurred between positions 10 and 37, where specific nucleotides showed strong selection signatures, highlighting this region as a functionally critical core of the origin, which aligns with the linker scan sensitive area (**Figure 6A**, the numbers refer to the numbering of base pairs in **Figure S4N**). A similar pattern was observed for *OriC-061*, though with slightly lower resolution due to reduced transformation efficiency and higher background signal (**Figure S6A**), confirming that both sequence preference and positional sensitivity are conserved features in this origin.

**Figure 6.**
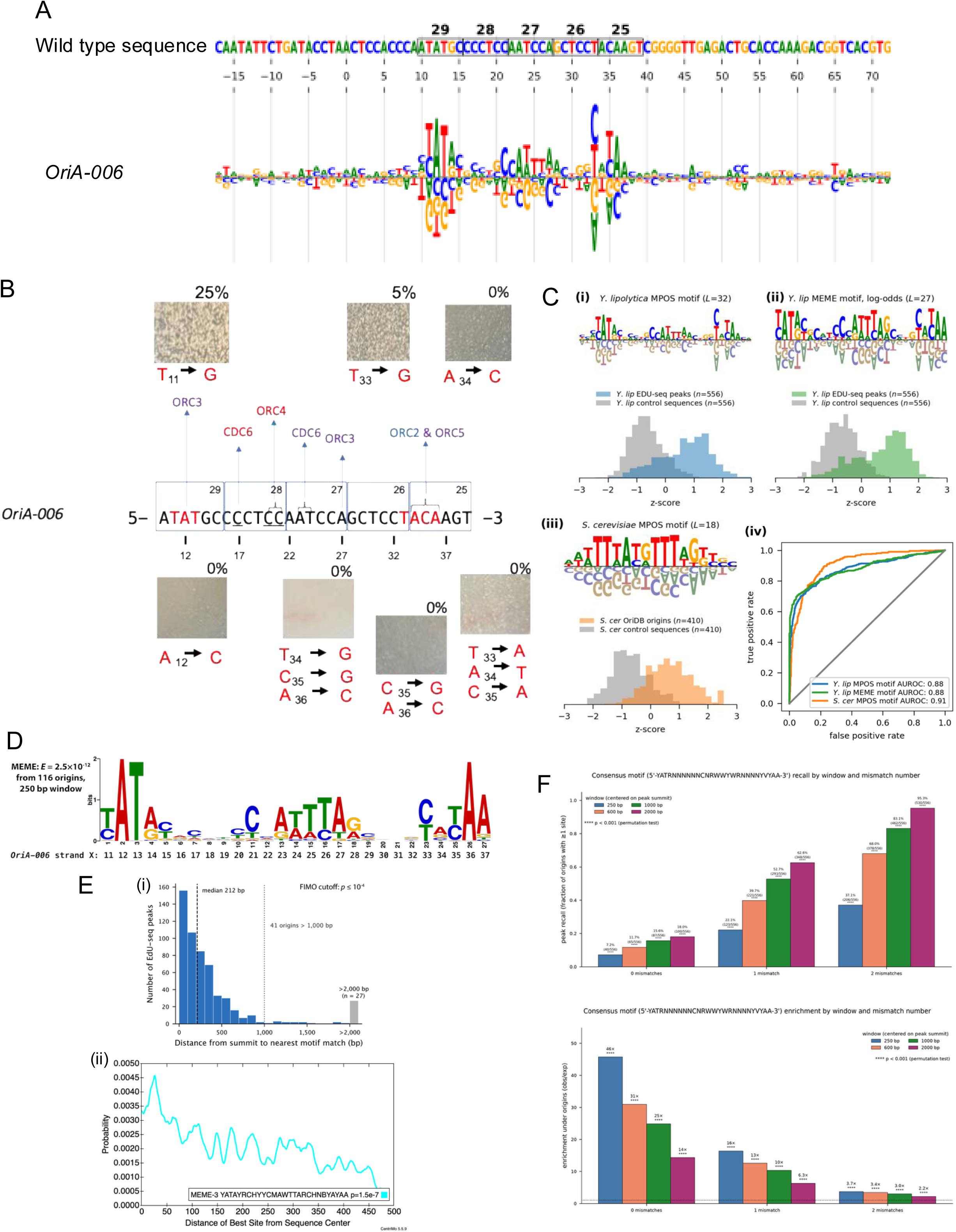
Massively parallel origin selection assay and quantitative modeling. **(A)** Quantitative model for *Y. lipolytica* origin specificity derived from a massively parallel origin selection (MPOS) assay carried out on 90 bp sequences containing *OriA-006* mutagenized at 15% per bp. Logo illustrates an additive model trained using MAVE-NN ^80^ to distinguish selected variants from input variants. Sequence coordinates match those in **Fig. S4M**. The endogenous *OriA-006* sequence is shown above, and linker positions 25-29 from Fig. 3B are boxed. **Fig S6A** provides a similar analysis for *OriC-061*. **(B)** Functional origin assay of single-nucleotide mutants within the AT and the Orc5-BP motifs. Images show colony formation following plasmid transformation and selection and percentage (% URA+ retention) for different mutations. **(C)** Sensitivity and specificity of core motifs in MPOS-derived models. (i) Core motif of the *Y. lipolytica* MPOS model and corresponding z-score distributions for EdU peaks and random *Y. lipolytica* genomic regions. (ii) The log-odds *Y. lipolytica de novo* MEME motif and corresponding z-score distributions for EdU peaks and random *Y. lipolytica* genomic regions. (iii) Core motif from a model trained on *S. cerevisiae* MPOS data ^28^, together with z-score distributions of this model on origins from OriDB ^18^ and on random *S. cerevisiae* genomic regions. (iv) ROC curves for the three motifs on their respective positive and negative genomic regions. **(D)** A *de novo* motif generated using MEME in differential enrichment mode compared to dinucleotide-preserved shuffled genomic sequences, with enrichment confirmed using AME analysis (*p* < 10^−37^ at all window sizes). Equivalent positions to those on *OriA-006* are shown. **(E)** Positioning of de novo motif hits within peaks. (i) Distance of *de novo* motif matches called by FIMO (cutoff: p < 10^−4^) from EdU-seq peak summits. (ii) CentriMo analysis illustrating the enrichment of *de novo* motif matches near peak summits (central window width = 236 bp). **(F)** Consensus motif matches and enrichment (calculated as observed/expected from whole-genome hit frequency) using multiple window sizes and mismatch tolerances, with p-values calculated from a one-sided Monte Carlo permutation test using per-chromosome shuffles of the origin windows (2,000 iterations preserving window number, size, and total coverage; empirical bias-corrected p = ((# of iterations where shuffled windows outperform origin peak windows)+1)/2001). This analysis was carried out using Claude Science and manually validated.

Bases critical for ORC–CDC6 binding showed enrichment when unmutated and depletion when altered, reflecting their essential role in origin establishment. Conversely, mutations that allowed initial transformation but led to reduced plasmid stability highlighted bases important for maintenance. This dual-phase insight was central to parsing the functional logic of ARS activity in *Y. lipolytica*.

A conserved sequence motif, 5’-YATRNNNNNNCNAWTTNNNNNNYNYAA-3’, emerged from MPOS assay analysis as a central feature of functional origins. Targeted mutagenesis within the YAT, YNYA, and flanking regions revealed key nucleotide requirements for *Ori* function (**Figure 6B**). Mutation in a critical position such as A12-to-C within the YAT region abolished colony formation, indicating a loss of *Ori* activity. Multiple substitutions in the YNYA region, including mutations at positions 33, 34, 35 and 36 resulted in either reduced plasmid instability or no transformation, suggesting that these positions are critical for origin activity. These findings align with MPOS results, which showed selection against guanine (G) at T33, reinforcing a preference for thymine (T) or cytosine (C). This preference likely reflects the structural compatibility of pyrimidines with ORC binding, particularly through minor groove interactions involving ORC3, ORC2, and ORC5. For example, the YA motif appears critical for minor groove engagement by ORC5, and substitutions that disrupt this local DNA shape abolish origin function.

### Cross-Species Validation of a Replication Origin Motif Using ROC Analysis

Next we evaluated the predictive power of MPOS-identified motifs by evaluating their ability to distinguish functional origins from genomic background. The motif derived from the *Y. lipolytica* MPOS assay exhibited a remarkable ability to distinguish EDU-seq peaks from randomly sampled DNA from the *Y. lipolytica* genome, yielding an AUROC of 0.88 (**Figure 6C**). By comparison, an AUROC of 0.91 was obtained when an analogous motif was inferred from *S. cerevisiae* MPOS data of ^28^ and used to distinguish sequences in OriDB ^18^ from random DNA in the *S. cerevisiae* genome (**Figure 6C**).

An analysis of nucleotide frequencies across 54 *Y. lipolytica* early origins of replication compared to the frequencies across 5351 close matches to the motif in non-origin sequences showed a skew for T/A sequences 5’ to the core consensus sequence and a skew for A/T base pairs 3’ to the core consensus sequence only in early origins, but not in non-origins (**Figure S6B**). This suggests that the arrangement facilitates initiation of DNA replication in the generally G/C rich genome since the average %G/C in 160 bp around motif in early origins is 44%, whereas the *Yarrowia* genome is on average 48.9% G/C ^73^.

### *De novo* motif discovery of origins reconciles structural and MPOS-derived motifs

Using a 250bp window centered on the full set of EdU-seq peak calls, *de novo* motif generation using MEME was used to identify 20-30 bp origin-enriched motifs. The second-highest enriched motif closely resembled previous iterations of the origin sequence motif we described and was found to be highly enriched in origins, with near-identical motifs reproduced using both 600 bp and 1000 bp windows around the peaks using MEME-ChIP (**Figure 6D**). Within this *de novo* motif, specific positions are seen as consistent across origins, equivalent to X-12, X13, X-33, X-35, and X-36 in *OriA-006*, with other sites more flexible (**Figure 6D**). The vast majority of origins contain sequences consistent with this motif, with only 41 peaks that are not within 1000 bp of a match with a p-value of less than 10^−4^ (**Figure 6E**). The *de novo* motif appears to reconcile both the structural- and MPOS-based motifs. It is consistent with the MPOS data for the enrichment of positions at the edges of sequence elements that lacked concrete structural observations (**Figure 6C, 6D**. On the other hand, the site in *OriA-006* coordinated by the flexible Cdc6 R557 (X-17) shows modest enrichment for a cytosine at that position.

To confirm that matches to the *de novo* motif correspond to true origins, three putative origins, the highest scoring match (*OriE-033*) and two low-scoring origins (OriE-062 and OriE-061) were tested for Cdc6 loading. All three were found to bind ORC and load Cdc6 to varying extents, supporting that the de novo motif matches support YlODC complex formation **(Fig S6C)**.

Based on these observations, along with the structural and MPOS data, the consensus motif for YlODC is 5’-YATRNNNNNNCNRWWYWRNNNNYVYAA-3’. Allowing for up to two mismatches expands the number of peaks that contain a match, >80% within a 1000 bp window, all while maintaining significance (*p* < 0.001) for recall and enrichment over randomly placed windows of equal size in the genome for all window sizes and mismatch tolerances (**Figure 6F**). Motif mismatches are expected: only one putative origin tested, *OriE-066*, matches the consensus sequence perfectly, with all others containing one or two mismatches at varying locations across the motif (**Figure S6C**). This supports the conclusion that *Yarrowia lipolytica* origin licensing is very flexible and does not require exact consensus sequence matches for efficient origin licensing.

## Discussion

The *Yarrowia lipolytica* genome has at least 556 origins of DNA replication that are distributed into large replication timing domains of early and late replicating regions, much like the A and B replication timing domains of vertebrate chromosome replication ^81,82^. Interestingly, both centromeres and telomeres replicate early in *Y. lipolytica*. The early and late replicating domains in vertebrates, including in human cells, correspond to topologically associated domains (TADs) and euchromatin and heterochromatin, respectively. What determines the structure of the *Y. lipolytica* replication timing domains remains to be determined, but the global temporal pattern of replication in this yeast is very different from the replication timing in *S. cerevisiae*, which falls into two classes based on temporal control by the S-phase cyclin-dependent protein kinases ^83^.

Origins of DNA replication in *S. cerevisiae* are specified by ORC recognizing the ARS consensus sequence 5’-WTTTAYRTTTW-3’and bending the DNA ^17,31,84^. After interacting with ORC, the Cdc6 initiator-specific motif (ISM) and WH domains bind to the DNA phosphate backbone (but do not form base-specific interactions), thereby contributing to origin DNA binding ^68,70^. Thus, in *S. cerevisiae*, the location of origins in the genome is primarily due to ORC. The ScOrc4 and ScOrc2 base-specific contacts are major contributors to DNA-sequence-specific interactions, so much so that it was predicted that the presence of the Orc4 α-helix correlated with DNA sequence-specific binding ^28,32^. But *S. cerevisiae* ORC also has minor groove and backbone interactions that contribute to ORC-DNA binding. Clearly, however, analysis of origin recognition in *Y. lipolytica* suggests that there are alternative mechanisms for base-specific origin recognition because in this yeast both YlORC and YlCdc6 are required. The *Y. lipolytica* consensus sequence, 5’-YATRNNNNNNCNRWWYWRNNNNYVYAA-3’, supported by both structural data and high-throughput mutagenesis data, is substantially different from the *S. cerevisiae* sequence, both in length and flexible base composition. The YlOrc4 α-helix is substantially reduced and only a single lysine (K465) contacts adjacent G/C base pairs in the 5’-CCNRW-3’ Orc4/Cdc6-interaction motif. This α-helix is completely missing in the human ORC4 subunit. Mutation of single YlOrc4 interacting bases did not affect *Y. lipolytica* origin activity (**Figure 5A**), in contrast to single base changes in the ScORC binding site significantly compromising origin activity ^85^. When two adjacent YlOrc4 interacting base pairs are mutated, the Orc4 α-helix collapses and the base-interacting K465 residue, switches with R466, which now makes base-specific interactions, albeit with guanines now located on the opposite strand (**Figure 5C)**. Thus, there is considerable plasticity in ORC binding to origins in *Y. lipolytica* since the different origins display alternative protein-DNA interactions.

We suggest that the ancestral ORC4 lacked the α-helix and its associated base-specific interactions, but in budding yeasts there has been a gradual evolution of the Orc4 protein toward increasing sequence-specific origin recognition (**Figure 7**). This may be due to the smaller genome size in fungi, where DNA sequence-sequence origins need to be placed in the small intergeneic regions so as to avoid conflicts between the initiation of DNA replication and gene transcription. Indeed we observed that most origins in *Y. lipolytica* are in intergenic regions of the genome, as they are in *S. cerevisiae*.

**Figure 7.**
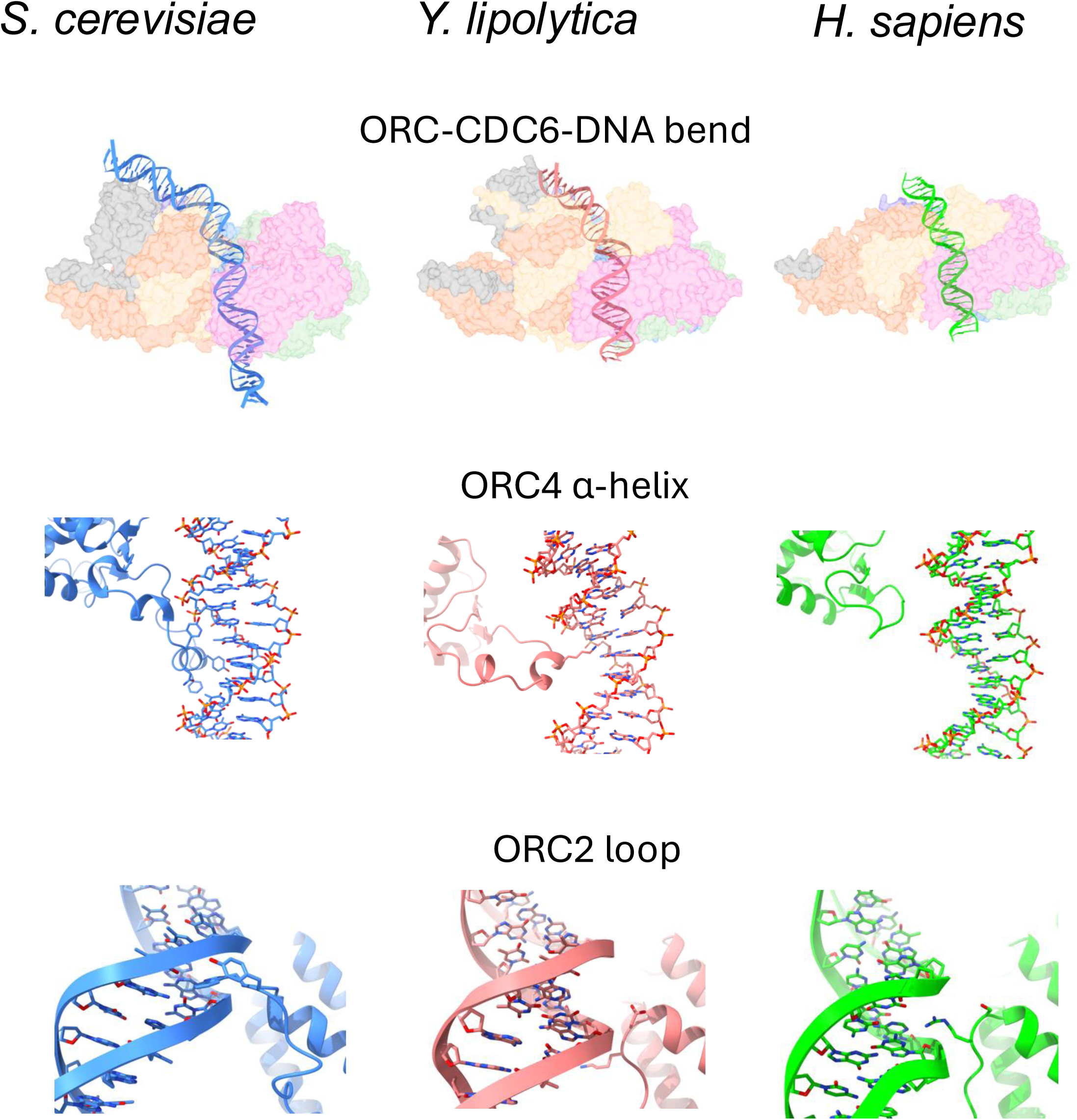
Origin recognition throughout evolution. A comparison of analogous structural features within the ODC complex of metazoans (*Homo sapiens*), *Saccharomyces cerevisiae,* and *Yarrowia lipolytica*.

Unlike the situation in *S. cerevisiae*, the YlORC bound weakly to origin DNA and YlCdc6 increased the specificity and affinity for origin DNA by making several contacts, including a base-specific interaction via R557 to the Orc4/Cdc6-interacting element, adjacent to the Orc4 α-helix interaction site (**Figure 4D**). Interestingly, a single base change in the base in the major grove of either *OriA-006 or OriC-061* that interacts with YlCdc6 R557 does not affect origin function. This may be due to the flexibility of the YlCdc6 R557 ability to interact with neighboring base-pairs in the major groove, depending on the origin (**Figure S4N**). In contrast, in both origins, mutation in the minor groove base pair that interacts with YlCdc6 K548 severely compromises origin activity, underscoring the importance of YlCdc6 in origin recognition.

The extensive DNA interactions in the AT and Orc5-BP elements in the consensus sequence contribute to origin function, since single-point mutations in both elements eliminate origin activity. These include base pairs that interact with Orc3 in the AT element and Orc2 and Orc5 in the Orc5-BP element. An Orc1 loop (residues 300-305) interacts with the minor groove non-specifically in addition to other residues binding the phosphate backbone. A conserved β-loop in this region binds to a minor groove in the *Drosophila* ORC-CDC6-DNA structure, but does not make nucleotide-specific contacts ^69^. Indeed, unlike *Y. lipolytica* ORC-Cdc6 interactions with DNA, all DNA contacts in the *Drosophila* ORC-CDC6-DNA structure on a 60 bp AT-rich DNA lack base pair specificity. One common feature in all structures, however, is the interaction between Orc5/ORC5 with the DNA to stabilize the DNA bend (**Figure 7**). As noted previously ^25,86–88^, it is possible that DNA sequences that have the propensity to bend upon ORC or ORC-CDC6 binding is an essential feature of all origins of DNA replication.

It was surprising that, in the structure of the human ORC-CDC6-DNA, we observed that HsORC2 residue R367 appears to interact in a minor grove with the edge of a nucleotide, suggesting for the first time that human ORC-CDC6 might have direct DNA sequence dependent interactions or origin recognition. In an analogous way, the *S. cerevisiae* Orc2 W396 residue makes π-interactions in a minor groove of *ARS1* origin DNA, contributing to origin specificity. In contrast, in *Yarrowia*, residues in the non-conserved Orc2 loop interact with the phosphate backbone of DNA, likely contributing to affinity but not specificity (**Figure 7**). The DNA in the human ORC-CDC6-DNA structure is a G/C rich sequence that is not an origin of DNA replication or a known ORC-CDC6 binding site in the human genome, so it is possible that ORC-CDC6 interaction with an authentic origin or binding site might reveal additional base-specific interactions.

It is remarkable that for such a fundamental process as specification of origins of DNA replication that there have been many solutions to defining the location of origin sequences in the eukaryotic genome. This variation may be a result of differences in genome organization and gene density, as well as other features such as genome size, G/C content and 3D structure. The *Yarrowia* and human ORC-Cdc6-DNA structures presented here, together with previous analyses of *S. cerevisiae* and *Drosophila* structures have highlighted some key aspects of the evolution of origin recognition and specification.

### Limitations of the study

We have only characterized in detail two *Y. lipolytica* origins and yet there are some sequences under EdU peaks that lack matches to the *de novo* motif and/or the consensus sequence identified here. 41 (7.4%) origins lack a *de novo* motif hit and 26 (4.7%) lack a consensus motif match when allowing up to two mismatches within 1000 bp. Conversely, some origins may contain matches that fail to pass the *de novo* motif cutoff (*p* < 1e-4) but still load Cdc6, as seen with *OriE-081*, suggesting that the p-value cutoff may be too strict and that additional context-dependent factors may prevent some motif matches from becoming active origins. Mutagenesis and structural studies with ORC and Cdc6 using these origins may reveal even greater flexibility in origin specification uncovered in this study. Furthermore, continued analysis of the HsODC using different DNA sequences, including known ORC-CDC6 DNA binding sites, may reveal additional base-specific origins recognition in human cells.

## Acknowledgements

This research was supported by grants from the National Institutes of Health (GM045436, GM133777, HG011787), the Howard Hughes Medical Institute and the Goldring Family Foundation. Core DNA sequencing was facilitated by the Cold Spring Harbor Laboratory (CSHL) DNA sequencing and analysis shared resource, supported by the Cancer Center grant (CA13106). We thank Dennis Thomas for managing the CSHL Cryogenic Microscopy Core shared resource, and members of the Stillman and Joshua-Tor laboratories for suggestions and advice. L.J. is an Investigator of the Howard Hughes Medical Institute.

## Author contributions

B.S. and L. J. conceived the study. B.S., L. J., and J.B.K. designed the study. J.B, N.Z., O.P.C. and K.L. performed the experiments. J.B., N.Z., O.E.D, O.P.C., K.L., J.B.K., L.J., and B.S. analyzed the data. N.Z, J.B, O.P.C, J.B.K, L.J. and B.S. wrote the paper with input from all authors. B.S., J.B.K. and L. J. provided funding and oversaw the project.

## Methods

### Yeast Strain Construction

Yeast strains for EdU-sequencing and ARS assays were derived from *Yarrowia lipolytica* PO1f (MATA, leu2-270, ura3-302, xpr2-322, axp1-2). Strains used for genome annotation were provided by Dr. Richard Rachubinski (University of Alberta). To generate the TK+ strain (YlB0002), a BrdU-Inc cassette with a *URA3* marker was integrated at the *IntE1* locus on chromosome 5. The cassette, under TEF and GPD promoters, expressed Herpes Simplex Virus thymidine kinase (HSV-TK) and the human Equilibrative Nucleoside Transporter 1 (hENT1) genes. Correct integration was confirmed via selective growth on -URA and FOA plates, colony PCR, and vector verification before transformation.

### Plasmid Construction for BrdU-Inc Cassette

The BrdU-Inc cassette was cloned into the EasyClone vector pCfB6677 ^89^ obtained from Addgene and targeted to the *IntE1* locus. The cassette was flanked by loxP sites for *URA3* marker removal via Cre recombinase. HSV-TK and hENT1 were placed under the TEF and GPD promoters, respectively, and amplified from pNC1164 and pCfB8742. USER cloning enabled precise assembly using uracil-containing primers and enzymatic treatment to generate overhangs for directional ligation. The final construct was validated by PCR, restriction mapping, and sequencing.

### ARS Plasmid Construction

The pSCARS1 plasmid ^90^ was modified to create pYl001 by removing SC-Trp and ORI1068, and adding KpnI and BglII sites. GFP remained under TEF promoter control. To construct pYl002 - pYl011, replication origins were amplified from PO1f genomic DNA and ligated into pYl001 at KpnI or BglII sites. Constructs were verified by PCR and restriction digestion for downstream ARS assays and stability tests.

### Yeast Transformation

PO1f cells (5 × 10⁷) were grown overnight and transformed according to Dahlin et al 2021 ^89^. For genomic integrations, 500 ng of the linearized vector was used; for ARS assays, 15 µg of the circular plasmid. Cells were heat-shocked at 39°C, recovered in YPD, and plated on -URA. Integration was confirmed using PCR.

### Synchronization of YlB0002 (TK+)

To synchronize cells in G0/G1, YlB0002 was grown in YPD for 72 hours when they reached stationary phase and then diluted 1:10 into fresh medium to re-enter the cell cycle. EdU (500 µM for sequencing, 100 µM for imaging) (Thermo Fisher, E10187) and HU (5 mM) (Sigma, H8627) were added as needed. Samples were collected at multiple time points for EdU imaging, flow cytometry, and sequencing.

### EdU Imaging in YlB0002

Synchronized or log-phase cells were labeled with 100 µM EdU (Thermo Fisher, E10187). After fixation (3.7% PFA), cells were permeabilized with Triton X-100 and subjected to Click-iT chemistry using Alexa Fluor 488 (Thermo Fisher, C10387). DNA was stained with Hoechst 33342, and cells were mounted in anti-fade solution for imaging with 63X or 100X oil objectives.

### Flow Cytometry

To assess cell cycle progression in *Y. lipolytica*, cultures were grown at 30°C to mid-log phase or harvested at specific time points. Cells were pelleted and washed twice with sterile water, then fixed in 70% ethanol and incubated overnight at 4°C. Following fixation, cells were pelleted, washed twice with sterile water, and resuspended in FC buffer (50 mM sodium citrate, pH 7.0, 0.1% sodium azide). For RNA and protein degradation, samples were sequentially treated with RNase A (0.1 mg/ml) and proteinase K (0.2 mg/ml) for 1 hour each at 55°C. Cells were then stained with SYTOX Green Nucleic Acid stain (Thermo Fisher S7020). Samples were sonicated and diluted before flow cytometry or FACS analysis to assess DNA content and cell cycle distribution.

### EdU-Seq of Synchronous *Yarrowia* Cells ± HU

To track DNA replication dynamics, *Y. lipolytica* cells were synchronized in G0/G1 by 72-hour culture in YPD. Cells were then released into fresh YPD containing 500 µM EdU (± 5 mM HU), and samples were collected over a 210-minute time course. Flow cytometry confirmed synchronization.

For mapping sites of EdU incorporation, DNA from the time-course samples was fragmented, EdU-labeled DNA was captured via biotin-azide Click-iT reaction (Thermo Fisher, C10365) and Streptavidin T1 beads (Thermo Fisher 65602). Libraries were prepared using the Illumina TruSeq Kit (Illumina IP-202-1012). Sequencing identified newly replicated regions, providing high-resolution replication timing profiles. Sequencing data revealed replication origin firing across S phase ^35^.

### EdU Incorporation and DNA Preparation

To identify early- and late-firing origins, stationary phase cells were transferred into fresh YPD containing either EdU+HU or EdU alone. A mock (no EdU) control was included. At each time point, replication was halted with sodium azide (0.1%), and cells were collected for flow cytometry and DNA extraction. DNA was purified using a Qiagen Genomic-tip after Zymolyase-20T (Sunrise Science Products, N0766391) digestion and lysis. Purified genomic DNA was fragmented using the Bioruptor Pico and checked on a Bioanalyzer (100–550 bp fragments).

### Click Labeling and DNA Enrichment

EdU-labeled DNA was tagged with biotin via Click-iT chemistry and purified using magnetic Streptavidin beads. Bound DNA was eluted and further cleaned using MinElute columns (Qiagen).

### Library Prep and Sequencing

Biotinylated DNA was ligated to adapters and PCR-amplified using the Illumina TruSeq kit. The resulting libraries were sequenced to profile replication timing at high resolution.

### EdU-seq Pre-processing and Genome Alignment

The paired-end reads were trimmed using Fastp v.0.23.2 ^91^ using the default settings and automatic adapter detection, and quality control was performed using FastQC (v0.12.1; available online at: http://www.bioinformatics.babraham.ac.uk/projects/fastqc/). The trimmed reads were aligned to *Yarrowia Lipolytica* strain E122 (MATA) using Bowtie2 v.2.4.4 ^92^ with the default options. The output SAM alignment files were converted to BAM format, sorted and indexed using SAMtools Samtools v.1.19.2 ^93^ Additionally, BAM files were filtered for proper pairs and minimum quality scores of q30; multiple mappings, unmapped queries, and duplicate reads were dropped. BAM coverages were normalized using counts per million mapped reads (CPM) then converted to bigWig (.bw) format to produce illustrated genome coverage tracks.

To produce the illustrated genome coverage tracks and for visualization purposes, we used bamCoverage from Deeptools v.3.5.1 ^94^ was used to generate the coverage tracks with normalization option of bins per million mapped reads (BPM).

A total of 75 EdU-seq libraries were prepared across 11 time points (15–210 minutes), with biological and technical replicates ensuring data reproducibility (**Supplement Table 1**). Fifty high-quality libraries were selected for downstream analysis. Temporal replication patterns were consistent across replicates, demonstrating the robustness of the protocol.

**Supplemental Table 1:**

| Time points | Number of libraries for each time point |
| --- | --- |
| 15 minutes | 7 |
| 30 minutes | 12 |
| 45 minutes | 10 |
| 60 minutes | 3 |
| 75 minutes | 4 |
| 90 minutes | 4 |
| 120 minutes | 4 |
| 150 minutes | 2 |
| 180 minutes | 4 |
| 210 minutes | 4 |
| Total | 75 libraries from which these 50 were<br>chosen (excluding all the Mock samples) |

**Supplemental Table 1**
| Accession Number | File name | Hydroxyurea | Time (minutes) |
| --- | --- | --- | --- |
| GSM9595226 | 30min_S4_trm.bw | No | 30 |
| GSM9595227 | S03_S3_trm.bw | Yes | 45 (# 1) |
| GSM9595228 | NZ2770_S07_S7_trm.bw | Yes | 45 (#3) |
| GSM9595229 | S14_S14_trm.bw | Yes | 120 |
| GSM9595230 | S04_S4_trm.bw | Yes | 60 |
| GSM9595231 | 72-45-eh-5_S3_trm.bw | Yes | 45 (#2) |
| GSM9595232 | S07_S7_trm.bw | Yes | 90 (#1) |
| GSM9595233 | S09_S9_trm.bw | Yes | 75 |
| GSM9595234 | S11_S11_trm.bw | Yes | 180 |
| GSM9595235 | 9-30min-eh5_S3_trm.bw | Yes | 30 |
| GSM9595236 | 45min_S5_trm.bw | No | 45 |
| GSM9595237 | 2770_S04_S4_trm.bw | No | 60 |
| GSM9595238 | 2818_S06_S6_trm.bw | No | 20 |
| GSM9595239 | S12_S12_trm.bw | Yes | 210 |
| GSM9595240 | S08_S8_trm.bw | Yes | 90 (#2) |
| GSM9595241 | S06_S6_trm.bw | No | 120 |

### EdU-seq Peak calling and annotation

For peak calling, we employed MACS3 ^95^ on the paired-end BAM files for each EdU+HU timepoint up to 180 minutes using a q-value filter of 0.05 (0.01 for the 30-minute timepoint) and a minimum length of 400 bp, with a maximum gap of 1800 bp and optimization of background calculation parameters (--slocal 500 –llocal 30000) to prevent duplicate peak calling or peaks called due to the noisiness of the BAM files at later time points (--slocal and llocal parameter optimization was done with the assistance of Claude Science (Anthropic. Claude Opus 4.8 (Claude Science 0.1.22) [Large language model]. https://claude.com/product/claude-science (2026)). Reproducible and consistent calls across time points were then aggregated, with the median peak summit used for anchoring. From there, BAM files for early (30, 45, 60 min), medium (60, 75, 90 min), and late (90, 120, 180 min) time points were depth-normalized, pooled, and underwent a round of peak calling (max gap reduced to 1250 bp). This allowed for stricter quality control. the calling of additional late origins that did not reach the signal threshold in individual timepoints, adjusting the position of the anchors with pooled early and medium peak calls when the signal amplitude was stronger in pooled peaks than in the anchor, the splitting of nearby peaks that converged in later timepoints, and for manual review of anchors that were not called in any pooled segment. The final peak set was then centered on the summit of each peak.

The final peak set was annotated using annotatePeaks.pl from HOMER v5.1 http://homer.ucsd.edu/homer^96^, which also provides the annotation class (Promoter-TSS/TTS/Gene/Intergenic) enrichment analysis.

### Stability Assays

For plasmid stability, transformants were grown in selective media, then diluted into YPD and regrown. After 30 hours, cells were plated on selective and non-selective plates. Colony counts on both plates were used to estimate plasmid loss, expressed as percentage. In GFP-based assays, cells harboring plasmids expressing Green Fluorescent Protein (GFP) under the control of the *TEF* promoter with the *CYC1* terminator were analyzed via flow cytometry according to Lopez et al. ^90^. GFP+ percentages were calculated using BD FACSC, with wild-type cells as a negative control.

### XhoI Linker Scanning via In-Fusion Cloning

XhoI linker scanning mutants of *OriA-006* and *OriC-061* were generated using In-Fusion cloning (TaKaRa 638945). Primers with XhoI sites and 15-bp overlaps were used in high-fidelity PCR (PrimeSTAR Max,TaKaRa R045A). Mutant plasmids were circularized via inverse PCR and In-Fusion assembled, purified, and transformed into *Y. lipolytica* for downstream analysis.

### Massively parallel origin selection (MPOS) assay

MPOS assays were carried out as in ^28^ with minor modifications. Plasmid libraries for OriA-006 and for OriC-061 were constructed starting with corresponding wild-type plasmids used in the linker scanning experiments. 90 bp regions centered on the essential sites identified by linker scanning were then mutagenized at 15% per bp. Mutagenesis was carried out using computationally designed oligo pools, each containing 7,000 variants, synthesized by Agilent. To increase the percentage of correctly cloned plasmids, variants were cloned using a *ccdB* cassette replacement strategy based on that of Kinney et al. (2010) ^97^, but with Gibson cloning instead of Golden Gate cloning. Each plasmid library was then transformed into *Y. lipolytica* and subjected to selection via growth on SC-URA plates, after which bulk DNA was extracted. Amplicons containing variant sequences flanked by barcodes and primers for Illumina sequencing were then prepared using template DNA (plasmid DNA from the initial libraries or bulk DNA extracted from cells) amplified by PCR using custom primers. Amplicons were subjected to PE150 sequencing on an Illumina NextSeq 2000 instrument using a P1 flow cell and XLEAP-SBS chemistry.

### Quantitative modeling of MPOS data

For each of the two loci, *OriA-006* and *OriC-061*, MAVE-NN ^80^ was used to train an additive model that distinguishes MPOS-selected origins from those in the initial library. The *OriA-006* model was trained using 2,911,260 pre-selection reads and 2,082,266 post-selection reads. The *OriC-061* model was trained using 8,820,054 pre-selection reads and 7,264,362 post-selection reads. Sequence logos illustrating these models are shown in **Figure 6A** (*OriA-006*) and **Figure S6A** (*OriC-061*). Logos were created using Logomaker ^98^. The additive model for origin specificity in S. *cerevisiae* was previously reported by Hu *et al.* ^35^ and computed in a similar manner using an early version of MAVE-NN.

### ROC analysis of MPOS-derived motifs

To carry out the ROC analysis in **Figure 6C** a 32 bp core motif was extracted from the *Y. lipolytica OriA-006* additive model. This motif was used to scan 1,000 bp regions of *Y. lipolytica* genomic DNA centered on either the EdU peaks identified above (positive set; 623 regions) or on randomly chosen genomic locations (control set; 623 regions). The maximum motif score observed in each genomic region was recorded. The scores for positive and negative regions were then converted to z-scores (**Figure 6Ci**). A similar analysis was performed using an 18 bp core motif extracted from the *S. cerevisiae ARS1* MPOS model (**Figure 6Cii**). This motif was used to scan 1,000 bp regions of S. cerevisiae genomic DNA centered either on origins in OriDB ^18^ (positive set; 410 regions) or on randomly chosen genomic DNA (control set; 410 regions). **Figure 6Ciii** shows ROC curves corresponding to these core motifs and their respective positive and negative control sets.

### *De novo* motif discovery from EdU-seq origins

The MEME suite software package was utilized for *de novo* motif discovery and generation. Shuffled control sequences were generated from the input sequences using a 2^nd^ order Markov model (fasta-shuffle-letters -m 2) for use in differential enrichment motif discovery, which was carried out to search for 20-30 bp motifs with any number of occurrences within sequences (meme -objfun de -dna -revcomp -mod anr -minw 20 -maxw 30 -nmotif 5). The MEME-ChIP and CentriMo subpackages were used to confirm the reproducible nature of the motif and enrichment of it near the center of peak calls. Motif scanning and scoring (both for regions under peaks as well as genome-wide) was done using FIMO, using *p*<1e-4 as a cutoff to find matches and p<1e-3 for broad scouting of similar sequences using the calculated p-value as a scoring function. Figures for motif discovery/search were generated either using the native output from MEME, or with the assistance of Claude Science (Anthropic. Claude Opus 4.8 (Claude Science 0.1.22) [Large language model]. https://claude.com/product/claude-science (2026).

### Expression and Purification of Human ORC Subunits (HsORC1–5)

Codon-optimized *HsORC1* (NP_004144.2), N-terminally fused with twin Strep and SUMO tags, was cloned into the pFL vector for expression in insect cells. The remaining synthetic human ORC genes—*HsORC2* (NP_006181.1), *HsORC3* (NP_862820.1), *HsORC4* (NP_859525.1), and *HsORC5* (NP_002544.1)—were cloned into the MultiBac baculovirus expression system ^99^. A twin StrepTag followed by a TEV cleavage site was also introduced at the N-terminus of *HsORC3* to facilitate affinity purification. All HsORC proteins are full length. Recombinant expression of HsORC1 and separately of the HsORC2-5 complex were performed in Sf9 insect cells infected with baculovirus and cultured in CCM3 medium (GE Healthcare Life Sciences, Pittsburgh, PA) for 48 hours. Cell pellets for both *HsORC1* and *HsORC2-5* were resuspended separately in lysis buffer containing 50 mM HEPES-NaOH (pH 7.5), 300 mM KCl, 30 mM potassium glutamate, 5 mM magnesium acetate, 5 mM dithiothreitol (DTT), and 2 mM ATP. HsORC1-expressing cells were lysed by sonication, and lysates were clarified by centrifugation at 143,000 × *g* for 45 minutes. The supernatant was loaded onto a 5 mL StrepTactin agarose beads onto a gravity flow column. After washing, bound *HsORC1* protein was eluted with lysis buffer supplemented with 5 mM desthiobiotin. The HsORC2-5 complex was purified in parallel using an identical Strep-tag affinity protocol. Purified HsORC1 and HsORC2-5 protein fractions were combined, and TEV protease was added for tag cleavage, followed by incubation at 4 °C for 12 hours. The mixture was subsequently diluted to 150 mM KCl and subjected to ion exchange chromatography using a HiTrap SP column with a linear gradient from 150 to 1000 mM KCl. Protein-containing fractions were analyzed by SDS-PAGE. Fractions containing all five ORC subunits were pooled, concentrated, and further purified by size exclusion chromatography using a Superose 6 Increase 10/300 GL column (GE Healthcare) equilibrated with minimal buffer (25 mM HEPES-NaOH (pH 7.5), 100 mM KCl, 2 mM DTT). Final protein purity was assessed by SDS-PAGE, and pure fractions were concentrated using an Amicon® Ultra centrifugal filter (50 kDa MWCO). Aliquots of ∼3–5 μM were snap-frozen in liquid nitrogen and stored at –80 °C.

### Expression and Purification of Human CDC6 Protein

The human CDC6 (HsCDC6) gene was cloned into the pET28a vector to allow for IPTG inducible expression in *Escherichia coli*. The construct encoded full-length HsCDC6 with an N-terminal His-SUMO tag. The resulting plasmid was transformed into *E. coli* Rosetta (DE3) cells and cultured in 4 liters of Terrific Broth (TB) supplemented with kanamycin. Cells were grown at 37°C until reaching an optical density (OD₆₀₀) of 0.8–1.0, at which point expression was induced with 0.5 mM IPTG. Protein expression was carried out for 16 hours at 16°C, and cells were harvested by centrifugation at 3,500 × g for 15 minutes. Cell pellets were resuspended in lysis buffer (50 mM HEPES, pH 7.0, 300 mM NaCl, 10 mM imidazole, 2 mM β-mercaptoethanol) and lysed by sonication on ice. The lysate was clarified by centrifugation and loaded onto a gravity-flow column packed with Ni-NTA agarose beads and pre-equilibrated with lysis buffer. The column was washed with 10 column volumes (CV) of lysis buffer, followed by 5 CV of high-salt buffer (lysis buffer with 500 mM NaCl). Bound protein was eluted using lysis buffer containing 400 mM imidazole. Eluted protein was incubated with TEV protease overnight at 4°C to cleave the His-SUMO-TEV tag. The cleaved sample was diluted and applied to a HiTrap SP column (GE Healthcare) equilibrated in buffer (50 mM HEPES-NaOH, pH 7.0, 150 mM NaCl, 1 mM DTT) for ion exchange chromatography. Protein was eluted with a linear gradient of Buffer B (50 mM HEPES-NaOH, pH 7.0, 1 M NaCl, 1 mM DTT). Protein peak fractions were pooled and subjected to size exclusion chromatography (SEC) using a Superdex 200 Increase 10/300 GL column, equilibrated with SEC buffer (25 mM HEPES-NaOH, pH 7.5, 100 mM KCl, 2 mM DTT). Protein purity was assessed by SDS-PAGE, and pure fractions were concentrated using an Amicon® Ultra centrifugal filter (30 kDa MWCO). Final protein aliquots at concentrations of approximately 10–15 μM were snap-frozen in liquid nitrogen and stored at –80°C.

### Protein purification and preparation of YlODC

Synthetic, full-length genes of Yarrowia lipolytica *ORC1* (*YlORC1*) (RefSeq: XP_502645.1), *YlORC2* (XP_503147.3), *YlORC3* (XP_505428.1), *YlORC4* (XP_504002.3), *YlORC5* (XP_500387.1), and *YlORC6* (XP_506105.1) were codon optimized and cloned into either pFL (pH promoter *YlORC1*, p10 promoter *YlORC6*), pSPL (pH promoter *YlORC2*, p10 promoter *YlORC5*), or pUCDM (pH promoter *YlORC4*, p10 promoter *YlORC3*) plasmids for expression via the MultiBac baculovirus expression system ^99^. To increase protein expression and solubility, in addition to providing a tag for affinity chromatography, an N-terminal Twin Strep-SumoStar-TEV tag was added to YlORC1. The tagged version of YlOrc1 was utilized for the Cdc6 loading assay and EM studies. *YlCDC6* (XP_501295.1) was codon optimized and cloned into a pET28b bacterial expression cassette (Novagen) containing an N-terminal 8xHis-TEV tag for affinity purification.

For YlORC, Sf9 insect cells were incubated with baculovirus for 72 hr in Hyclone CCM3 media (GE Healthcare Life Sciences, Pittsburg, PA). For YlCdc6 expression, BL21 (DE3) strain *E. coli* were transformed and then selected for with a kanamycin-supplemented LB starter culture, followed by inoculation of kanamycin-supplemented Terrific Broth (TB). Expression was induced at OD 1 with the addition of 0.5 mM IPTG and incubated at 17°C overnight.

Unless otherwise noted, each purification step was carried out at 4°C. For YlORC purification, insect cell pellets were thawed in a 30°C water bath, resuspended in lysis buffer (50 mM HEPES-NaOH (pH 7.5), 150 mM potassium acetate (KOAc) (pH 7.5), 50 mM potassium glutamate (K-Glu), 50 mM arginine hydrochloride, 10 mM Mg(OAc)_2_, 6.5 mM dithiothreitol (DTT), 1.65 mM adenosine triphosphate (ATP), 10% glycerol), supplemented with a protease inhibitor cocktail (1 mM PMSF, 2 µM pepstatin, 2 µM leupeptin, 1 mM benzamidine, 1:1725 dilution of Millipore-Sigma aprotinin (A6279)) in addition to 1X cOmplete EDTA-free protease inhibitor cocktail (Roche) and sonicated. Lysate was then centrifuged at 38,000g for 1 hour, after which the supernatant was collected and 3 mL of either StrepTactin Superflow resin or StrepTactin 4Flow resin was added to the supernatant and incubated for 90 minutes. The resin was washed and YlORC was eluted using lysis buffer supplemented with 5 mM desthiobiotin. YlORC-containing fractions were pooled together, and λ-phosphatase was added at a ∼2:1 YlORC:phosphatase molar ratio, along with 1 mM manganese chloride, and incubated at 4°C for 36-48 hours. In preparations used for biochemical assays, YlORC was simultaneously treated with TEV protease at an YlORC:TEV mass ratio of 15:1. The phosphatase-treated elution then underwent anion exchange chromatography (HiTrap Q HP 5 mL, Cytiva) followed by size exclusion chromatography (Superose 6 increase 10/300 GL, Cytiva) equilibrated in minimal buffer (25 mM HEPES-NaOH (pH 7.5), 100 mM KOAc (pH 7.5), 50 mM K-Glu, 5 mM Mg(OAc)_2_, 1 mM DTT, 5% glycerol). Aliquots were made with YlORC concentrated to ≍2.4 mg/mL.

For YlCdc6 purification, cell pellets were thawed in similar conditions except that the lysis buffer contained an additional 10 mM imidazole. For YlCdc6 affinity purification, 5 mL of Ni-NTA agarose was added to clarified lysate, with resin washes done with 25 mM imidazole supplemented lysis buffer, and eluted with 50 mM, 100 mM, 250 mM, and 500 mM imidazole supplemented lysis buffer. TEV protease was added and incubated overnight at 4°. YlCdc6 was further purified using cation exchange chromatography (HiTrap SP HP 5 mL, Cytiva) followed by size exclusion chromatography (Superdex 200 increase 10/300 GL, Cytiva) into minimal buffer (25 mM HEPES-NaOH (pH 7.5), 100 mM NaCl, 1 mM DTT). Aliquots were made by supplementation with glycerol to 5%, and YlCdc6 was concentrated to 5 mg/mL.

### Synthetic *ori* oligonucleotides for structural analysis

Oligonucleotides were ordered from IDT and were then annealed to their complementary oligonucleotides by incubating them together at 95°C for 5 minutes, followed by a temperature decrease of 2°C/min until 25°C was reached. The ordered oligonucleotides are displayed in the table below:

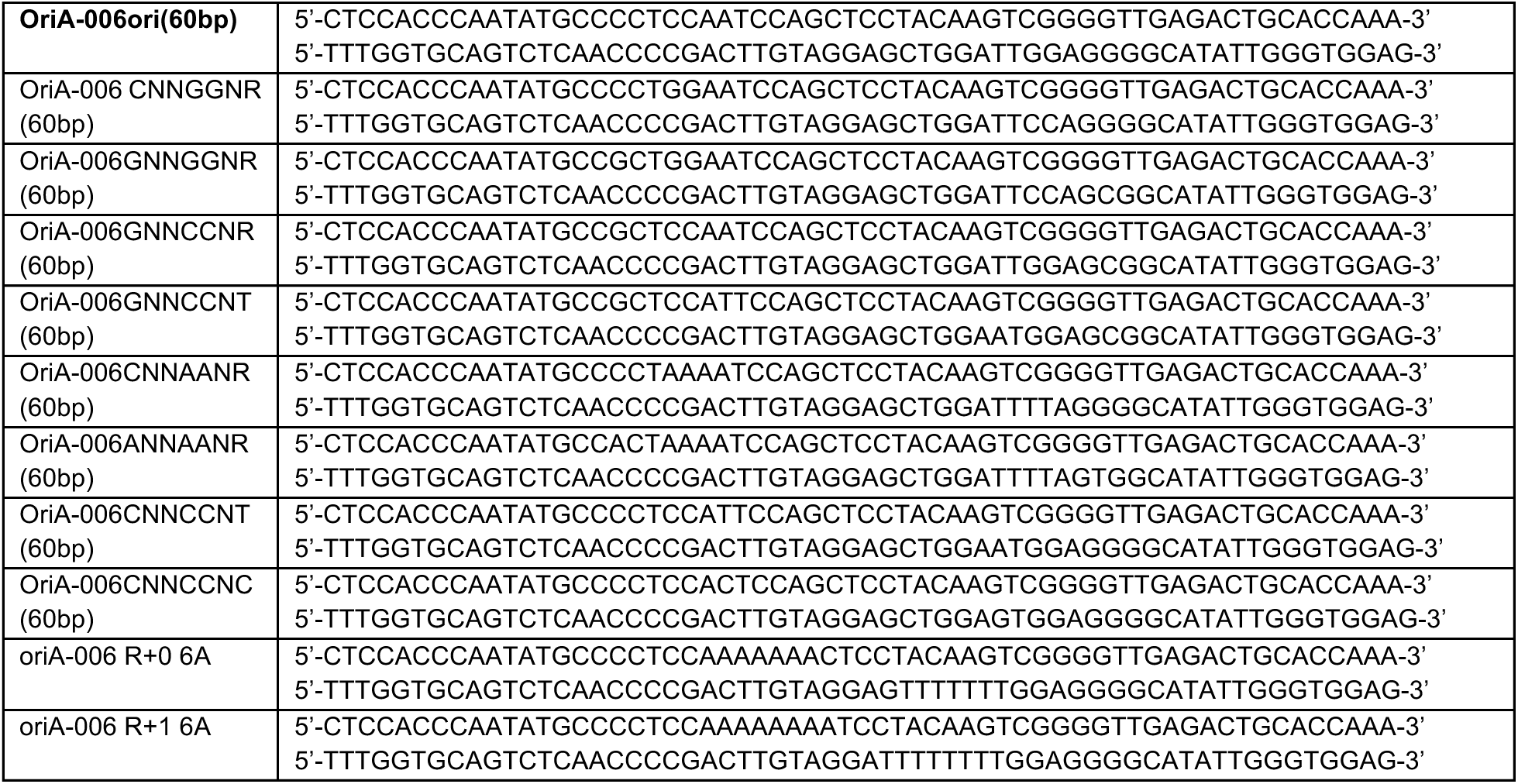
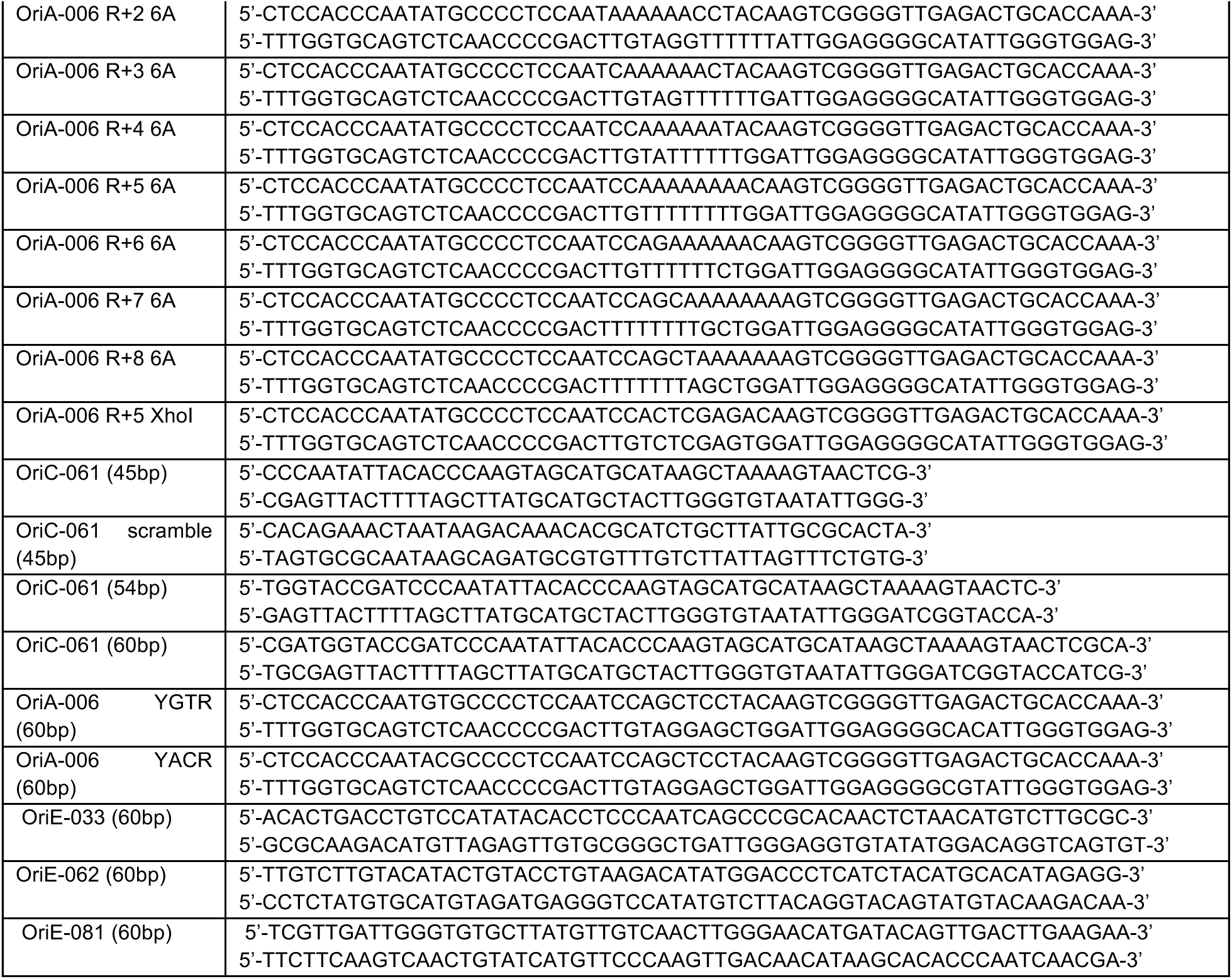

### Cdc6 loading assay

To test the dependence of YlORC’s ability to bind DNA and load YlCdc6 on the origin, annealed oligonucleotides representing *ori* variants and 5X reaction buffer (750 mM KOAc (pH 7.5), 250 mM HEPES-NaOH (pH 7.5), 50 mM Mg(OAc)_2_, 5 mM DTT, 5 mM ATP, 50% glycerol) were added to purified YlORC1-6, with an ORC:DNA molar ratio of 1:1.5, and allowed to incubate at room temperature for 10 minutes. For YlCdc6-containing assays, purified YlCdc6 was added following the ORC-DNA incubation at an ORC:Cdc6 molar ratio of 1:4 and was incubated at room temperature for 10 minutes. Samples were then loaded onto a Superose 6 increase 3.2/300 microkit column (Cytiva) equilibrated in SEC buffer (25 mM HEPES-NaOH (pH 7.5), 100 mM NaCl, 1 mM DTT) and fraction samples were run on SDS-PAGE and visualized using ReadyBlue Protein Gel Stain (Sigma). For experiments determining the total quantity of DNA in each fraction, SDS-PAGE gels were stained with SYBR Gold and imaged prior to ReadyBlue staining. 260/280nm UV absorbance ratios obtained from chromatograms were also used for qualitative analysis of DNA and Cdc6 binding to ORC.

Gel images were quantified using the ImageJ-based software package Fiji ^100^. For calculating relative loading efficiency, the intensity ratio of the YlCdc6 band to each ORC subunit band was normalized to the intensity ratios observed while running the assay with a 60bp fragment of the WT *OriA-006* and averaged. All images of the gels used for figures and quantification data are available in a Mendeley Data Repository (10.17632/zd96z6h9c5.1).

### Cryo-EM sample preparation

**HsODC:** Purified HsORC1–5, HsCDC6, DNA and AMP-PNP were mixed at final concentrations of 2.5 μM, 5.0 μM, 7.5 μM and 10 μM, respectively. To reduce preferred particle orientations and promote the formation of thin ice layers over grid holes, lauryl maltose neopentyl glycol (LMNG; Anatrace, Maumee, OH) was added to a final concentration of 0.05% (w/v). For cryo-electron microscopy, 4 μL of the protein–DNA complex was applied to a glow-discharged Quantifoil R 0.6/1, 300 mesh copper grid. The grid was incubated for 10 seconds at 25°C and 90% humidity, blotted for 3.0 seconds, and then rapidly plunge-frozen into liquid ethane using a Leica EM GP2 automatic plunge freezer (Leica Microsystems, Buffalo Grove, IL).

**YlODC^60bp*Ori-A006*–WT^**: a similar protocol was used with key differences. A 60 bp *OriA-006* fragment was used instead of the 45 bp *OriC-061* fragment, and the final protein concentration was 1 mg/mL. The sample was applied to a glow-discharged lacey carbon grid and blotted for 3.0 seconds.

**YlODC^54bp*OriC-061*^**: purified YlORC1-6 was mixed with glycerol-free 5X loading assay reaction buffer, a 54 bp *OriC-061* fragment, and YlCdc6 at an ORC:DNA:Cdc6 molar ratio of 1:1.5:4 in a stepwise fashion, followed by gel filtration using a Superose 6 increase 3.2/300 microkit column (Cytiva), identical to the Cdc6 loading assay. Fractions containing the YlORC-DNA-Cdc6 complex were then concentrated with a 0.5 mL Amicon Ultra 50kDa MWCO centrifugal filter to a final protein concentration of 1-1.25 mg/mL without additional ATP supplementation. Following concentration, lauryl maltose neopentyl glycol (LMNG) was added to 0.05% to reduce a preferred orientation issue. 4 µL of sample were applied to a non-glow discharged Quantifoil R 1.2/1.3 300 mesh copper grid (previously washed with ethyl acetate), incubated for 10 seconds at 25°C and 95% humidity, blotted for 2.9 seconds, and plunged into liquid ethane using a Leica Automatic Plunge Freezer EM GP2.

**YlODC^60bp*Ori-A006*–CNNGGNR^**: sample preparation followed the protocol for the YlODC^45bp*OriC-061*^ sample, with key differences. A 60 bp *OriA-006* fragment containing the CNNGGNR mutation (see oligo table for sequence) was substituted, and samples had a final protein concentration of

1.6 mg/mL. Additionally, a non-glow discharged Quantifoil R 1.2/1.3 300 mesh copper grid (previously washed with ethyl acetate) was used and was blotted for 2.7 seconds.

### Cryo-EM data acquisition

**HsODC:** Cryo-EM data were collected on a Titan Krios transmission electron microscope (ThermoFisher Scientific) operating at 300 keV. Data were acquired using EPU software (v2.10.0.5, ThermoFisher Scientific), and dose-fractionated movies were recorded on a K3 direct electron detector (Gatan) in electron counting mode. HsODC samples were applied to Quantifoil R 0.6/1 grids, and 30-frame movies were collected at an exposure rate of 1.44 e⁻/Å²/frame, yielding a cumulative dose of 43.2 e⁻/Å². A total of 7088 micrographs were acquired at 81,000× nominal magnification, with a defocus range of 0.6–2.2 μm.

**YlODC**: Cryo-electron microscopy data were collected using an FEI/ThermoFisher Titan Krios TEM operating at 300 keV. A Gatan K3 direct electron detector equipped with a BioQuantum energy filter was utilized to semi-automatically collect dose-fractionated movies with ThermoFisher EPU data collection software. For the YlODC^45bp*OriC-061*^ maps, two collections on consecutive days accrued 8978 and 9310 exposures, respectively, with movies collected with 30 frames at a dose rate of 1.98 e/Å^2^ per frame, resulting in a cumulative dose of 59.4 e/Å^2^. For the YlODC^54bp*OriC-061*^ data collection, 30-frame movies were collected over three consecutive days, resulting in 9309, 8758, and 2274 exposures taken, respectively, at a dose rate of 1.44 e/Å^2^ per frame, totaling a cumulative dose of 43.2 e/Å^2^. For the YlODC^60bp*Ori-A006*–WT^ data collection, a single session was used to collect 8340 exposures, with movies containing 40 frames at a dose rate of 1.97 e/Å^2^ per frame, totaling 78.8 e/Å^2^ in cumulative dose. YlODC^60bp*Ori-A006*–CNNGGNR^ data collection included 8428 exposures from a single session, with 40 frames per movie, a dose rate of 1.37 e/Å^2^ per frame, and a cumulative dose of 54.8 e/Å^2^.

### Cryo-EM data processing

**HsODC**: Real-time preprocessing, including motion correction, CTF estimation, and particle picking, was performed in WARP (v1.0.9). Particle picking used the BoxNet pretrained neural network implemented in TensorFlow, with a particle diameter of 180 Å and a threshold score of 0.6, resulting in 898,455 coordinates. Subsequent image processing was carried out in cryoSPARC v3.2. Particles were extracted and subjected to multiple rounds of 2D classification, and well-resolved subsets were selected for *ab initio* 3D reconstruction. Separation into 3–5 *ab initio* classes proved critical for improving map quality. These models were used for 3D heterogeneous refinement against the full dataset, yielding 443,190 selected particles for HsODC. This subset was further classified into three classes, and the best class was refined. Homogeneous and non-uniform refinements for the best 3D class (130,819 particles) produced a cryo-EM map at **2.6 Å resolution**, as determined by the gold-standard FSC (GSFSC) criterion. The final sharpened map was used for model building and visualization (**Figure S1B**).

**YlODC:** WARP was utilized for motion correction, CTF estimation, and particle picking (via BoxNet neural network trained on manually picked micrographs) from the collected micrographs for all datasets ^101^. For particle picking, a particle diameter of either 180 Å or 200 Å and a thresholding score of either 0.3, 0.4, or 0.5 were used, yielding 1,822,720 particles for YlODC^45bp*OriC-061*^, 3,805,196 particles for YlODC^54bp*OriC-061*^, and 793,509 particles for YlODC^60bp*Ori-*^ *^A^*^006^–^WT^. While WARP pre-processing and picking was carried out for the YlODC^60bp*Ori-A006*–CNNGGNR^ exposures originally, cryoSPARC’s pre-processing and particle picking tools were used instead, described below. All downstream processing was carried out using cryoSPARC v4 ^102–104^. All 2D classifications underwent an extra final iteration, all heterogeneous refinements listed used particles binned to 128 pixels, and all non-uniform refinements used “minimize over per-particle scale” and underwent one extra final pass.

**YlODC^45bp*OriC-061*^**: 1,822,720 particles were picked and extracted by WARP/BoxNet unbinned with a box size of 480 pixels and imported into cryoSPARC for processing. Each day of collection underwent its own set of 2D classification, resulting in two 2D classification jobs, one with 970,731 starting particles and 194,059 particles selected for further processing, and another with 851,989 particles with 457,613 selected. The 194,059 particles were used for *ab-initio* model generation of three classes. The generated *ab-initio* structures were then used in a heterogeneous refinement that included the entire particle dataset (1,822,720 particles), resulting in 802,503 particles constituting the best class. This class then underwent homogenous refinement followed by non-uniform refinement, resulting in a 2.9 Å resolution map. Particles from this map underwent another round of 2D classification, selecting for 533,196 particles. Another heterogeneous refinement was carried out, using the *ab-initio* models as templates, resulting in a class with 302,465 particles. This class was then subjected to non-uniform refinement with simultaneous per-particle scale minimization, per-particle defocus refinement, and CTF refinement of per-group CTF parameters, spherical aberration, and tetrafoil, producing a 2.7 Å map. Finally, 3D classification was carried out, and the class containing the strongest Orc2-WHD was selected and underwent a final non-uniform refinement, resulting in a final map with a resolution of 2.7 Å from 84,914 particles

**YlODC^54bp*OriC-061*^**: 3,805,196 particles were picked and extracted by WARP/BoxNet unbinned with a box size of 440 pixels and imported into cryoSPARC for processing. Following a check for corrupt particles, each day’s full particle set was used in ab-initio model generation to generate 4 classes. The best class of each set was used in non-uniform refinement with per-particle scale minimization, per-particle defocus refinement, and CTF refinement of per-group CTF parameters, spherical aberration, and tetrafoil to produce 3 maps (1,512,226 particles → 538,187 particles, 2.6 Å resolution map; 509,844 particles → 210,193 particles, 2.6 Å resolution map; 1,377,300 particles → 488,199 particles, 2.5 Å resolution map) to validate pixel size and spherical aberration parameters. 2D classification with 250 classes was then carried out on the entire dataset, from which most (3,705,630) particles were selected for ab-initio model generation commenced from 3,705,630 particles sorted into eight classes, the best of which was then flipped for proper handedness. Heterogeneous refinement was then carried out, producing a 2.3 Å resolution map from 1,453,157 particles that appeared heterogeneous in density. 3D classification was carried out with 10 classes at a filter resolution of 8 Å, which were then used to produce 6 maps via non-uniform refinement as the result of combining some 3D classification output classes into one structure. One of these structures underwent a further 3D classification into 2 classes at a filter resolution of 10 Å, selecting for the class that contained the Orc1-AAA domain while bound to DNA. Following this, a large-scale heterogeneous refinement of the entire dataset was done again, this time split into 14 classes: the aforementioned Orc1-AAA containing map, three maps derived from the 3D classification, the YlODC^45bpOriC-061^ map, and a map of YlORC-DNA produced from a previous collection. Following this large-scale classification/refinement, a YlODC^54bpOriC-061^ map at 2.5 Å resolution was produced with weak Cdc6 density from 558,557 particles. This class then underwent 3 cycles of 3D classification and non-uniform refinement with per-particle scale minimization, per-particle defocus refinement, CTF refinement of per-group CTF parameters, spherical aberration, and tetrafoil refinement. Each 3D classification consisted of only 2 classes and used a filter resolution of 10 Å. After the 3 rounds of refinement and re-classification, a 2.7 Å map from 70,712 particles was produced. Following this, the Subset Particles job was used to select particles by per-particle scale, leaving 51,599 particles for use in another non-uniform refinement, producing a 2.7 Å resolution map to be used as the final YlODC^54bpOriC-061^ map. A schematic summary of data processing is shown in **Figure S4B**.

**YlODC^60bp*Ori-A006*–WT^**: 793,509 particles were picked and extracted by WARP/BoxNet unbinned with a box size of 440 pixels and imported into cryoSPARC for processing. Following a check for corrupt particles, 2D classification into 200 classes was done, with 416,680 particles selected for *ab-initio* model generation of 6 maps. The YlODC^54bp*OriC-061*^ map with weak Cdc6 density was imported into the project and heterogeneous refinement with the six *ab-initio* maps and the imported YlODC^54bp*OriC-061*^ map was carried out on the entire particle dataset. The best heterogeneous refinement volume/particles underwent non-uniform refinement with the same scale, defocus, and CTF refinement corrections done in previously mentioned non-uniform refinements, and a 2.8 Å map from 255,191 particles was produced. Particles used in this map were then repicked from motion- and CTF-corrected micrographs (carried out in cryoSPARC) to generate a 2.9 Å resolution map, which then underwent Reference-Based Motion Correction to produce a 2.7 Å resolution map from 246,739 particles. 3D classification at a filter resolution of 10 Å into 3 classes were performed, and the best two classes were combined for another round of non-uniform refinement, followed by a Subset Particles job (by per-particle scale) and a final non-uniform refinement to produce a 2.6 Å resolution map from 125,267 particles, which was then used for model building of YlODC^60bp*Ori-A006*–WT^. A schematic summary of data processing is shown in **Figure S4K**.

**YlODC^60bp*Ori-A006*–CNNGGNR^**: 8,428 movies were imported into cryoSPARC and underwent patch motion correction and patch CTF corrections to generate corrected micrographs. Template picking of micrographs commenced using representative 2D class averages of the particles used in the final YlODC^60bp*Ori-A006*–WT^ refinement, resulting in 4,708,382 particles being picked. Micrographs and respective particles were then analyzed using the Micrograph Junk Detector job, and after exposure and particle curation resulted in 2,710,398 particles from 7,330 micrographs. Particles were then extracted with a box size of 432 px and Fourier cropped to 128 px. Initial 2D classification utilized 200 classes, of which 2,4,69,383 particles from 184 classes were selected for use in further processing. *Ab-initio* model generation of 8 maps from 400,000 particles failed to produce a YlODC structure, so the YlODC^60bp*Ori-A006*–WT^ map was imported into the project, and heterogeneous refinement with the eight *ab-initio* maps along with the imported YlODC^60bp*Ori-A006*^– ^WT^ map was performed on the particle dataset. The best heterogeneous refinement volume/class (481,746 particles) underwent non-uniform refinement and further 2D classification to produce a cleaned stack of particles (305,326 particles) for particle re-extraction without Fourier cropping. Non-uniform refinement of the extracted particles produced a 2.9 Å resolution map, which was used as the reference volume for Reference-Based Motion Correction and re-refined to produce a 2.6 Å resolution map. Particles were then subset by per-particle scale and re-refined, generating a 2.5 Å resolution map of the complex with variable Cdc6 density. 3D classification into three classes generated a class from 51,222 particles containing the full YlODC complex, which was used for a final non-uniform refinement to produce the final 2.56 Å resolution map of YlODC^60bp*Ori-*^ *^A^*^006^–^CNNGGNR^ used for model building. All non-uniform refinements of the unbinned particles utilized per-particle scale minimization, per-particle defocus refinement, and CTF refinement of per-group CTF parameters, spherical aberration, and tetrafoil. A schematic summary of data processing is shown in **Figure S5D**.

### Model building and validation

**HsODC:** The atomic model of HsORC (PDB ID: 7JPS) was used as the starting model for HsODC and rigid-body fitted into the cryo-EM density using ChimeraX. Regions of HsORC that were missing or did not fit well into the density were rebuilt manually in Coot. Iterative model building and refinement were performed in PHENIX (v1.20.1–4487-000), with secondary-structure restraints applied throughout. Model validation was carried out using MolProbity and PHENIX validation tools. The final model showed good stereochemistry, with >95% of residues in favored regions of the Ramachandran plot, <0.5% outliers, and all bond length and bond angle deviations within acceptable limits. Structural figures were generated using ChimeraX and PyMOL (v2.5.5, Schrödinger, LLC).

**YlODC**: For the YlODC^45bp*OriC-061*^ structure, AlphaFold 2 models for each subunit were docked into the density individually using the “fit to map” functionality in ChimeraX ^105^, then refined using the Coot software package ^106^. The density for the DNA was sharp enough to allow us to discern purines and pyrimidines, allowing us to produce a generic DNA-B form model of the respective DNA sequence and manually rebuild it in Coot. This structure was then used as the basis of the other ODC structures. Structures were then further refined in Coot using “Real Space Refinement” function and ligands, ions, and waters were manually built. The Phenix software package was then used to further refine and finalize the structures, as well as provide validation metrics, via its “Real Space Refine” functionality ^107^. Figures using these structures, along with comparisons to previously published ORC/ODC structures, were generated using ChimeraX. The preliminary YlODC^45bp*OriC-061*^ structure was then used as the basis for the YlODC^54bp*OriC-061*^ structure, which itself was used as the starting point for the YlODC^60bp*Ori-A006*–WT^ structure. Additionally, the YlODC^60bp*Ori-A006*–WT^ structure was used as a reference for building the YlODC^60bp*Ori-A006*–CNNGGNR^ structure.

---

**Figure S1 (related to Figure 1).**
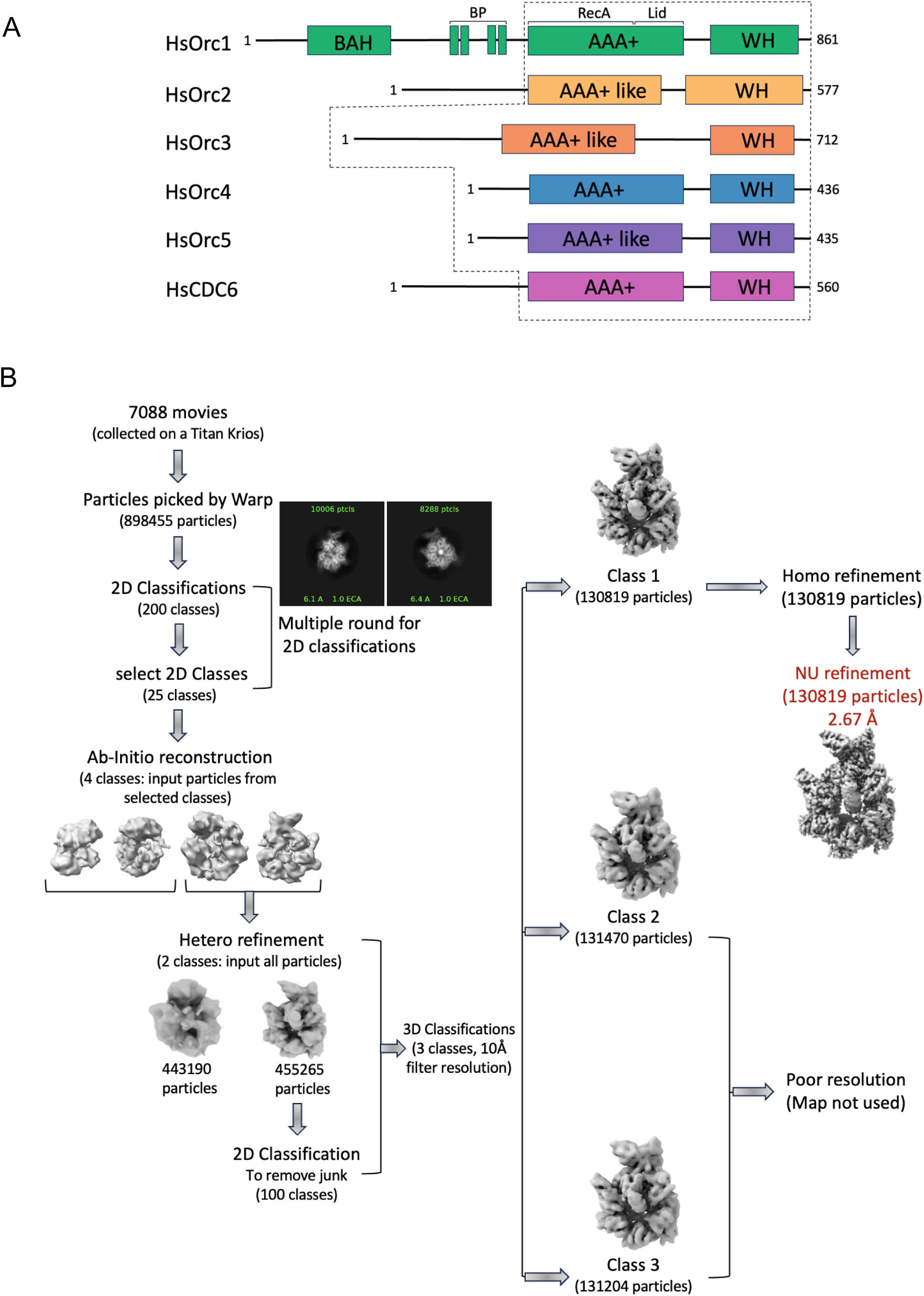
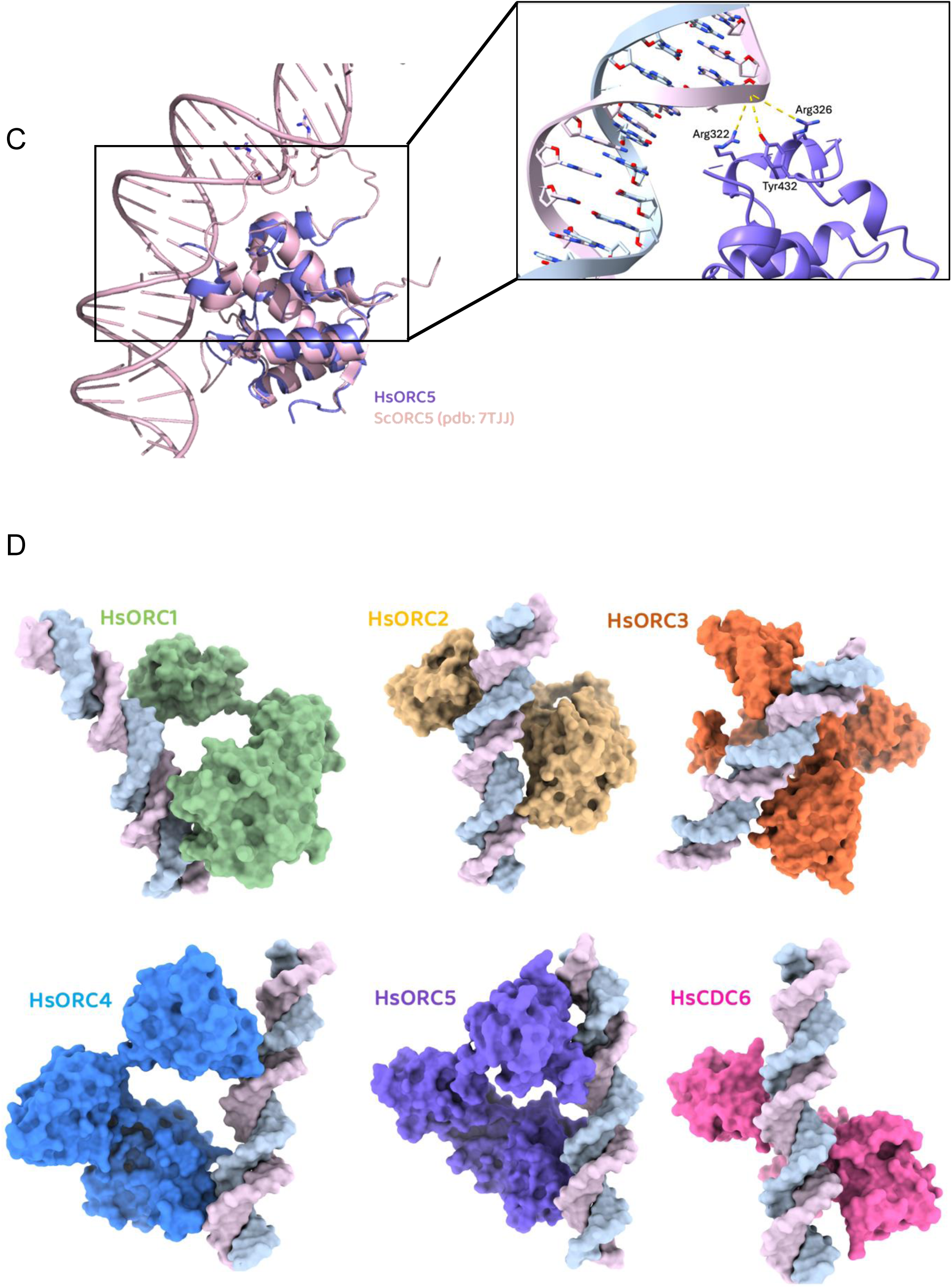
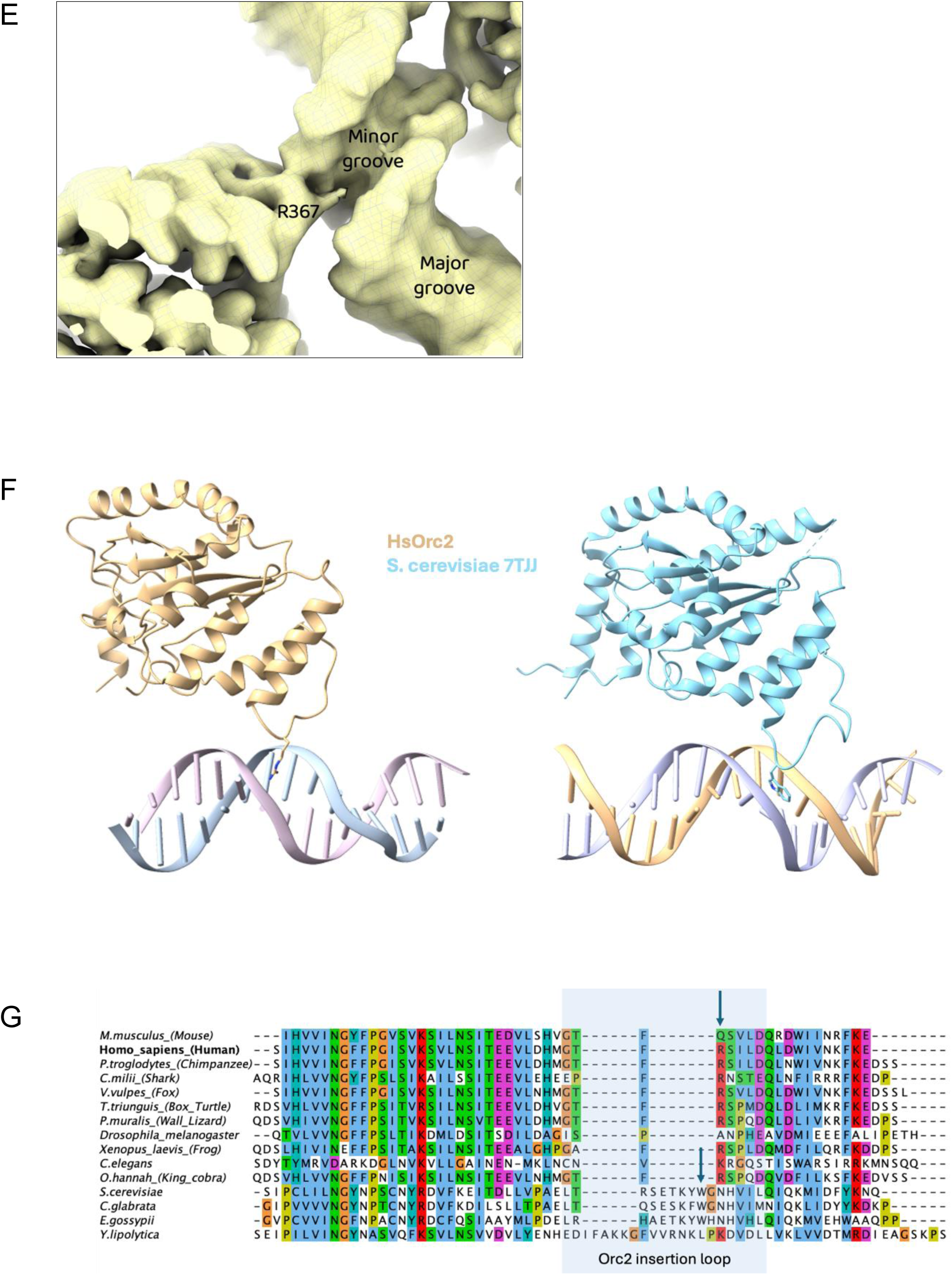
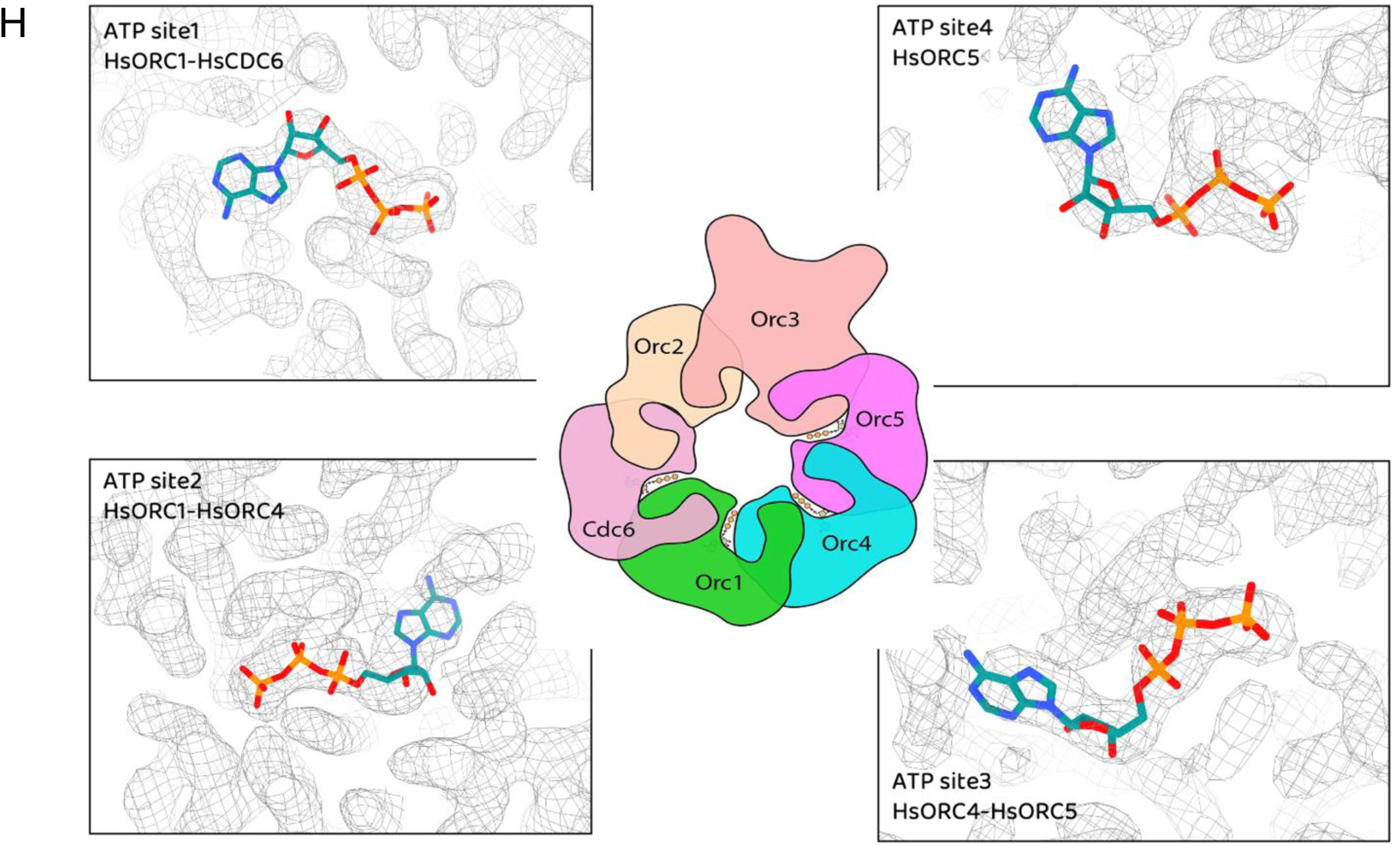
(A) Domain organization and subunit–DNA interactions in the HsORC–Cdc6 complex. Schematic representation of the domain organization of all HsORC subunits (Orc1–5) and HsCdc6. Protein regions that were resolved and modeled in the cryo-EM structure are outlined with dashed gray lines. **(B)** Cryo-EM workflow of HsODC structure. Motion correction, averaging, CTF estimation, and particle picking steps were performed on movies in real time via the WARP (v1.0.9) software package. To determine an *ab initio* reconstruction, 2D classification and all further processing steps were performed in Cryosparc v3.2. See Methods section for details. **(C)** Structure alignment of the human ORC5 subunit with the *S. cerevisiae* DNA-bound ORC5. Alignment shows that *S. cerevisiae* ORC5 contains an extended, flexible loop enriched in basic residues that engages DNA and supports pronounced DNA bending, whereas the corresponding region in human ORC5 is disordered, resulting in DNA contacts limited to the basic patch residues shown in stick representation on the right. **(D)** Structural views of individual protein subunits interacting with DNA, derived from the cryo-EM density map. Figures were generated using ChimeraX. **(E)** Close-up view of the density map around residue R367 from the HsORC2 subunit. R367 is observed making direct contacts with DNA bases within the minor groove. **(F)** Side-by side structures of HsORC2 (yellow, this work) and ScOrc2 (PDBID 7TJJ) (cyan) with R367 from human and W396 of *S. cerevisiae* in stick representation. **(G)** Sequence alignment of the region surrounding R367. Arrows indicate conservation of R367 and the equivalent residue, W396, in *S. cerevisiae* that contacts DNA specifically. **(H)** Cryo-EM density surrounding the ATP molecules at all four conserved ATP-binding sites within the HsORC–Cdc6 complex. The density is well-resolved, allowing precise placement of ATP and modeling of coordinating residues.

**Figure S2 (related to Figure 2).**
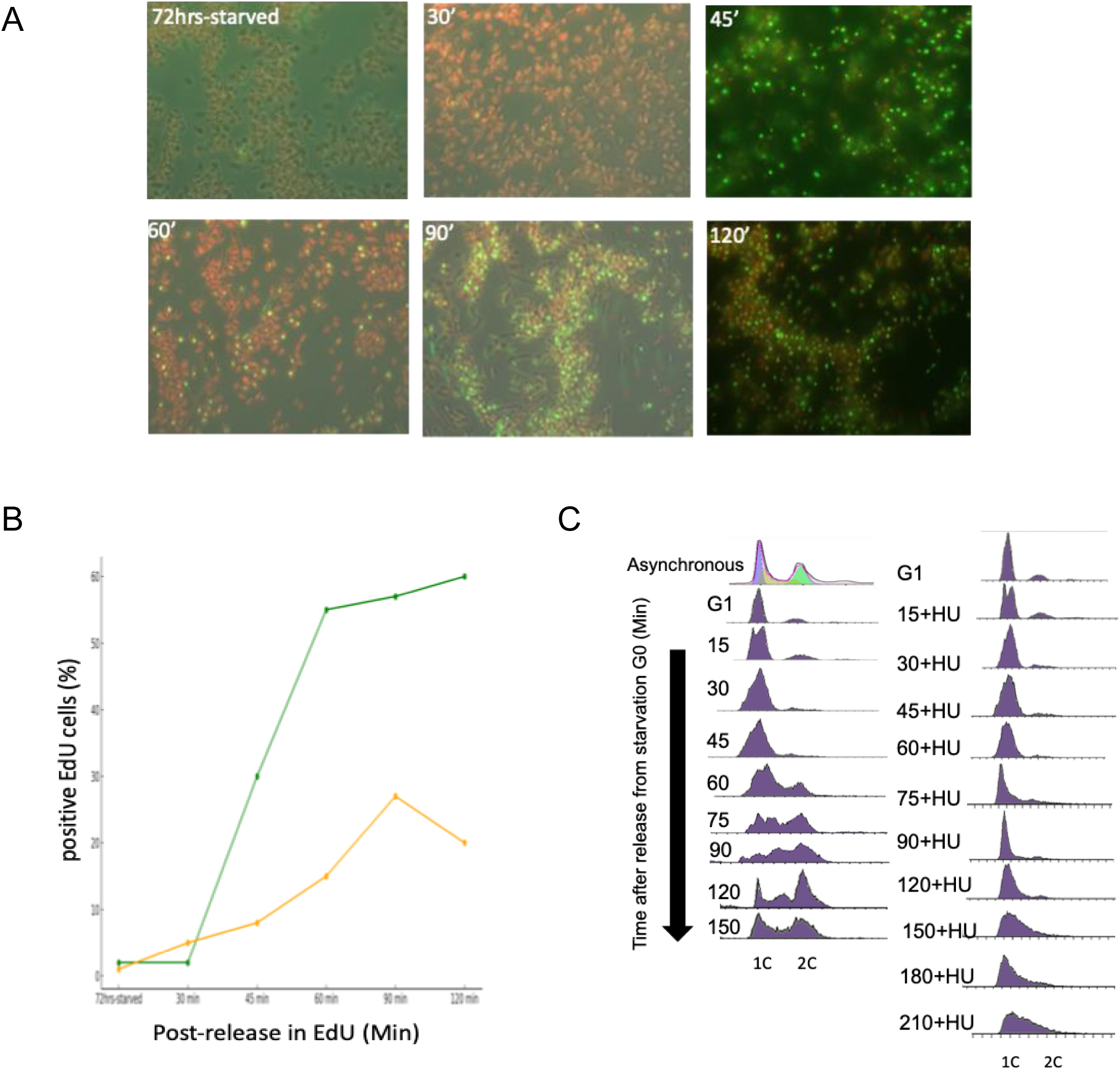
Time-Course Analysis of S-Phase Progression in the Tk+ Strain Using EdU Incorporation. **(A)** Fluorescence microscopy images cells either starved for released into fresh, complete media for various times were treated with 100 µM EdU and DNA synthesis was visualized using Click-iT™ chemistry with Alexa Fluor™ 488 (green fluorescence). Red fluorescence (DAPI) was used to stain total nuclear DNA for cell visualization. **(B)** The percentage of EdU-positive cells at each time point (Green line) and percentage of budded cells (orange line). **(C)** flow cytometry of DNA content in asynchronously growing cells, cells arrested in G1 by starvation fro 72 hours and cells released for different times in the absence (left) or presence (right) of 5 mM HU.

**Figure S3 (related to Figure 3).**
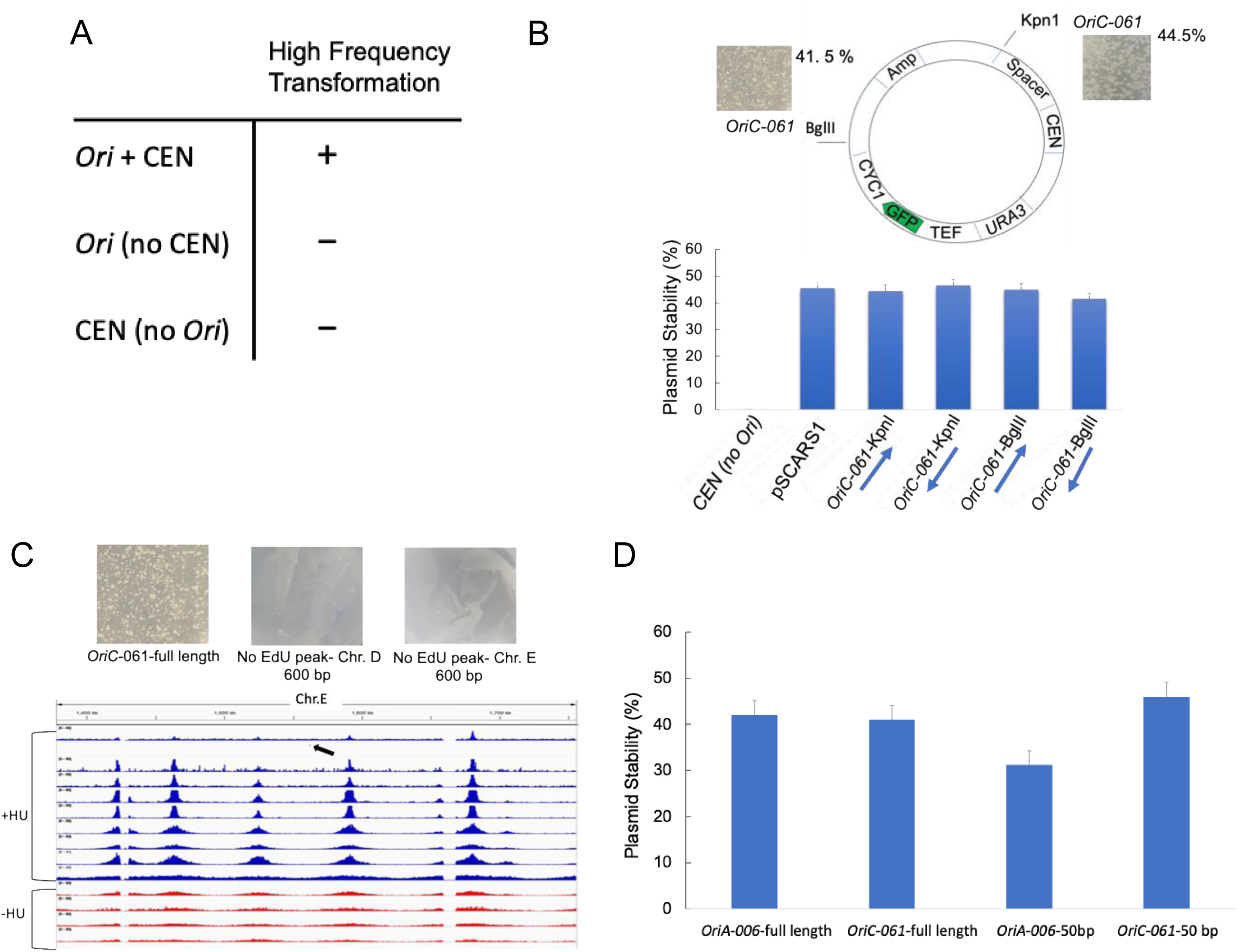
Plasmid assay for origin activity. **(A)** in *Y. lipolytica*, autonomous replication of a plasmid required both a centromere (*CEN*) and origin of DNA replication (*Ori)* Vernis et al. DOI: 10.1128/mcb.17.4.1995. The plasmid has the *URA3* selectable marker. **(B)** Plasmid construct showing a CEN, and two *Ori* insertions sites at either Kpn1 of BglII. When *OriC-061* was inserted at either site, high frequency transformation occurred and the plasmid stability was comparable. **(C)** The *OriC-061* origin, but not a 600 bp form Chromosome D or Chromosome E (arrow showing region between two origins) show high frequency transformation upon *URA3* selection. The *OriC-061* plasmid is the same as shown in panel B. **(D)** 50 base pair region containing the core 30 base pair region of *OriA-006* or *OriC-061* function as efficiently as a full length 600 base pair region of the genome. The graph shows percentage of plasmid stability and bars represent mean ± SD from three independent replicates.

**Figure S4 (related to Figure 4).**
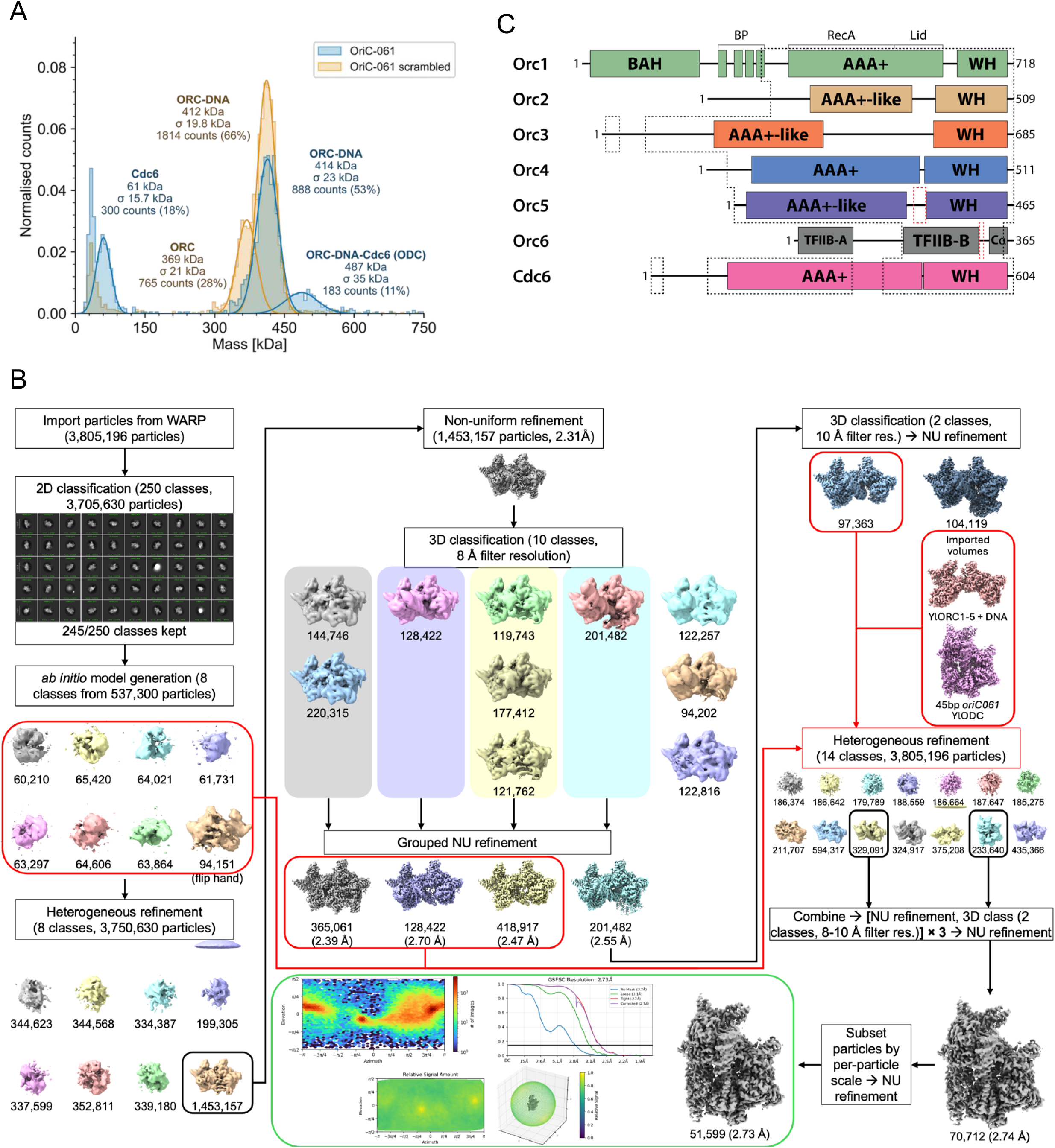
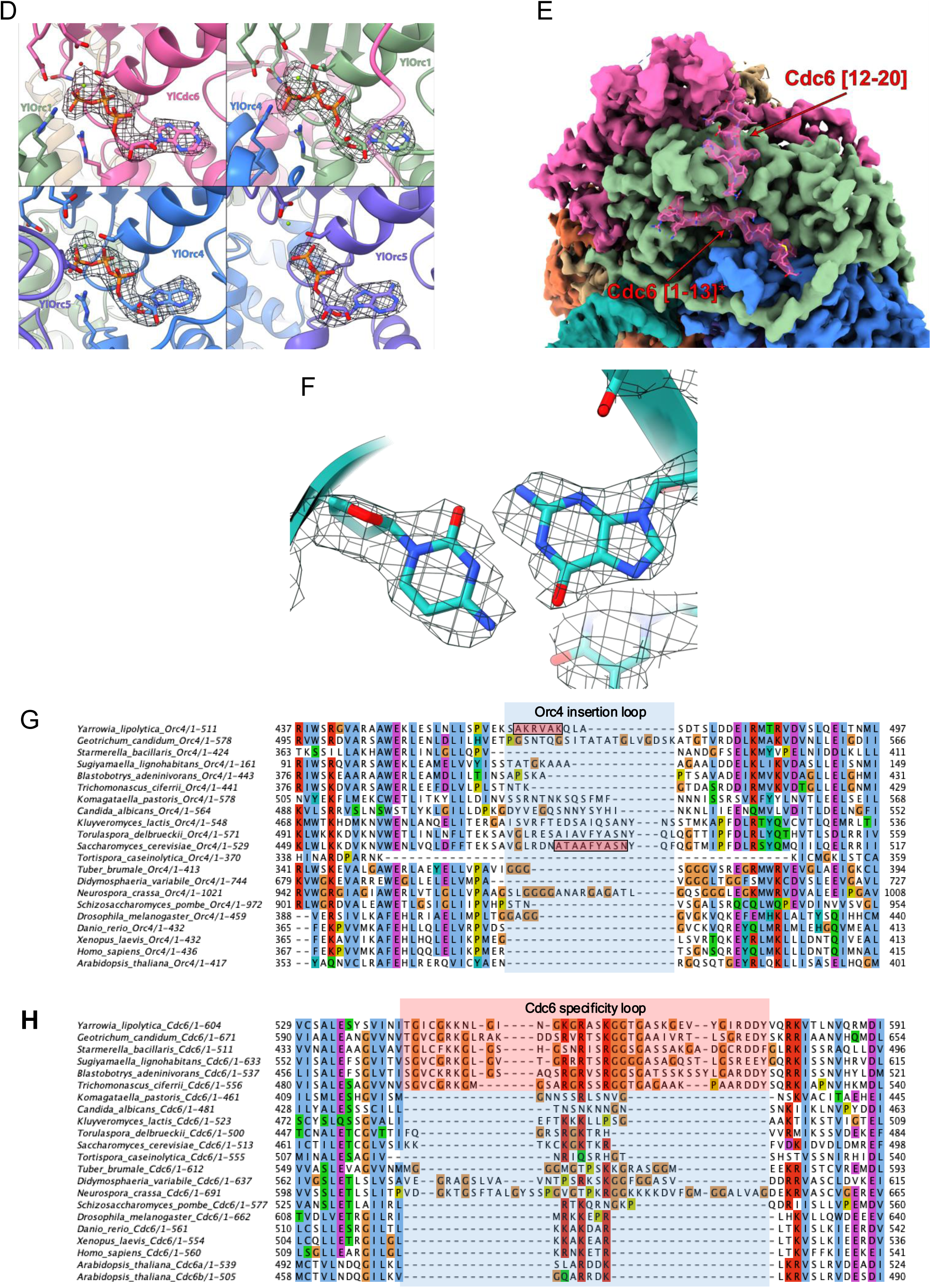
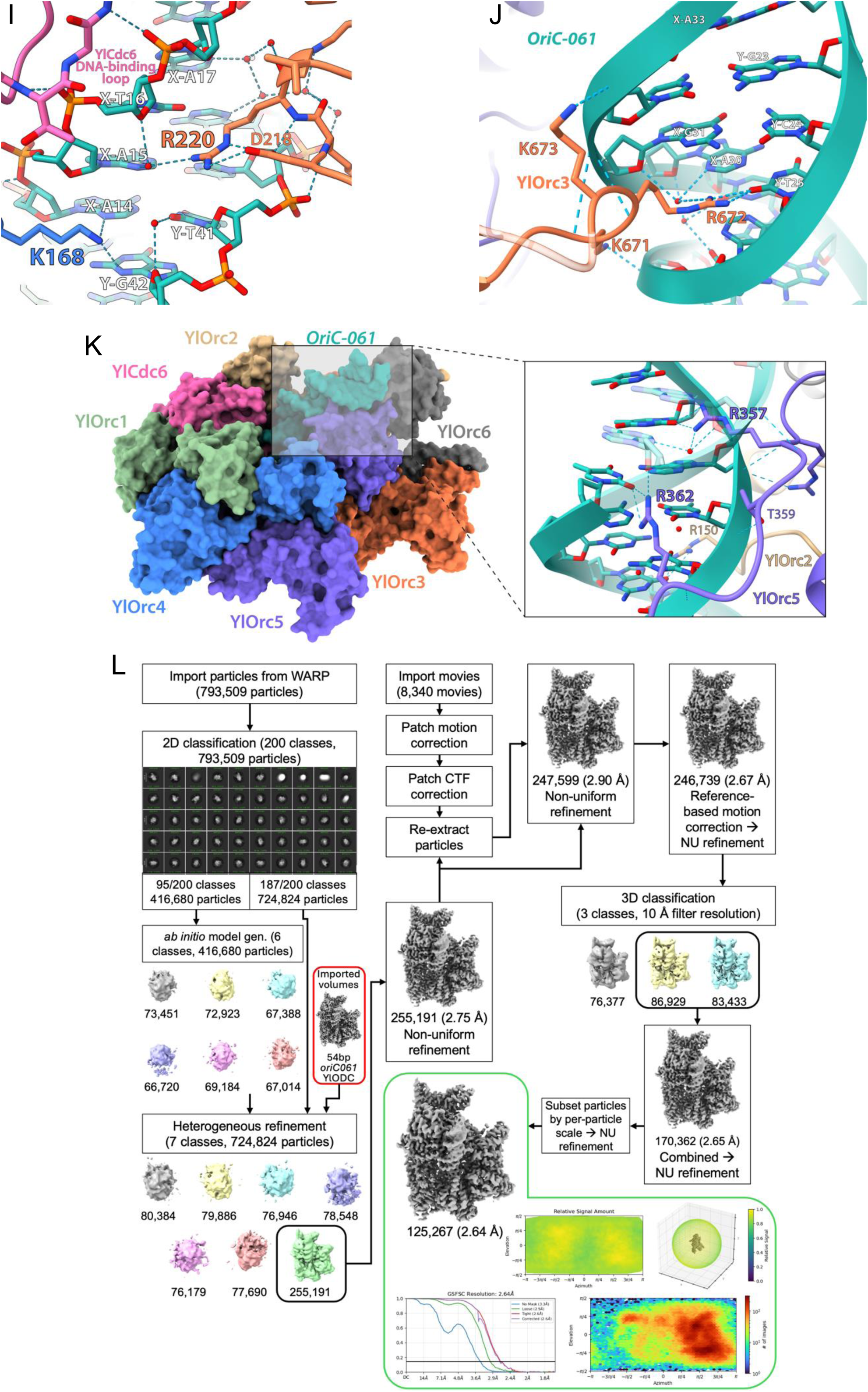
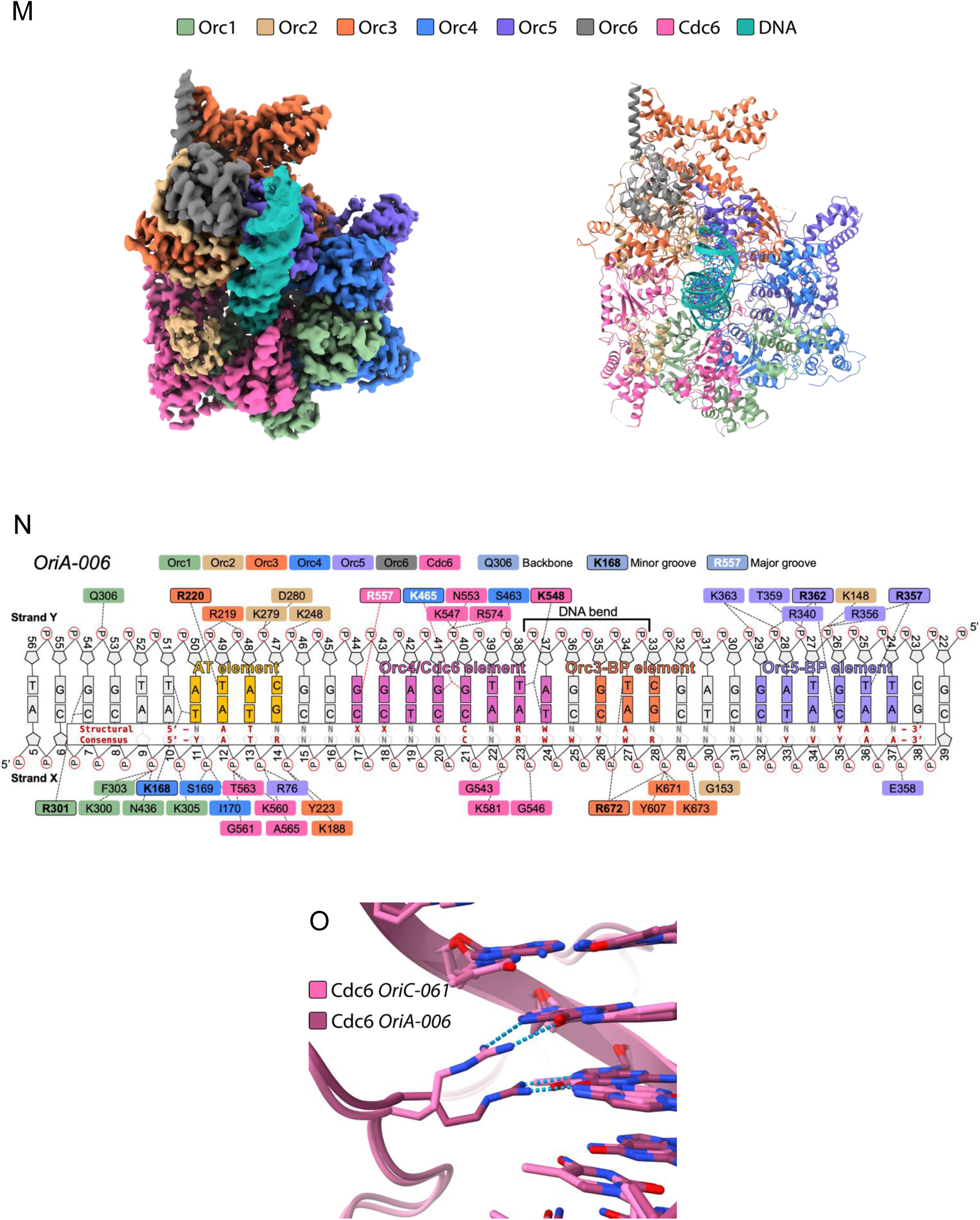
**(A)** Mass photometry data of the gel filtration peak fractions from the loading assay in Figure 4A. **(B)** Schematic diagram of the cryo-EM data processing workflow and statistics from the generation of the YlORC-DNA^54bpOriC-061^–YlCdc6 cryo-EM map. **(C)** Schematic of the domain organization of YlORC and YlCdc6. Dotted lines surround the regions visible within either YlORC-DNA-YlCdc6 maps. **(D)** ADP/ATP in stick representation at the four ATP-binding sites within the YlORC-DNA^54bpOriC-061^–YlCdc6 complex, with a mesh overlay of the representative cryo-EM density of these nucleotides. **(E)** A view of the YlCdc6 IDR sections binding to the YlOrc1•YlOrc4 AAA domain interface and the YlOrc1-AAA•YlCdc6-WHD interface. * denotes the visibility of a leftover glycine on the N-terminus of YlCdc6 resulting from TEV protease cleavage of an N-terminal 8xHis-SUMO tag. **(F)** Representative cryo-EM density of a base pair of *OriC-061* in the YlORC-DNA^54bpOriC-061^–YlCdc6 complex. **(G)** An MSA of the Orc4 region surrounding the Orc4 insertion loop including various related yeasts and eukaryotes. Pink boxes in the YlOrc4 and ScOrc4 sequences denote the region that encompasses the species’ insertion helix. **(H)** An MSA of the Cdc6-WHD surrounding the specificity loop. The red box indicates species containing a conserved specificity loop, while blue indicates those that appear to lack sequence homology in this region. **(I)** View of the AT element (positions X-A14 to X-A17) of *OriC-061* in the YlORC-DNA^54bpOriC-061^–YlCdc6 structure, displaying the hydrogen bond interactions between protein, DNA, and water. **(J)** Same as **(I)** but of the Orc3 basic patch (Orc3-BP) element of *OriC-061.* **(K)** Same as **(I)** but of the Orc5 basic patch (Orc5-BP) element (positions X-G35 to X-A40) of *OriC-061*. **(L)** Schematic diagram of the cryo-EM data processing workflow and statistics from the generation of the YlORC - DNA^60bpOriA-006^–YlCdc6 cryo-EM map. **(M, N)** Same as Figure 4B and 4C, respectively, but of the 2.6 Å resolution cryo-EM map YlORC-DNA^60bpOriA-006^–YlCdc6 and corresponding structure. **(O)** A view of the difference in the major groove base interaction of YlCdc6 R557 between YlORC-DNA^54bpOriC-061^– YlCdc6 and YlORC-DNA^60bpOriA-006^–YlCdc6.

**Figure S5 (related to Figure 5).**
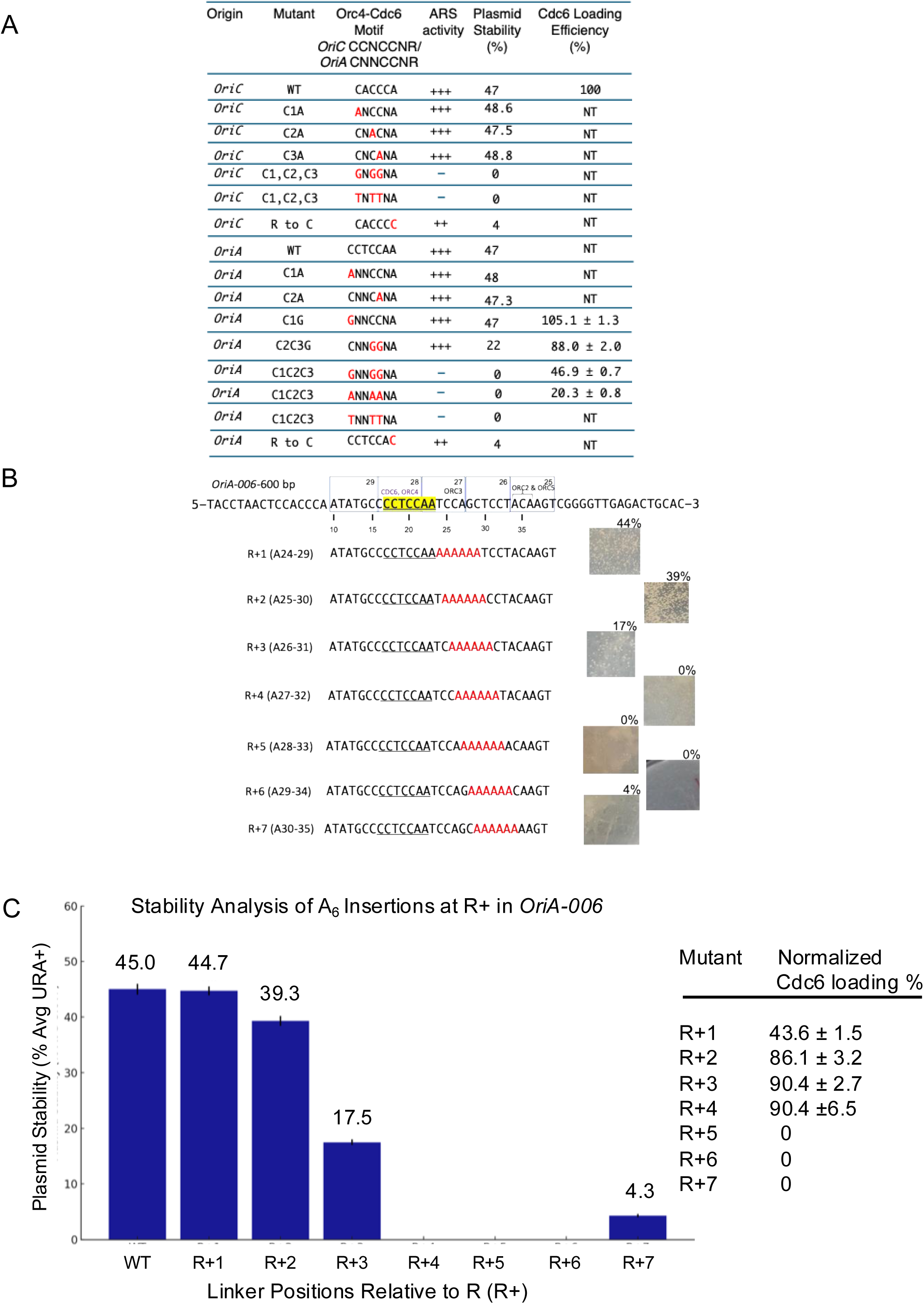
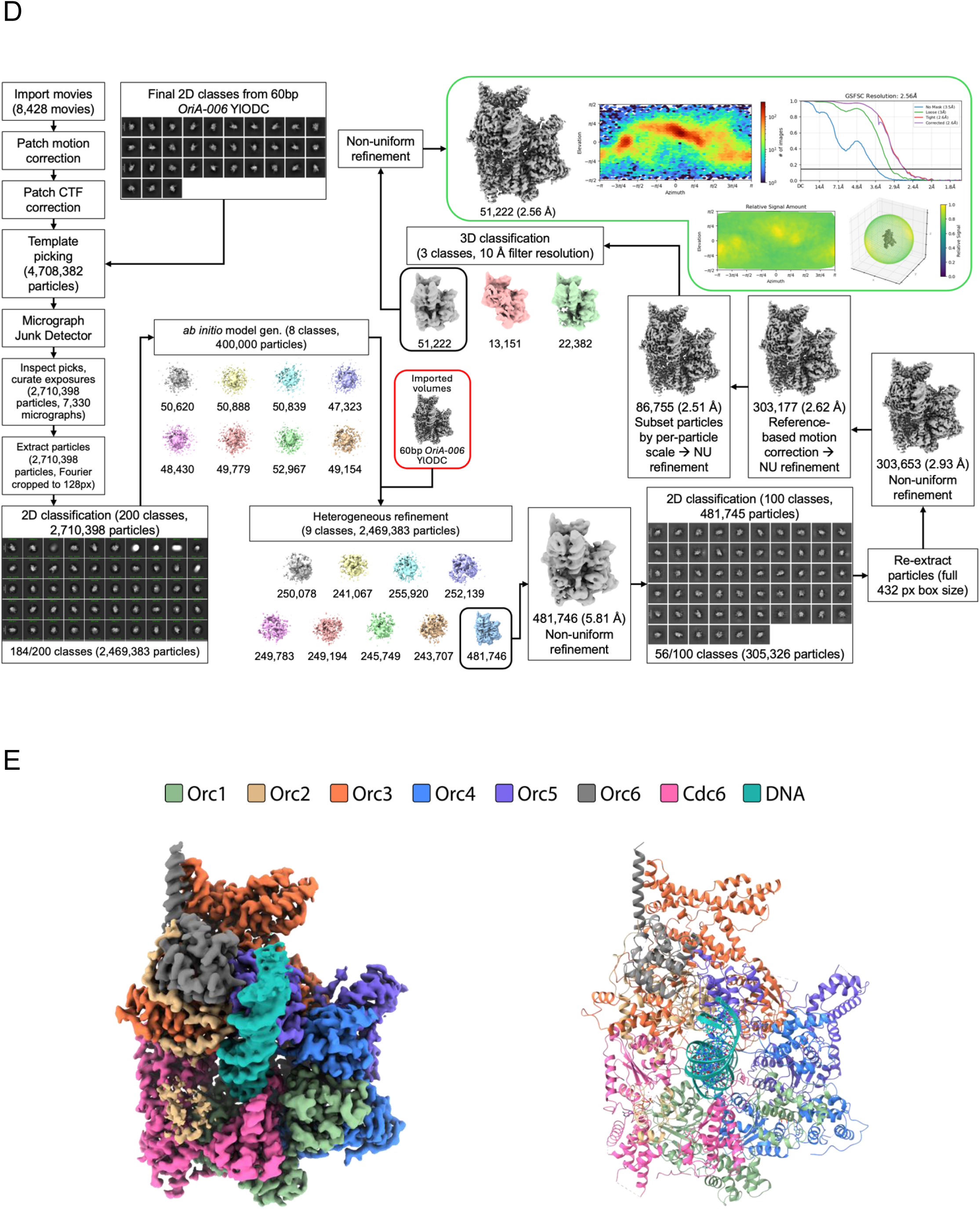
Analysis of *Y. lipolytica* mutant origins. **(A)** Summary table of mutations in *Oria-006* and *OriC-061 reporting* high frequency transformation *(ARS* activity) of point mutations and A_6_ insertions. **(B)** Analysis of mutant *OriA-006* with insertion of six A residues at different positions. The high frequency transformation images and percent plasmid stability was done as described in Figure 5 legend. **(C)** Summary of the plasmid stability of the A_6_ insertion mutations. **(D)** Schematic diagram of the cryo-EM data processing workflow and statistics from the generation of the YlORC-DNA^60bp*OriA-CNNGGNR*^-YlCdc6 cryo-EM map. **(E)** Same as Figure 4B and Figure S4L, but of the 2.6 Å resolution YlORC-DNA^60bpOriA-CNNGGNR^-YlCdc6 cryo-EM map and the accompanying structure.

**Figure S6 (related to Figure 6).**
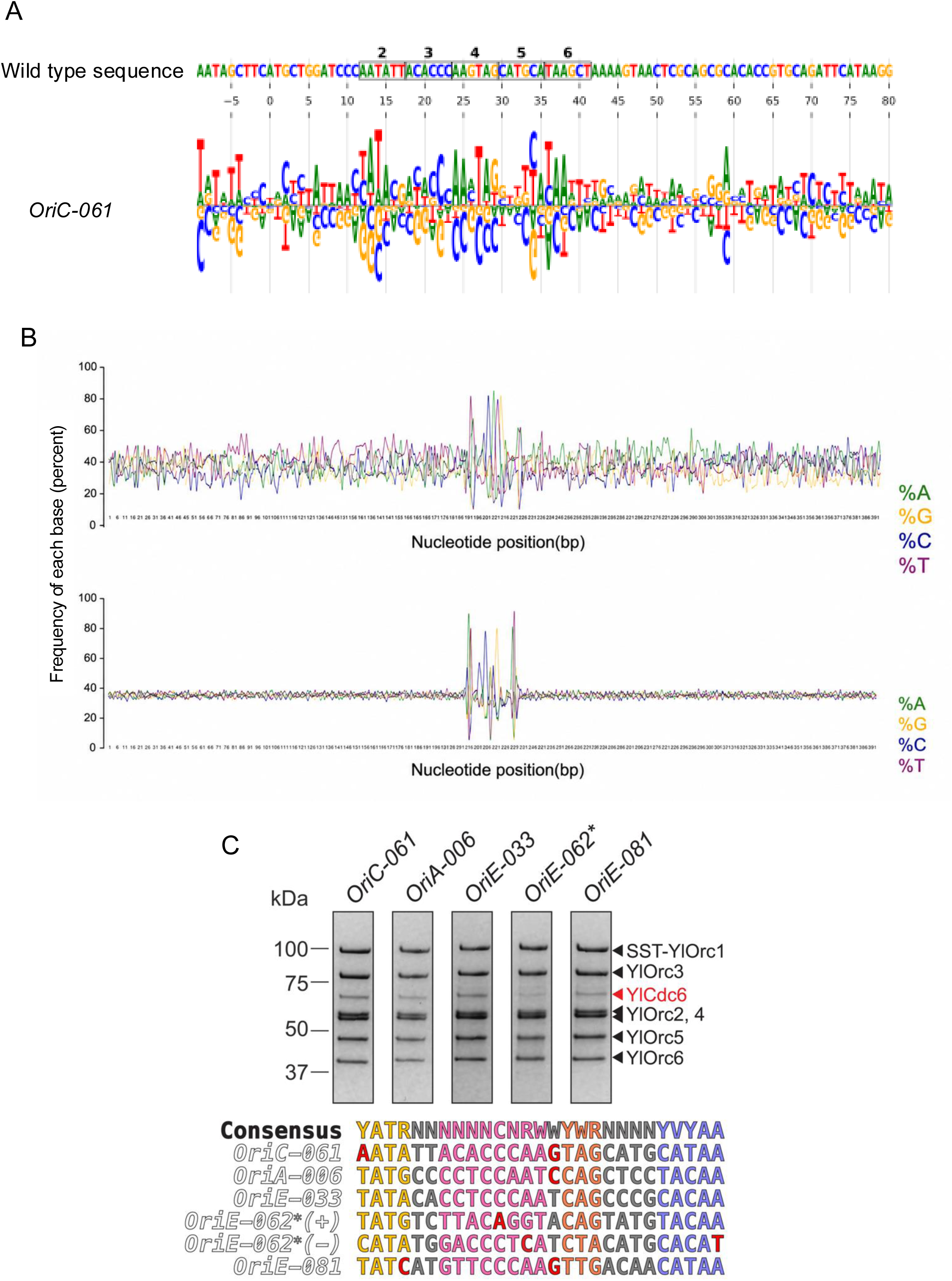
**(A)** Additive model for origin specificity derived from an MPOS assay on 90 bp sequences containing *OriC-061*. Boxed region indicates the core region identified by linker scanning (linker positions 2-6 in Fig. 3A). Note that the base numbers refer to the position within the 90 bp and the numbers refer to the numbering of nucleotides in Figure 4C. **(B).** Analysis of nucleotide frequencies across 54 *Y. lipolytica early* origins of replication compared to 5351 non-origin sequences matched the consensus sequence. The early origins showed a skew for T/A sequences 5’ to the core consensus sequence and a skew for A/T base pairs 3’ to the core consensus sequence. There is no skew surrounding the non-origin consensus sequences. There are 5545 matches to the consensus sequence in the genome. **(C)** Cdc6 loading assays carried out on 60bp dsDNA segments containing *bona fide* origin sequences *OriC-061* (*p* = 2.12e-06) and *OriA-006* (*p* = 3.79e-08) or presumed origin sequences *OriE-033* (*p* = 3.76e-12), *OriE-062\** (contains two matches on opposing strands, + strand *p* = 1.34e-05, - strand *p* = 1.87e-05), and *OriE-081 (p* = 1.19e-04), in addition to each origin’s comparison to the consensus motif, with sequence elements using the previous coloring scheme and mismatches colored red.

## REFERENCES

1. Hu, Y. & Stillman, B. Origins of DNA replication in eukaryotes. Mol. Cell 83, 352–372 (2023).

2. Hyrien, O., Guilbaud, G. & Krude, T. The double life of mammalian DNA replication origins. Genes Dev. 39, 304–324 (2025).

3. Ekundayo, B. & Bleichert, F. Origins of DNA replication. PLoS genetics 15, e1008320 (2019).

4. Lee, C. S. K., Weiβ, M. & Hamperl, S. Where and when to start: Regulating DNA replication origin activity in eukaryotic genomes. Nucleus 14, 2229642 (2023).

5. Costa, A. & Diffley, J. F. X. The Initiation of Eukaryotic DNA Replication. Annu. Rev. Biochem. 91, 107–131 (2022).

6. Bleichert, F., Botchan, M. R. & Berger, J. M. Mechanisms for initiating cellular DNA replication. Science 355, eaah6317 (2017).

7. Bell, S. P. & Labib, K. Chromosome Duplication in Saccharomyces cerevisiae. Genetics 203, 1027–1067 (2016).

8. Musiałek, M. W. & Rybaczek, D. Behavior of replication origins in Eukaryota - spatio-temporal dynamics of licensing and firing. *Cell cycle (Georgetown*, Tex) 0 (2015) doi:10.1080/15384101.2015.1056421.

9. Nishitani, H. & Lygerou, Z. Control of DNA replication licensing in a cell cycle. Genes Cells 7, 523–534 (2002).

10. Rhind, N. DNA replication timing: Biochemical mechanisms and biological significance. Bioessays 2200097 (2022) doi:10.1002/bies.202200097.

11. Limas, J. C. & Cook, J. G. Preparation for DNA replication: the key to a successful S phase. FEBS letters 593, 2853–2867 (2019).

12. Speck, C. & Reuter, L. M. Compact Origins and Where to Find Them: ORC’s Guide to Genome-Wide Licensing. BioEssays e70018 (2025) doi:10.1002/bies.70018.

13. Yeeles, J. T. P., Deegan, T. D., Janska, A., Early, A. & Diffley, J. F. X. Regulated eukaryotic DNA replication origin firing with purified proteins. Nature 519, 431–435 (2015).

14. Kurat, C. F., Yeeles, J. T. P., Patel, H., Early, A. & Diffley, J. F. X. Chromatin Controls DNA Replication Origin Selection, Lagging-Strand Synthesis, and Replication Fork Rates. Mol. Cell 65, 117–130 (2017).

15. Devbhandari, S., Jiang, J., Kumar, C., Whitehouse, I. & Remus, D. Chromatin Constrains the Initiation and Elongation of DNA Replication. Mol. Cell 65, 131–141 (2017).

16. Zhang, Q., Lam, W. H. & Zhai, Y. Assembly and activation of replicative helicases at origin DNA for replication initiation. Curr. Opin. Struct. Biol. 88, 102876 (2024).

17. Stillman, B., Diffley, J. F. X. & Iwasa, J. H. Mechanisms for licensing origins of DNA replication in eukaryotic cells. Nat. Struct. Mol. Biol. 1–11 (2025) doi:10.1038/s41594-025-01587-5.

18. Siow, C. C., Nieduszynska, S. R., Müller, C. A. & Nieduszynski, C. A. OriDB, the DNA replication origin database updated and extended. Nucleic acids research 40, D682–6 (2012).

19. Brewer, B. J. & Fangman, W. L. The localization of replication origins on ARS plasmids in S. cerevisiae. Cell 51, 463–471 (1987).

20. Attali, I., Botchan, M. R. & Berger, J. M. Structural Mechanisms for Replicating DNA in Eukaryotes. Annu. Rev. Biochem. 90, 1–30 (2021).

21. Moyer, S. E., Lewis, P. W. & Botchan, M. R. Isolation of the Cdc45/Mcm2-7/GINS (CMG) complex, a candidate for the eukaryotic DNA replication fork helicase. Proceedings of the National Academy of Sciences of the United States of America 103, 10236–10241 (2006).

22. Marahrens, Y. & Stillman, B. A Yeast Chromosomal Origin of DNA Replication Defined by Multiple Functional Elements. Science 255, 817–823 (1992).

23. Theis, J. F. & Newlon, C. S. Domain B of ARS307 contains two functional elements and contributes to chromosomal replication origin function. Mol Cell Biol 14, 7652–7659 (1994).

24. Rao, H., Marahrens, Y. & Stillman, B. Functional Conservation of Multiple Elements in Yeast Chromosomal Replicators. Mol. Cell. Biol. 14, 7643–7651 (1994).

25. Reuter, L. M. et al. MCM2-7 loading-dependent ORC release ensures genome-wide origin licensing. Nat. Commun. 15, 7306 (2024).

26. Coster, G. & Diffley, J. F. X. Bidirectional eukaryotic DNA replication is established by quasi-symmetrical helicase loading. Science 357, 314–318 (2017).

27. Gupta, S., Friedman, L. J., Gelles, J. & Bell, S. P. A helicase-tethered ORC flip enables bidirectional helicase loading. Elife 10, e74282 (2021).

28. Hu, Y. et al. Evolution of DNA replication origin specification and gene silencing mechanisms. Nat. Commun. 11, 5175 (2020).

29. Wang, D. & Gao, F. Comprehensive Analysis of Replication Origins in Saccharomyces cerevisiae Genomes. Frontiers in Microbiology 10, 2122 (2019).

30. Yuan, Z. et al. Structural basis of Mcm2–7 replicative helicase loading by ORC–Cdc6 and Cdt1. Nat. Struct. Mol. Biol. 24, 316–324 (2017).

31. Li, N. et al. Structure of the origin recognition complex bound to DNA replication origin. Nature 559, 217–222 (2018).

32. Lee, C. S. K. et al. Humanizing the yeast origin recognition complex. Nat. Commun. 12, 33 (2021).

33. Liachko, I. et al. A comprehensive genome-wide map of autonomously replicating sequences in a naive genome. PLoS genetics 6, e1000946 (2010).

34. Liachko, I. et al. Novel features of ARS selection in budding yeast Lachancea kluyveri. BMC Genom. 12, 633 (2011).

35. Hu, Y. et al. Evolution of DNA replication origin specification and gene silencing mechanisms. Nat. Commun. 11, 5175 (2020).

36. Tsai, H.-J. et al. Origin replication complex binding, nucleosome depletion patterns, and a primary sequence motif can predict origins of replication in a genome with epigenetic centromeres. Mbio 5, e01703–14 (2014).

37. Liachko, I. et al. GC-rich DNA elements enable replication origin activity in the methylotrophic yeast Pichia pastoris. PLoS genetics 10, e1004169 (2014).

38. Evrin, C. et al. A double-hexameric MCM2-7 complex is loaded onto origin DNA during licensing of eukaryotic DNA replication. Proc. Natl. Acad. Sci. 106, 20240–20245 (2009).

39. Remus, D. et al. Concerted Loading of Mcm2–7 Double Hexamers around DNA during DNA Replication Origin Licensing. Cell 139, 719–730 (2009).

40. Weissmann, F. et al. MCM double hexamer loading visualized with human proteins. Nature 636, 499–508 (2024).

41. Yang, R., Hunker, O., Wise, M. & Bleichert, F. Multiple mechanisms for licensing human replication origins. Nature 636, 488–498 (2024).

42. Wells, J. N. et al. Reconstitution of human DNA licensing and the structural and functional analysis of key intermediates. Nat. Commun. 16, 478 (2025).

43. Tian, M. et al. Integrative analysis of DNA replication origins and ORC-/MCM-binding sites in human cells reveals a lack of overlap. eLife 12, RP89548 (2024).

44. Ganier, O., Prorok, P., Akerman, I. & Méchali, M. Metazoan DNA replication origins. Current opinion in cell biology 58, 134–141 (2019).

45. Wang, W. et al. Genome-wide mapping of human DNA replication by optical replication mapping supports a stochastic model of eukaryotic replication. Mol Cell (2021) doi:10.1016/j.molcel.2021.05.024.

46. Pourkarimi, E., Bellush, J. M. & Whitehouse, I. Spatiotemporal coupling and decoupling of gene transcription with DNA replication origins during embryogenesis in C. elegans. Elife 5, e21728 (2016).

47. Comoglio, F. et al. High-Resolution Profiling of Drosophila Replication Start Sites Reveals a DNA Shape and Chromatin Signature of Metazoan Origins. Cell Reports 11, 821–834 (2015).

48. Masai, H. & Tanaka, T. G-quadruplex DNA and RNA: Their roles in regulation of DNA replication and other biological functions. Biochem. Biophys. Res. Commun. 531, 25–38 (2020).

49. Valton, A.-L. & Prioleau, M.-N. G-Quadruplexes in DNA Replication: A Problem or a Necessity? Trends in Genetics 32, 697–706 (2016).

50. Valton, A. L. et al. G4 motifs affect origin positioning and efficiency in two vertebrate replicators. The EMBO journal 33, 732–746 (2014).

51. Prorok, P. et al. Involvement of G-quadruplex regions in mammalian replication origin activity. Nat Commun 10, 3274 (2019).

52. Bartlett, D. A., Dileep, V., Baslan, T. & Gilbert, D. M. Mapping Replication Timing in Single Mammalian Cells. Curr Protoc 2, e334 (2022).

53. Carrington, J. T. et al. Most human DNA replication initiation is dispersed throughout the genome with only a minority within previously identified initiation zones. Genome Biol. 26, 122 (2025).

54. Akerman, I. et al. A predictable conserved DNA base composition signature defines human core DNA replication origins. Nat Commun 11, 4826 (2020).

55. Guilbaud, G. et al. Determination of human DNA replication origin position and efficiency reveals principles of initiation zone organisation. Nucleic Acids Res (2022) doi:10.1093/nar/gkac555.

56. Langley, A. R., Gräf, S., Smith, J. C. & Krude, T. Genome-wide identification and characterisation of human DNA replication origins by initiation site sequencing (ini-seq). Nucleic acids research 44, 10230–10247 (2016).

57. Petryk, N. et al. Replication landscape of the human genome. Nature Communications 7, 10208 (2016).

58. Krude, T., Bi, J., Doran, R., Jones, R. A. & Smith, J. C. Human DNA replication initiation sites are specified epigenetically by oxidation of 5-methyl-deoxycytidine. Nucleic Acids Res. 53, gkaf362 (2025).

59. Méchali, M., Yoshida, K., Coulombe, P. & Pasero, P. Genetic and epigenetic determinants of DNA replication origins, position and activation. Curr. Opin. Genet. Dev. 23, 124–131 (2013).

60. Srikant, S., Gaudet, R. & Murray, A. W. Extending the reach of homology by using successive computational filters to find yeast pheromone genes. Curr. Biol. (2023) doi:10.1016/j.cub.2023.08.039.

61. Vernis, L. et al. An Origin of Replication and a Centromere Are Both Needed To Establish a Replicative Plasmid in the Yeast Yarrowia lipolytica. Mol. Cell. Biol. 17, 1995–2004 (1997).

62. Vernis, L. et al. Short DNA Fragments without Sequence Similarity Are Initiation Sites for Replication in the Chromosome of the Yeast Yarrowia lipolytica. Mol. Biol. Cell 10, 757–769 (1999).

63. Vernis, L. et al. Only Centromeres Can Supply the Partition System Required for ARS Function in the Yeast Yarrowia lipolytica. J. Mol. Biol. 305, 203–217 (2001).

64. Lopez, C., Cao, M., Yao, Z. & Shao, Z. Investigating the role of noncoding regulatory DNA in plasmid development for Yarrowia lipolytica. doi:10.22541/au.160691063.38058320/v1.

65. Prioleau, M.-N. & MacAlpine, D. M. DNA replication origins—where do we begin? Genes Dev. 30, 1683–1697 (2016).

66. Jaremko, M. J., On, K. F., Thomas, D. R., Stillman, B. & Joshua-Tor, L. The dynamic nature of the human origin recognition complex revealed through five cryoEM structures. eLife 9, e58622 (2020).

67. Tocilj, A. et al. Structure of the active form of human Origin Recognition Complex and its ATPase motor module. (2017) doi:10.2210/pdb5ujm/pdb.

68. Schmidt, J. M. et al. A mechanism of origin licensing control through autoinhibition of S. cerevisiae ORC·DNA·Cdc6. Nat. Commun. 13, 1059 (2022).

69. Schmidt, J. M. & Bleichert, F. Structural mechanism for replication origin binding and remodeling by a metazoan origin recognition complex and its co-loader Cdc6. Nat Commun 11, 4263 (2020).

70. Feng, X. et al. The structure of ORC–Cdc6 on an origin DNA reveals the mechanism of ORC activation by the replication initiator Cdc6. Nat. Commun. 12, 3883 (2021).

71. Lam, W. H. et al. DNA bending mediated by ORC is essential for replication licensing in budding yeast. Proc. Natl. Acad. Sci. 122, e2502277122 (2025).

72. Erzberger, J. P. & Berger, J. M. Evolutionary Relationships and Structural Mechanism of AAA+ Proteins. Annu. Rev. Biophys. Biomol. Struct. 35, 93–114 (2006).

73. Zali, N. et al. Genome sequence assembly and annotation of MATA and MATB strains of Yarrowia lipolytica. NAR Genom. Bioinform. 7, lqaf175 (2025).

74. Fu, H., Baris, A. & Aladjem, M. I. Replication timing and nuclear structure. Curr. Opin. Cell Biol. 52, 43–50 (2018).

75. Marchal, C., Sima, J. & Gilbert, D. M. Control of DNA replication timing in the 3D genome. Nat. Rev. Mol. Cell Biol. 20, 721–737 (2019).

76. McKnight, S. L. & Kingsbury, R. Transcriptional Control Signals of a Eukaryotic Protein-Coding Gene. Science 217, 316–324 (1982).

77. Nelson, H. C. M., Finch, J. T., Luisi, B. F. & Klug, A. The structure of an oligo(dA)·oligo(dT) tract and its biological implications. Nature 330, 221–226 (1987).

78. Haran, T. E., Kahn, J. D. & Crothers, D. M. Sequence Elements Responsible for DNA Curvature. J. Mol. Biol. 244, 135–143 (1994).

79. Remus, D., Beall, E. L. & Botchan, M. R. DNA topology, not DNA sequence, is a critical determinant for Drosophila ORC–DNA binding. EMBO J. 23, 897–907 (2004).

80. Tareen, A. et al. MAVE-NN: learning genotype-phenotype maps from multiplex assays of variant effect. Genome Biol. 23, 98 (2022).

81. Oji, A., Choubani, L., Miura, H. & Hiratani, I. Structure and dynamics of nuclear A/B compartments and subcompartments. Curr. Opin. Cell Biol. 90, 102406 (2024).

82. Vouzas, A. E. & Gilbert, D. M. Replication timing and transcriptional control: beyond cause and effect — part IV. Curr Opin Genet Dev 79, 102031 (2023).

83. McCune, H. J. et al. The Temporal Program of Chromosome Replication: Genomewide Replication in clb5Δ Saccharomyces cerevisiae. Genetics 180, 1833–1847 (2008).

84. Chang, F. et al. High-resolution analysis of four efficient yeast replication origins reveals new insights into the ORC and putative MCM binding elements. Nucleic Acids Res. 39, 6523–6535 (2011).

85. Bell, S. P. & Stillman, B. ATP-dependent recognition of eukaryotic origins of DNA replication by a multiprotein complex. Nature 357, 128–134 (1992).

86. Bleichert, F., Leitner, A., Aebersold, R., Botchan, M. R. & Berger, J. M. Conformational control and DNA-binding mechanism of the metazoan origin recognition complex. Proceedings of the National Academy of Sciences 115, E5906–E5915 (2018).

87. Miller, T. C. R., Locke, J., Greiwe, J. F., Diffley, J. F. X. & Costa, A. Mechanism of head-to-head MCM double-hexamer formation revealed by cryo-EM. Nature 575, 704–710 (2019).

88. Yuan, Z. et al. Structural mechanism of helicase loading onto replication origin DNA by ORC-Cdc6. Proc. Natl. Acad. Sci. 117, 17747–17756 (2020).

89. Dahlin, J., Holkenbrink, C. & Borodina, I. Yarrowia lipolytica, Methods and Protocols. in vol. 2307 41–68 (2021).

90. Lopez, C., Cao, M., Yao, Z. & Shao, Z. Revisiting the unique structure of autonomously replicating sequences in Yarrowia lipolytica and its role in pathway engineering. Appl Microbiol Biot 1–14 (2021) doi:10.1007/s00253-021-11399-4.

91. Chen, S., Zhou, Y., Chen, Y. & Gu, J. fastp: an ultra-fast all-in-one FASTQ preprocessor. Bioinformatics 34, i884–i890 (2018).

92. Langmead, B. & Salzberg, S. L. Fast gapped-read alignment with Bowtie 2. Nat. Methods 9, 357–359 (2012).

93. Danecek, P. et al. Twelve years of SAMtools and BCFtools. GigaScience 10, giab008 (2021).

94. Ramírez, F. et al. deepTools2: a next generation web server for deep-sequencing data analysis. Nucleic Acids Res. 44, W160–W165 (2016).

95. Zhang, Y. et al. Model-based Analysis of ChIP-Seq (MACS). Genome Biol. 9, R137 (2008).

96. Heinz, S. et al. Simple Combinations of Lineage-Determining Transcription Factors Prime cis-Regulatory Elements Required for Macrophage and B Cell Identities. Mol. Cell 38, 576–589 (2010).

97. Kinney, J. B., Murugan, A., Callan, C. G. & Cox, E. C. Using deep sequencing to characterize the biophysical mechanism of a transcriptional regulatory sequence. Proceedings of the National Academy of Sciences 107, 9158–9163 (2010).

98. Tareen, A. & Kinney, J. B. Logomaker: beautiful sequence logos in Python. Bioinformatics 36, 2272–2274 (2020).

99. Bieniossek, C., Richmond, T. J. & Berger, I. MultiBac: Multigene Baculovirus-Based Eukaryotic Protein Complex Production. Curr. Protoc. Protein Sci. 51, 5.20.1–5.20.26 (2008).

100. Schindelin, J., et al. Fiji: an open-source platform for biological-image analysis. Nat. Methods 9, 676–682 (2012).

101. Tegunov, D. & Cramer, P. Real-time cryo-electron microscopy data preprocessing with Warp. Nat. Methods 16, 1146–1152 (2019).

102. Zivanov, J., Nakane, T. & Scheres, S. H. W. Estimation of high-order aberrations and anisotropic magnification from cryo-EM data sets in RELION-3.1. IUCrJ 7, 253–267 (2020).

103. Punjani, A., Rubinstein, J. L., Fleet, D. J. & Brubaker, M. A. cryoSPARC: algorithms for rapid unsupervised cryo-EM structure determination. Nat. Methods 14, 290–296 (2017).

104. Punjani, A., Zhang, H. & Fleet, D. J. Non-uniform refinement: adaptive regularization improves single-particle cryo-EM reconstruction. Nat. Methods 17, 1214–1221 (2020).

105. Goddard, T. D. et al. UCSF ChimeraX: Meeting modern challenges in visualization and analysis. Protein science : a publication of the Protein Society 27, 14–25 (2018).

106. Emsley, P., Lohkamp, B., Scott, W. G. & Cowtan, K. Features and development of Coot. Acta Crystallogr. Sect. D 66, 486–501 (2010).

107. Liebschner, D. et al. Macromolecular structure determination using X-rays, neutrons and electrons: recent developments in Phenix. Acta Crystallogr. Sect. D 75, 861–877 (2019).

