## Supplemental Videos 1 and 2 for "Evolution of Origin Sequence and Recognition for Licensing of Eukaryotic DNA Replication"

### Slide 1
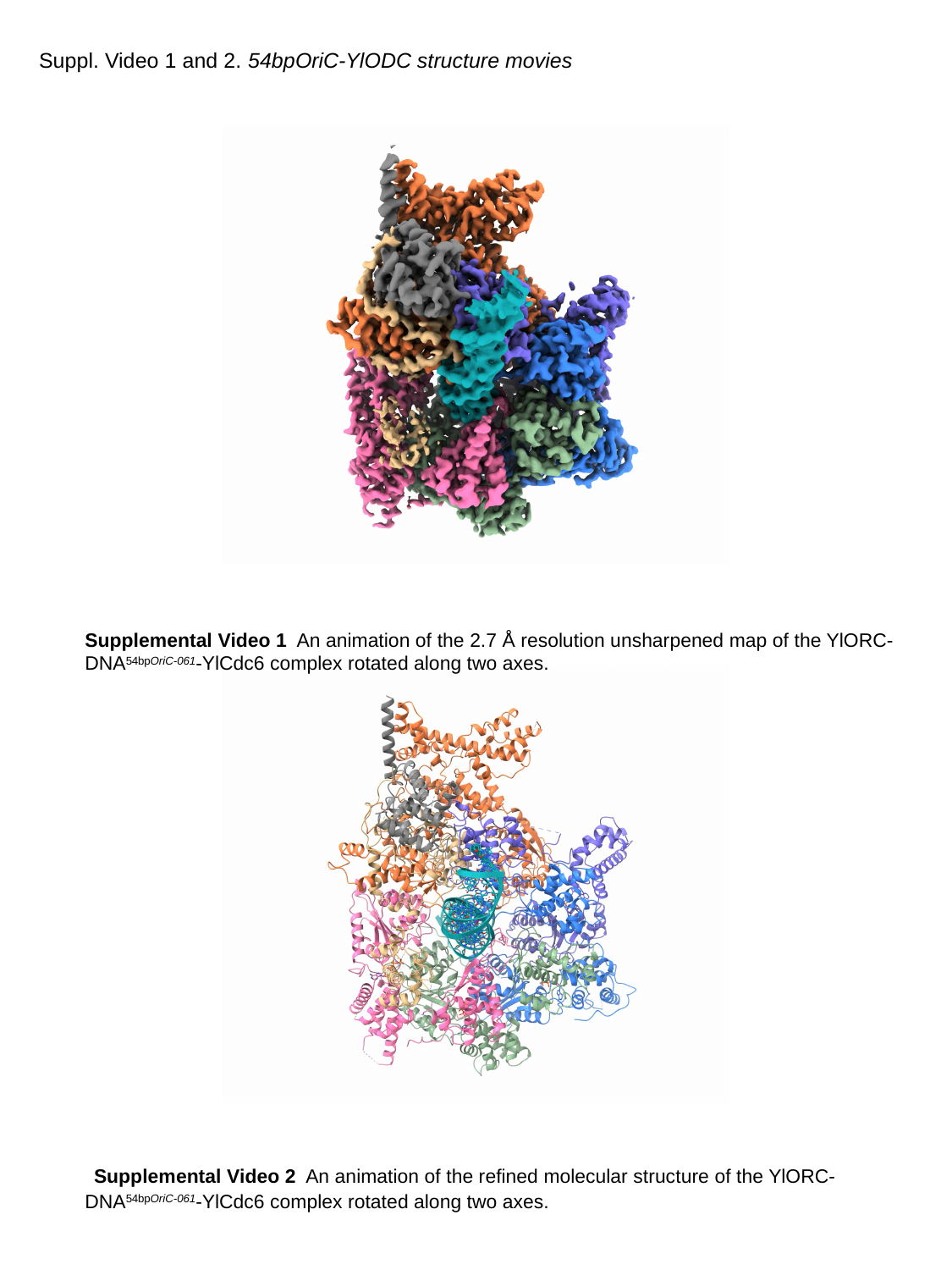

#### Suppl. Video 1 and 2. 54bpOriC-YlODC structure movies
Supplemental Video 1 An animation of the 2.7 Å resolution unsharpened map of the YlORC-DNA^54bpOriC-061^-YlCdc6 complex rotated along two axes.
 Supplemental Video 2 An animation of the refined molecular structure of the YlORC-DNA^54bpOriC-061^-YlCdc6 complex rotated along two axes.
