## Supplemental Video 3 for "Evolution of Origin Sequence and Recognition for Licensing of Eukaryotic DNA Replication"

### Slide 1
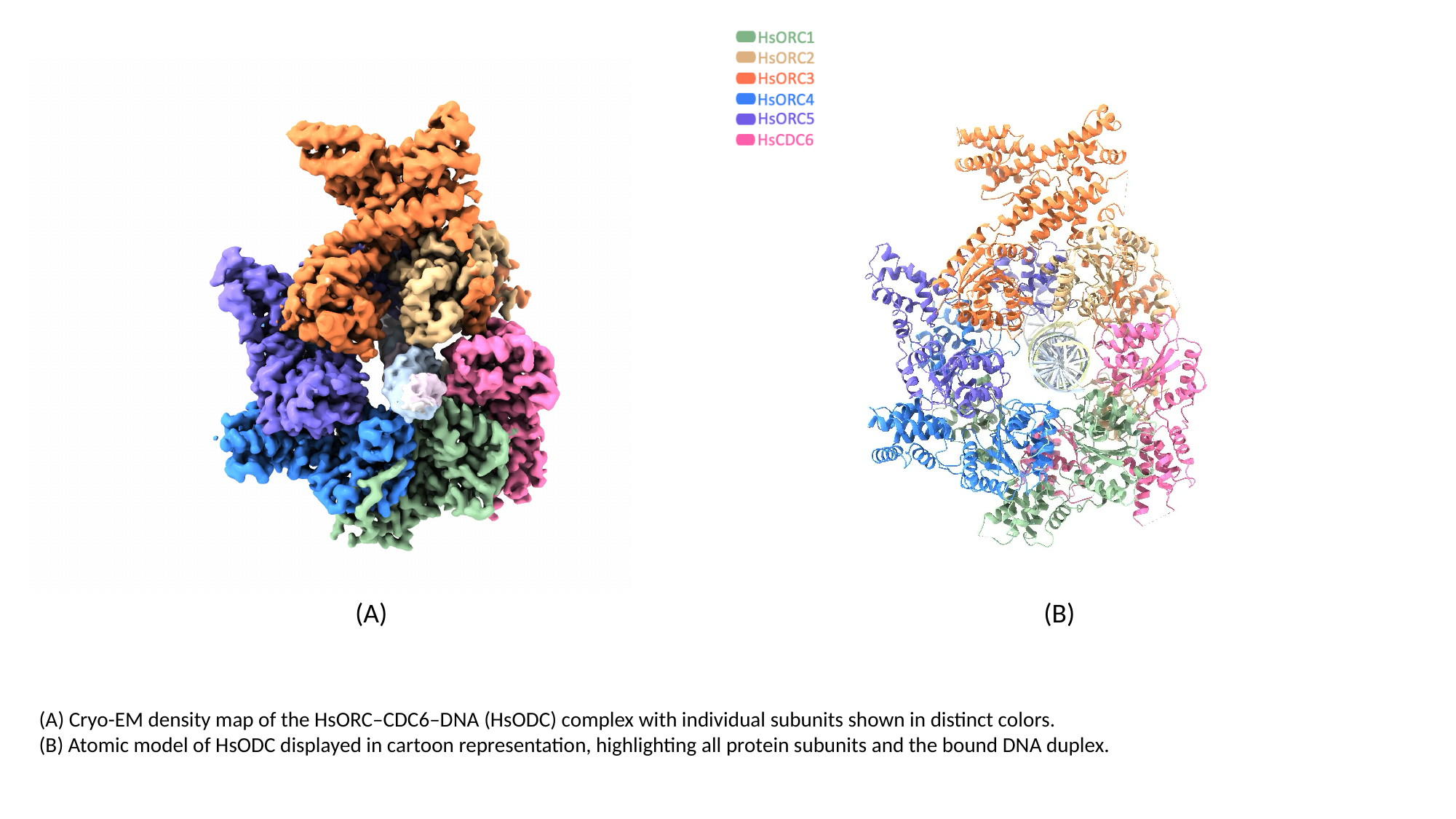

(A)
(B)
(A) Cryo-EM density map of the HsORC–CDC6–DNA (HsODC) complex with individual subunits shown in distinct colors.
(B) Atomic model of HsODC displayed in cartoon representation, highlighting all protein subunits and the bound DNA duplex.
